# Organic matter degradation in the deep, sulfidic waters of the Black Sea: Insights into the ecophysiology of novel anaerobic bacteria

**DOI:** 10.1101/2023.09.28.559987

**Authors:** Subhash Yadav, Michel Koenen, Nicole J Bale, Wietse Reitsma, Julia C Engelmann, Jaap S. Sinninghe Damsté, Laura Villanueva

## Abstract

Our knowledge about the physiology of deep sea (>1,000 m) microorganisms involved in organic matter (OM) degradation is still scare due to the lack of available isolates, especially from sulfidic environments. In this study, we successfully cultivated and characterized the physiology of a wide range of novel piezotolerant anaerobic bacteria affiliated with the phyla *Fusobacteriota*, *Bacillota*, *Spirochaetota*, *Bacteroidota*, *Cloacimonadota*, *Planctomycetota*, *Mycoplasmatota* and *Chloroflexota* involved in OM degradation in deep sulfidic waters of the Black Sea. The novel taxa are specialized in degrading specific types of OM and cover a wide range of physiological categories, including primary degraders, fermenters, and terminal oxidizers. This is the first report which demonstrates this for such a diverse physiological group from any sulfidic marine habitat. Collectively, this study provides a step forward in our understanding of the microbes thriving in the extreme conditions of the deep sulfidic waters of the Black Sea.

## Introduction

Mineralization of organic matter (OM) in the world’s oceans is crucial for global carbon cycling^1^. To understand these processes, it is important to identify the microorganisms involved. Recent advances in DNA sequencing and stable isotope labeling techniques have revealed valuable information about the identities of microorganisms involved in OM degradation in deep marine ecosystems^2–5^. However, to understand their physiology and specific role in these processes, it is essential to isolate these microorganisms. Unfortunately, cultivating deep sea microorganisms presents significant challenges due their adaptation to oligotrophic environments, low temperatures, and limited oxygen concentrations.

Recent studies have identified several key microbial taxa that play a role in the degradation of OM in deep sulfidic marine habitats^4–9^. For instance, *Psychrilyobacter* spp. have been established as important degrader of cellulose and the protein fraction of OM sinking into these habitats. Additionally, uncultivated clades of the order *Clostridiales* (*Fusibacteraceae* ASV_09916) and *Lutibacter* spp. have been observed as significant OM degraders^4,6,7^, while members of the phylum *Spirochaetota* are frequently involved in OM degradation but are often ignored due to their low abundance in incubation-based studies^4,5,7^. Members of the phylum *Mycoplasmatota* are also present in deep marine habitats and have been identified as specialized degraders of DNA^5,8–10^. In addition to these key taxa, there are also other microbial members affiliated with the phyla *Bacteroidota*(especially *Marinifilaceae*, *Marinilabiaceae*, *Lentimicrobiaceae*), *Desulfobacterota* (especially *Desulfovibrionaceae*, *Desulfobacteraceae*, *Desulfobulbaceae*), *Planctomycetota* (*Phycisphaerae*), *Ignavibacteriota*, *Chloroflexota* (*Anaerolineae*), *Cloacimonadota*, and *Nanoarchaeota,* which have been predicted to be involved in OM degradation in deep marine habitats^4,5,7,11–15^. Although the identity and potential functions of these microbial taxa are known, their physiology remains largely unknown due to the limited availability of isolated representatives and enrichment cultures.

The Black Sea is a unique ecosystem, offering unique opportunities to study microbial communities involved in the degradation of OM in sulfidic marine habitats. Its deep waters are permanently anoxic, containing high sulfide concentrations of 400 µM^16^. Additionally, the concentration of dissolved organic carbon (DOC) in the Black Sea is about 2.5 times higher than in any of the major ocean basins^17–19^. Recent studies by Suominen et al.^13–15^ focused on the role of uncultivated microbial communities in OM degradation in the deep sulfidic waters of the Black Sea, concluding that members of the *Gammaproteobacteria*, *Alphaproteobacteria*, *Bacteroidota*, and *Planctomycetota* thrive in incubations with complex carbon sources, such as chitin and peptidoglycan, while members of the phyla *Planctomycetota*, *Latescibacteria*, *Kirimatillaeota*, *Cloacimonadota*, *Nanoarchaeota*, and *Chloroflexota* also do so in incubations with fatty acids^13,14^. However, detailed investigations of the physiology of these microbial groups and their adaptation strategy are still lacking.

Here, we report the enrichment and isolation of novel anaerobic bacteria potentially involved in OM degradation at elevated hydrostatic pressure (20 MPa), high sulfide concentration (>1mM), and relatively low temperature (10°C). To elucidate the ecophysiology of novel isolates and enrichment cultures, we investigated their genomes, determined their membrane lipid composition, and tested the physiology and adaptation strategies by mimicking the *in*-*situ* physicochemical conditions. This study represents a significant progress in the culturing and ecophysiology of novel anaerobic microorganisms that degrade OM in deep sulfidic marine environments.

## 2. Results and Discussion

The microbial community composition in the sulfidic waters of the Black Sea is mostly comprised of uncultivated microbial taxa ubiquitously occurring in anoxic environments worldwide, like *Fusobacteriota*, *Cloacimonadota*, *Planctomycetota*, *Chloroflexota*, *Desulfobacterota*, *Bacteroidota*, and *Omnitrophica*^6,11–15,20–25^. While some uncultivated clades of the order *Clostridiales*, *Spirochaetota*, *Fusobacteriota*, and *Bacteroidota* occur in relatively low natural abundance (∼1% of the total microbial community), they rapidly respond to OM-rich conditions and become abundant^4,6,13,14^. In this study, we utilized various growth media containing different types of OM (yeast extract, tryptone, and various other carbon and nitrogen sources) and sulfide/sulfidic waters to enrich and isolate some of these microorganisms (Figs. 1A-B, 2-3; see M&M for further details). We successfully isolated pure cultures affiliated with the following phyla: *Fusobacteriota* (*Psychrilyobacter piezotolerans* strain S5), *Bacillota* (*Clostridiales* bacteria strains A1^T^ and A2), *Spirochaetota* (*Oceanispirochaeta* sp. strains M1^T^ and M2; *Sphaerochaeta* sp. strain S2), *Bacteroidota* (*Ancylomarina* sp. strain M2P; *Labilibaculum* sp. strain SYP; *Lentimicrobiales* bacteria strains S6, L6; *Lutibacter* sp. strains B1^T^, and B2), and *Desulfobacterota* (*Pseudodesulfovibrio* sp. strains S3^T^ and S3-i; Fig. S1-S6). Additionally, we obtained enrichment cultures of species belonging to *Planctomycetota* (Plnct-SY6), *Cloacimonadota* (Cloa-SY6), *Ignavibacteriota* (Igna-SY6), *Chloroflexota* (Chflx-SY6), and *Mycoplasmatota* (Izemo-BS; Fig. S7-S11). However, we did not achieve to isolate them, possibly due to their strict syntrophic nature ^21,26^ or their dependence on unknown growth factors produced under dynamic conditions.

**Fig. 1:**
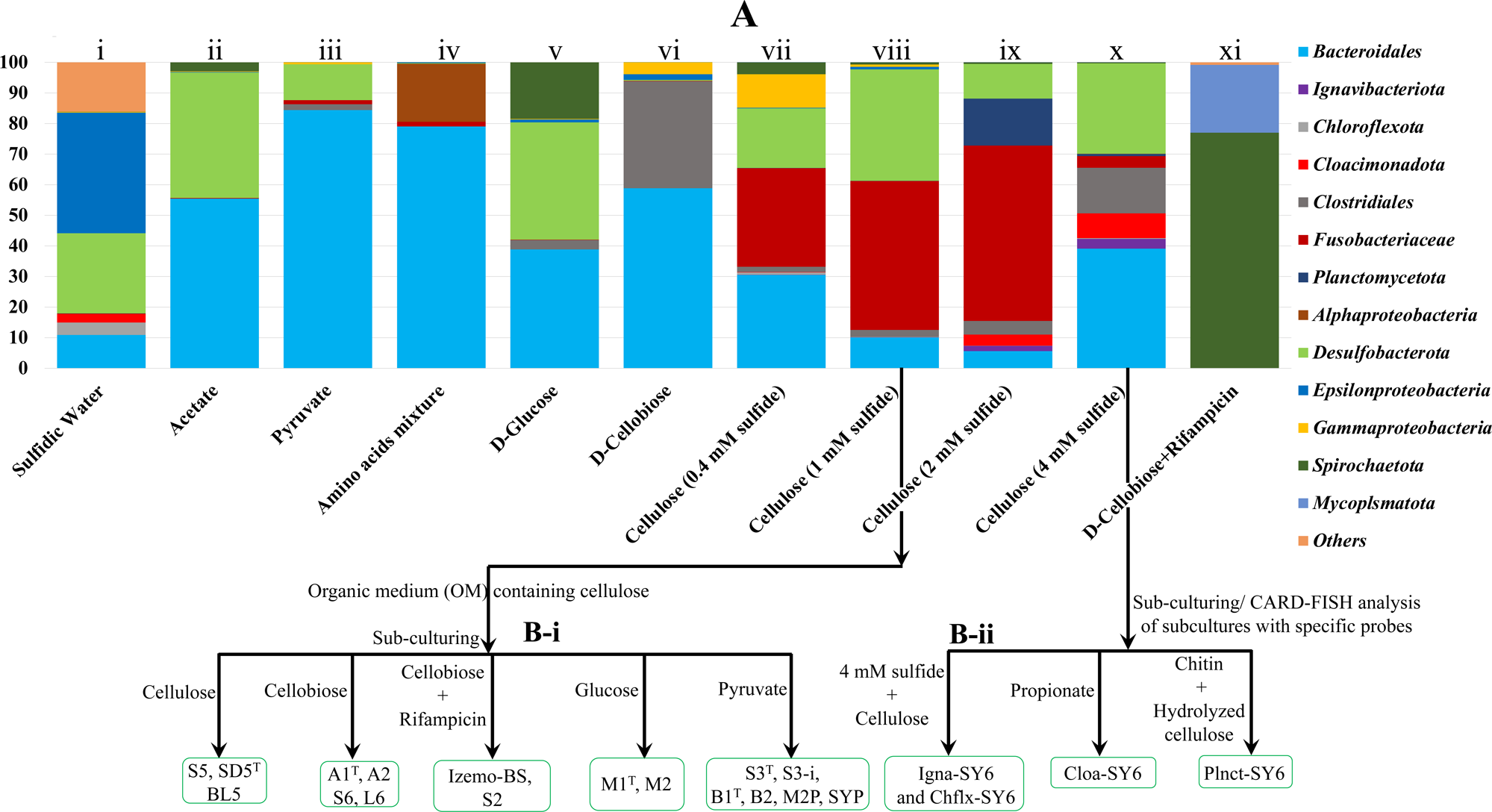
Schematic overview of the cultivation strategy of novel anaerobic microbial taxa from deep (2,000 m) sulfidic waters of the Black Sea. **(A)** Relative abundance (as determined by 16S rRNA gene amplicon sequencing) of the prokaryotic phyla in different enrichments incubated with specific carbon substrates (acetate, pyruvate, amino acid mixture, D-galactose, D-cellobiose, cellulose) at different growth conditions (i.e., 1-4 mM of sulfide at 10-20 °C). A repeated sub-culturing and streaking on various growth media from the master enrichment (A-viii and x) resulted in the pure (strains affiliated to the phyla *Fusobacteriota*, *Bacillota*, *Spirochaetota*, *Bacteroidota*) and enrichment cultures (*Cloacimonadota*, *Planctomycetota*, *Ignavibacteriota* and *Mycoplasmatota*) of novel piezotolerant anaerobic bacteria **(B)** Physiological characterization of all the novel taxa carried out at atmospheric pressure (0.1 MPa) and at elevated hydrostatic pressure (up to 50 MPa) by using pressure bottles and high-pressure cultivation assembly, respectively.

**Fig. 2:**
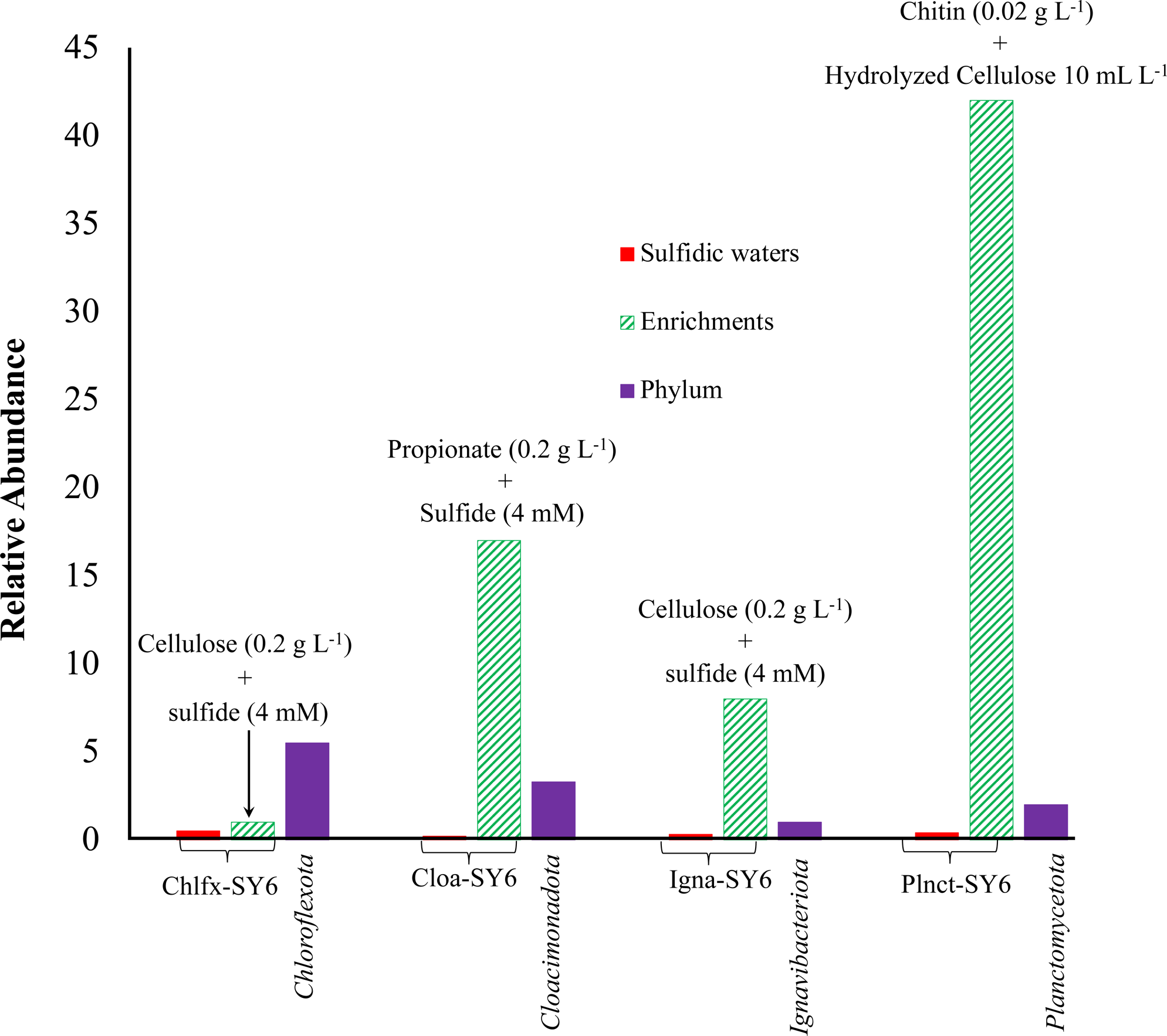
Enrichment of *Chloroflexota* bacterium Chflx-SY6, *Cloacimonadota* bacterium Cloa-SY6, *Ignavibacteriota* bacterium Igna-SY6, and the *Planctomycetota* bacterium Plnct-SY6 from the deep sulfidic waters of the Black Sea. The relative abundance (as determined by 16S rRNA gene amplicon sequencing) of the *Chloroflexota* bacterium Chflx-SY6, *Cloacimonadota* bacterium Cloa-SY6, *Ignavibacteriota* bacterium Igna-SY6, and the *Planctomycetota* bacterium Plnct-SY6 in the original sea water from 2000 m (red bars) and in the enrichment cultures (green hatched bars) using cellulose, propionate, cellulose, and chitin containing media, respectively. For reference, the relative abundance of the phyla in the original Black Sea water (purple bars) is indicated. The enrichment factor is shown in green hatched bars.

**Fig. 3:**
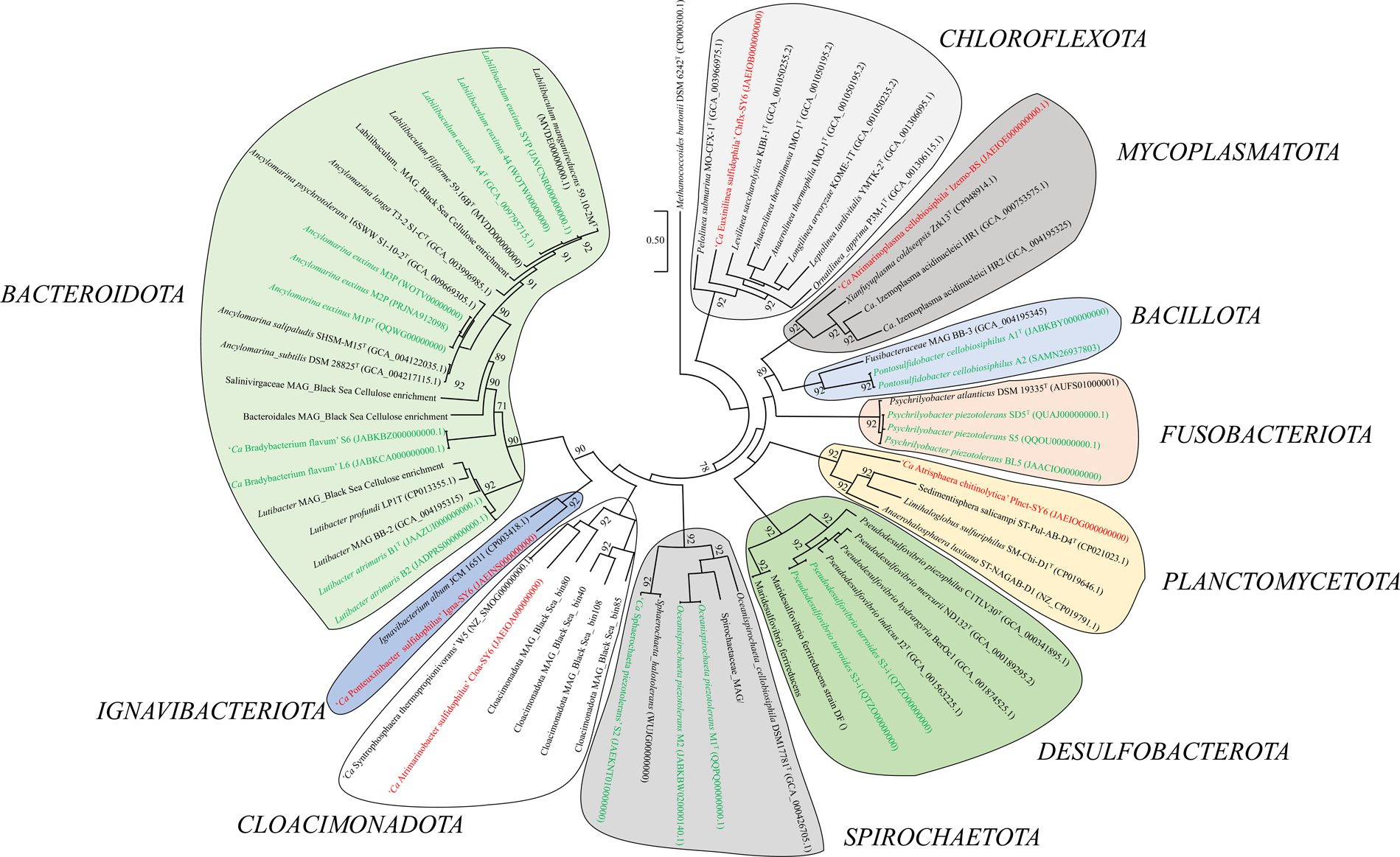
Phylogenomic tree showing the affiliation of novel taxa obtained from deep sulfidic waters of the Black Sea. The tree was built by the maximum likelihood method using the core gene phylogenomic tree constructed using UBCG pipeline^87^. The genome sequence accession numbers are shown between parentheses. Sequences highlighted in green color represent isolates obtained in this in this work, while taxa highlighted in red color represent strains from enrichments for which a proper MAG was obtained. The other sequences (in black) are used for reference. Numbers at nodes represent bootstrap values (1 refers to 1000). The length of the bar indicates five nucleotide substitutions per 100 nucleotides. *Methanococcoides burtonii* DSM 6242^T^ (CP000300.1) genome sequence was used as the outgroup for phylogenomic analysis.

### 2.1 Organic substrate utilization provides clues on the trophic behavior

We classified the isolated and enriched taxa into three categories, i.e., primary degraders, fermenters, and terminal oxidizers, based on their utilization and/or degradation capabilities of various substrates, and on the presence of various genes in their (meta)genomes capable of encoding enzymes involved in substrate utilization and/or degradation (Tables S1,S3-S5,S8-S41).

**Primary degraders.** *P. piezotolerans* strain S5 utilized a wide range of carbon sources, including cellulose, under strict anaerobic conditions, as described earlier for strains SD5^T^ and BL5^6^. Such properties were further supported by the presence of a variety of carbohydrate-active enzymes (CAZymes, n=77) encoded in its genome and 18 novel putative glycosyl hydrolases (GHs) with unknown substrate categories (Table S15). These could be involved in the hydrolysis of various carbon sources (carbohydrates) as observed in the culture-based analyses. Most notably, this was the only culture obtained in our isolation efforts showing cellulolytic activity (Fig. S12), even though genes encoding cellulases were detected in the genomes of other isolates (Table S15-S38). Moreover, an earlier study reported that *Psychrilyobacter* spp. constitutes up to 32-58% of the cellulose degrading microbial community in the deep sulfidic waters of the Black Sea^6^. Thus, it is anticipated that *P. piezotolerans* might be an important anaerobe involved in the hydrolysis of complex OM sources like cellulose in the sulfidic waters of the Black Sea.

*Clostridiales* bacteria strains A1^T^ and A2 grew rapidly (doubling time: 2-3 h) on cellobiose but twice as slow as on simpler carbon sources like glucose, while no growth occurred with pyruvate (Fig. S13). In accordance with their preference for cellobiose, genes encoding cellobiose-specific transporters (Fig. 4; Fig. S14) and cytoplasmic enzymes (e.g., β-glucosidase B, cellobiose 2-epimerase, β-glucosidase BoGH3B, 6-phospho-β-glucosidase, aryl-phospho-β-D-glucosidase BglC; Table S18-S19), able to convert cellobiose into less complex forms of carbon substrates like glucose, were identified in their genomes. The genomes also encoded a variety of CAZymes (A1^T^ and A2, n=86), including four amylases (Table S18-S19), but starch was not used as a substrate under the culture conditions tested here. A substrate preference for cellobiose was not evident in any other culture obtained in this study, in line with the absence of genes encoding for cellobiose-specific transporters in their genomes (Fig. 4). Thus, it may be that the *Clostridiales* bacteria strains A1^T^ and A2 may find a niche as specific cellobiose-degraders in sulfidic waters of the Black Sea.

**Fig. 4:**
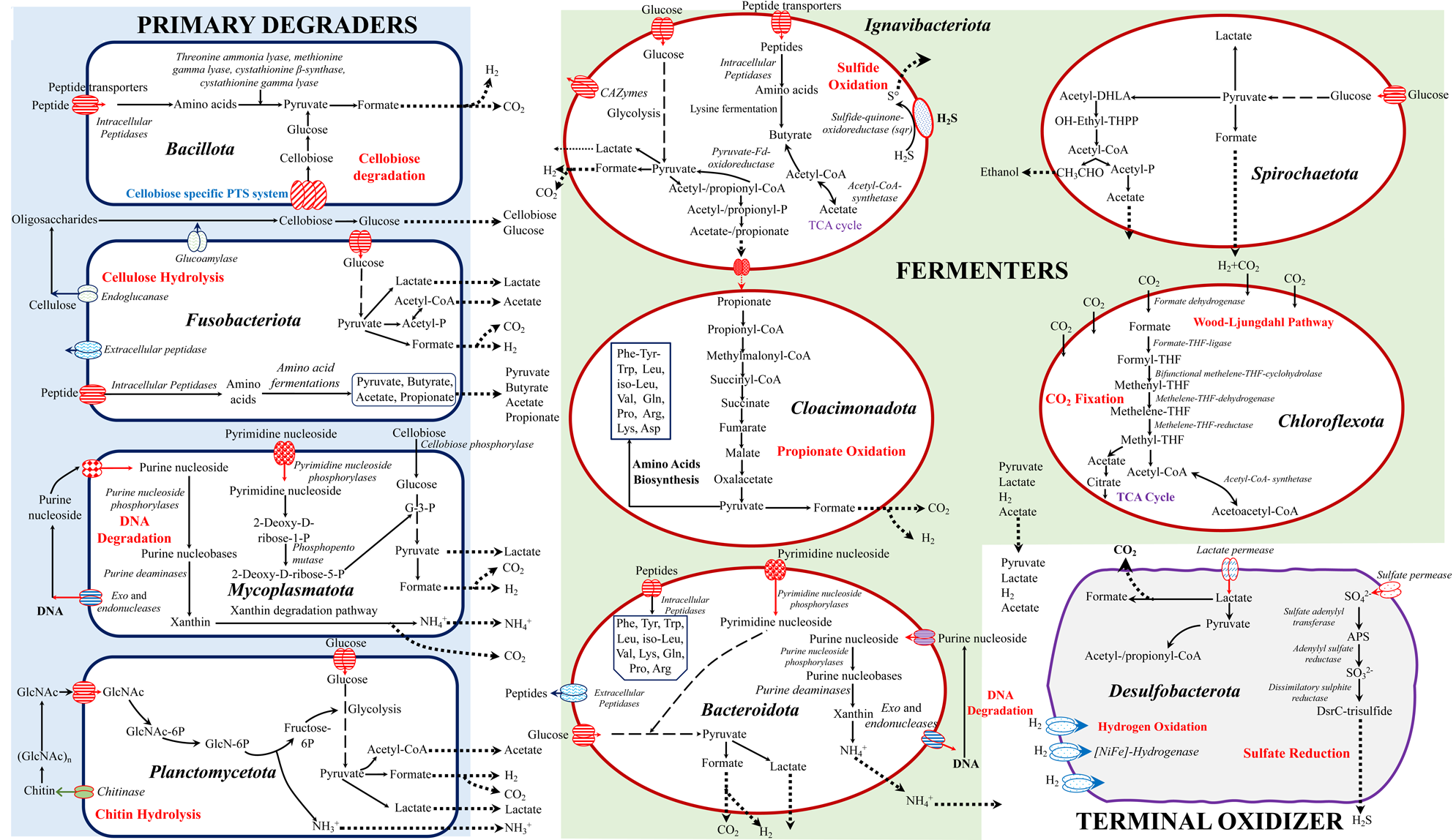
Metabolic models for labile OM (i.e., polysaccharides, proteins, nucleic acids, and lipids) degrading anaerobic microorganisms in the deep sulfidic waters of the Black Sea. The metabolic models are constructed based on the physiological and genomic (meta) analyses of pure cultures of the members of the phyla *Fusobacteriota* (*Psychrilyobacter piezotolerans* S5 and SD5^T^ and BL5), *Bacillota* (*Clostridiales* bacteria strains A1^T^ and A2), *Spirochaetota* (*Oceanispirochaeta* sp. strains M1^T^ and M2; *Sphaerochaeta* sp. strain S2), *Bacteroidota* (*Lutibacter* sp. strains B1^T^ and B2; *Lentimicrobiales* bacteria strains S6 and L6; *Ancylomarina euxinus* strain M2P, M3P and M1P^T^; *Labilibaculum euxinus* strains SYP, A4^T^ and 44), and *Desulfobacterota* (*Pseudodesulfovibrio* sp. strains S3^T^ and S3-i) and the enrichment cultures of the members of the phyla *Planctomycetota* (*Planctomycetota* bacterium Plnct-SY6), *Ignavibacteriota* (*Ignavibacteriota* bacterium Igna-SY6), *Mycoplasmatota* (Mycoplasmatota bacterium Izemo-BS) *Cloacimonadota* (*Cloacimonadota* bacterium Cloa-SY6) and *Chloroflexota* (*Chloroflexota* bacterium Chflx-SY6). Members of the phyla *Bacillota*, *Fusobacteriota*, *Mycoplasmatota*, and *Planctomycetota* are classified as primary degraders, simplifying larger biomolecules of organic matter such as carbohydrates (e.g., cellulose and chitin), proteins, nucleic acids, and lipids into simpler forms. These simpler forms are then further utilized or degraded by fermenters belonging to the phyla *Ignavibacteriota*, *Cloacimonadota*, *Bacteroidota*, *Spirochaetota*, and *Chloroflexota*. The fermented products are subsequently oxidized by terminal oxidizers (members of the phylum *Desulfobacterota*). GlcNAc, *N*-acetyl-D-glucosamine; GlcNAc-6P, *N*-acetyl-D-glucosamine-6-phosphate; TCA, tricarboxylic acid cycle; G-3-P, glycerol-3-phosphate.

The genome of *Planctomycetota* bacterium Plnct-SY6 contained 276 genes encoding CAZymes [glycosyl hydrolase (GHs; n=169); glycosyl transferase (GTs; n=45); carbohydrate esterase (CEs; n=6); carbohydrate binding modules (CBMs; n=49); auxiliary activities family (AAs; n=1); polysaccharide lyases (PLs, n=6); Table S20], including four copies of putative chitinases (GH20_e12; Table S20; Fig. 4; Fig. S15) and a complete set of enzymes required for the utilization of *N*-acetylglucosamine (GlcNAc; Fig. S15; Table S22). Both chitinase and N-acetylglucosaminidase are involved in the hydrolysis of chitin to oligomers and subsequent production of GlcNAc monomers (Fig. 4). The increased cell densities of strain Plnct-SY6 in a growth medium amended with chitin (0.02 gL^−1^) is in line with their affinity for chitin. This observation is consistent with previous studies reporting the increase in specific *Planctomycetes* members (class *Phycisphaerae*) in incubations with chitin-coated beads in the deep sulfidic waters of the Black Sea^14^, supporting our findings. To the best of our knowledge, only one confirmed chitinolytic planctomycete, *Fimbriiglobus ruber* SP5^T^, has been isolated but it originates from a peat bog^27^. Hence, our data suggests that strain Plnct-SY6 is the first marine planctomycete capable of degrading chitin and may have a significant role in mineralizing this abundant biopolymer in the deep sulfidic waters of the Black Sea.

Members of the phylum *Mycoplasmatota* are recognized for their ability to degrade DNA in marine environments^5,8,9^. We also identified genes encoding for exo- and endonucleases in the metagenome assembled genome (MAG) of the *Mycoplasmatota* bacterium Izemo-BS, which is consistent with this specialization (Table S21 and Fig. S16). Additionally, this bacterium may play a role in breaking down other, more structurally less complex carbon substrates such as sucrose, as its genome contains ten different genes encoding enzymes belonging to the GH13 family (Table S22), which are involved in the hydrolysis of sucrose.

**Fermenters:** The strains affiliated with the phyla *Spirochaetota* (M1^T^, M2 and S2) and *Bacteroidota* (B1^T^, B2, S2, M2P, SYP, S6 and L6), as shown in Fig. 3, were classified as fermenters based on their growth/substrate utilization characteristics.

Strains M1^T^ and M2 exhibited fast growth on simple molecules like glucose (doubling time 2-3 h) and pyruvate (with a doubling time; 1.5-2 h; Fig. S17-S18 and Table S1). However, they were unable to grow on more complex carbon sources like cellulose, cellobiose, and starch. Nonetheless, their genomes contain a variety of CAZymes, such as 114 GHs, 41 GTs for strain M1^T^, and 42 GTs for strain M2, as well as 8 CEs, 5 CBMs, 1AA, and 1 PL, (Tables S23-S24). Some of these CAZymes may potentially be involved in the hydrolysis of complex carbon sources like starch and cellulose.

Similar observations were made for *Lutibacter* sp. strains B1^T^ and B2. Both strains grew faster on pyruvate (doubling time 1.5-2 h; Fig. S19) than on glucose (doubling time 2-4 h; Fig. S19) or cellobiose (doubling time 3-6 h; Fig. S19), despite having a diverse range of CAZymes that include 72 GHs, 47 GTs, 8 CEs, 18 CBMs, 1 AA, and 3 PLs (Table S25-S26). Although their genomes contain genes that potentially encode enzymes that hydrolyze starch (the GH31 family, n=3) and sucrose (the GH13 family, n=7), the strains did not exhibit growth on these substrates.

Both *A. euxinus* strain M2P and *L. euxinus* strain SYP grew well on pyruvate (doubling time1.5-2.0 h; Fig. S20A-S20B) and relatively slower on glucose (doubling time, 1.8-2.0 h; Fig. S20A-S20B), but could not grow on cellobiose, cellulose, chitin, or starch. Although the genomes of strains M2P and SYP encoded 92 and 243 CAZymes, respectively (Table S27, S29), they were not able to utilize these complex carbon substrates. Specifically, the genome of SYP encoded CAZymes involved in cellulose utilization, such as those belonging to the GH5 family (n=5) (Table S29), but cellulolytic activity was not observed. Strain M2P encoded other CAZymes, including GHs, GTs, CEs, CBMs, AA, and PLs, (Table S27), which all could be involved in the hydrolysis of complex carbon sources, but such activities were not observed in our culture.

The *Lentimicrobiales* bacterial strains S6 and L6 were able to utilize various carbon sources, such as starch, glucose, sucrose, and pyruvate (Table S10). Their genomes encoded a total of 181 CAZymes, including 80 GHs, 74 GTs, 9 CEs, 17 CBMs, 2 AAs, and 1 PL. This is in line with their ability to grow on these carbon substrates (Table S31-S32). Despite their metabolic versatility, both strains exhibited relatively slow growth with doubling times of 2-3 days. Together with their relatively low abundance (approximately 1%; Fig. 1B) in the enrichments, this suggests a rather limited role in the degradation of OM.

To date, members of *Cloacimonadota* have not been enriched in any marine environment, and previous reports of their presence are based on incubations and metagenomic analyses ^20,28,29^. Here, we report an increase in the cell density of Cloa-SY6, a member of *Cloacimonadota*, in a medium containing propionate (0.2 gL^−1^), indicating its ability to use this substrate. The presence of genes involved in propionate oxidation in its MAG (Fig. S21) further supports this. Its MAG encoded a relatively limited number of 64 CAZymes, with most of them being GTs (n=45; Table S33) involved in the initiation and elongation of glycan chains^30^. Only 11 CAZymes were classified as GHs, indicating a limited carbohydrate utilization capability. Therefore, in the deep sulfidic waters of the Black Sea, *Cloacimonadota* bacteria may occupy a unique niche as anaerobic propionate oxidizers.

The *Ignavibacteriota* bacterium Igna-SY6 was enriched in an oligotrophic cellulose medium with a relatively high sulfide concentration (4 mM) and low temperature (10°C; Fig. 1A), indicating adaptation to cold and nutrient-poor environments. The bacterial MAG contained 246 genes that encode CAZymes, including GHs, GTs, CEs, CBMs, AAs, and PLs (Fig. S22; Table S34). The potential presence of CAZymes involved in cellulose, cellobiose, starch, and sucrose degradation suggests a role in carbohydrate utilization, as previously reported^31^. Similarly, the *Chloroflexota* bacterium Chflx-SY6 was also enriched using an oligotrophic medium (Fig. 1A). Its MAG encoded a variety of CAZymes (n=155), including GHs (n=82), GTs (n=60), CEs (n=8), CBMs (n=3), AAs (n=1), and PLs (n=1; Table S35), indicating its potential to degrade various extracellular carbohydrates (Fig. S23), which is consistent with its reported role in OM degradation^14^. Our findings suggest that both strains Igna-SY6 and Chflx-SY6 prefer oligotrophic conditions.

Overall, we observed that novel taxa belonging to various phyla possess remarkable abilities for breaking down specific carbohydrates. However, we also found that the mere presence of genes encoding for CAZymes may not result in their ability in hydrolyzing complex carbon sources like cellulose and chitin, at least not with the culture conditions employed in this study. This may suggest that context-specific factors may play a crucial role in determining the function of CAZymes, as exemplified by the cellulose biosynthetic cluster (BcsABZC) found in various cellulose-synthesizing microorganisms^32–34^. It is worth noting that this cellulose biosynthetic gene cluster not only regulates cellulose biosynthesis but also acts as a cellulase, highlighting the dynamic and versatile nature of CAZymes. Similar observations have also been made for *Escherichia coli*, which possesses numerous CAZymes involved in cellulose metabolism but does not exhibit a cellulolytic phenotype^32,33^, further underscoring the role of CAZymes in polysaccharide hydrolysis and utilization.

**Terminal oxidizers:** Bacterial strains belonging to the phylum *Desulfobacterota* (*Pseudodesulfovibrio* sp. strains S3^T^ and S3-i) utilized structurally less complex carbon substrates, such as lactate, pyruvate, and acetate (Table S5). Such simple organic molecules are typically derived from fermentative microorganisms involved in OM degradation (Fig. 4). Our strains indeed possessed specific genes in their genomes that reflect their ability to use these substrates (Fig. S24). Lactate was their preferred substrate, which can be oxidized to pyruvate by lactate dehydrogenase. Pyruvate is then further oxidized mainly by pyruvate: ferredoxin oxidoreductase, although their genomes also encode several alternative oxo-organic acid oxidoreductases that can react with pyruvate. Enzymes such as phosphate acetyltransferase and acetate kinase could lead to the production of ATP by substrate-level phosphorylation during the conversion of acetyl-CoA to CoA and acetate. In addition, the formate dehydrogenase enzyme complex encoded in their genomes could help in the oxidation of formate to CO_2_ and H_2_. These strains also possess the complete pathway for dissimilatory sulfate reduction (Fig. S24). Furthermore, these strains could use hydrogen as an energy source, indicating that they may actively remove hydrogen produced by fermentation processes in the enrichments and potentially in the sulfidic waters of the Black Sea.

### 2.2 Protein, lipid, and nucleic acid degradation

Strains S5, A1^T^, A2, M2P, SYP, B1^T^, B2, S6 and L6, belonging to different bacterial groups (Fig. 3), were also able to utilize protein (casein) as a source of energy and nutrients (Tables S4, S10-S12). In accordance with the observed proteolytic activity, we identified a variety of genes encoding peptidases [strain S5 (n=28), A1^T^ and A2 (n=23), M2P (n=27), SYP (n=24), B1^T^ and B2 (n=18), S6 and L6 (n=19)] (Table S40). Most of the peptidases were predicted to be intracellular. However, their genomes also encoded extracellular peptidases [strain S5 (n=1), A1^T^ and A2 (n=1 with multiple localization), M2P (n=3 with multiple localization), SYP (n=4 with multiple localization and n=1 extracellular), B1^T^ and B2 (n=3 with multiple localization), S6 and L6 (n=6 with multiple localization)], which might have aided in the protein hydrolysis. The genomes of strains S5, A1^T^, A2, M2P, SYP, B1^T^, B2, S6 and L6 encoded several aminotransferases and other amino acid-degrading enzymes (i.e., alanine dehydrogenases, cystathionine gamma-lyase, serine/threonine ammonia-lyase, tryptophanase, and methionine gamma-lyase; Fig. S12, S14, S24). These enzymes can convert amino acids to the keto-acids pyruvate and oxobutanoate, which could then be fermented to formate, and either acetate or propionate, respectively. The fermentation of amino acids by these strains to propionate, acetate, or formate, i.e., via aminotransferases and other amino acid degrading enzymes (i.e., L-threonine 3-dehydrogenase, cystathionine β-synthase, cystathionine gamma-lyase, serine/threonine ammonia-lyase, and methionine gamma-lyase) is possible.

In contrast, proteolytic activity was not observed in bacterial strains affiliated with the phylum *Spirochaetota* (strains M1^T^, M2, and S2), which is consistent with the absence of genes encoding extracellular peptidases (Table S40). Similarly, strains affiliated with the phylum *Desulfobacterota* (strains S3^T^, and S3-i) were also unable to hydrolyze protein (casein), but, in contrast, their genomes did encode extracellular peptidases (Table S40). These latter strains may have specialized in different protein sources under different conditions.

In addition to the confirmed proteolytic activities in pure cultures, we also identified a variety of peptidases encoded in the MAGs of *Cloacimonadota* bacterium Cloa-SY6, *Ignavibacteriota* bacterium Igna-SY6, *Chloroflexota* bacterium Chflx-SY6, *Planctomycetota* bacterium Plnct-SY6, and *Mycoplasmatota* bacterium Izemo-BS (Table S40). These peptidases suggest a potential role in protein degradation, but the majority were found to be localized in the cytoplasm. Hence, it is expected that these taxa may have a limited role in protein degradation in deep sulfidic waters. Nevertheless, a variety of aminotransferases and other amino acid degrading enzymes (i.e., alanine dehydrogenases, cystathionine gamma-lyase, serine/threonine ammonia-lyase, tryptophanase, and methionine gamma-lyase) indicate their potential role in amino acid degradation. These taxa can convert amino acids into the keto-acids pyruvate and oxobutanoate, which could then be fermented to formate, and either acetate or propionate, respectively.

The lipid-metabolizing capability of the novel taxa, both in pure and enrichment cultures, revealed the presence of numerous genes encoding lipases (Table S41). However, most of these lipases were predicted to be localized in the cytoplasm (Table S41). Notably, the genome of the *L. euxinus* strain SYP contained the highest number (n=15) of genes encoding lipases, which include a diverse array of enzymes such as rhamnogalacturonan acetylesterase RhgT, endo-1,4-beta-xylanase/feruloyl esterase, fumonisin B1 esterase, monoacylglycerol lipase, pectinesterase A, acetylxylan esterase, and carbohydrate acetyl esterase/feruloyl esterase (Fig. S25). These lipases were predicted to have multiple localizations along with a few unknown ones (Table S41). Therefore, we postulate that the *L. euxinus* strain SYP may play a significant role in lipid degradation in the deep sulfidic waters of the Black Sea, whereas the lipases encoded in the genomes of other taxa are restricted to intracellular lipid recycling, and, therefore, less important in terms of the biogeochemical carbon cycling in euxinic waters of the Black Sea.

We also identified a variety of nucleases encoded in the genomes of pure cultures [strain S5 (n=51), A1^T^ and A2 (n=58), M2P (n=43), SYP (n=45), B1^T^ (n=42), S6 (n=40), and L6 (n=39)] and MAGs of the enrichment cultures [*Cloacimonadota* bacterium Cloa-SY6 (n=33), *Ignavibacteriota* bacterium Igna-SY6 (n=55), *Chloroflexota* bacterium Chflx-SY6 (n=69), *Planctomycetota* bacterium Plnct-SY6 (n=40), and *Mycoplasmatota* bacterium Izemo-BS (n=43)]. These included proteins involved in the cleavage of nucleobases into sugar and base components. Purine deoxyribonucleoside phosphorylases were encoded in the genomes of most strains, while genes encoding pyrimidine deoxyribonucleoside phosphorylases were also detected (Table S39). Multiple copies of adenosine deaminase genes were identified, indicating their ability to use these molecules (Table S39). Xanthine dehydrogenases, key enzymes required for conversions of purine bases to xanthine, a common intermediate of purine breakdown, were also encoded by most of the genomes/MAGs (Table S39). Most of the nucleases identified were predicted to be localized to the cytoplasm, indicating their role in the intracellular recycling of nucleic acids. However, a variety of nucleases with unknown and multiple localization [(e.g., bifunctional oligoribonuclease, 5’-nucleotidase SurE, endoribonuclease (EndoA, YbeY, Maz F) AMP nucleosidase, exodeoxyribonuclease 7 ribonuclease (H, HI, M5, Y, BN and 3)], extracellular ribonuclease, AMP nucleosidase (Table S39) were encoded by the genomes of *Clostridiales* bacteria strains A1^T^ and A2, *A. euxinus* strain M2P, *L. euxinus* strain SYP, *Lutibacter* sp. strain B1^T^ and *Mycoplasmatota* bacterium Izemo-BS. These nucleases are expected to be involved in the degradation of extracellular nucleic acid in the deep sulfidic waters of the Black Sea. In good agreement with our analyses, a recent stable isotope probing (SIP) based study also reported that novel *Clostridiales* bacteria (closely related to strains A1^T^ and A2), *Mycoplasmatota* members, and *Lutibacter* spp. are crucial for nucleic acid degradation in sulfidic marine sediments^5^.

Based on these results and earlier work ^4,5,8,9,13–15^, it is anticipated that *Psychrilyobacter* spp., members of the order *Clostridiales* (closely related to strains A1^T^ and A2), *Marinifilaceae* members (closely related to *A. euxinus* and *L. euxinus*), *Lutibacter* spp. (closely related to strains B1^T^ and B2) and *Lentimicrobiales* members (related to strains S6 and L6), *Sphaerochaeta* spp. (closely related to strains S2), *Oceanispirochaeta* spp. (closely related to strains M1^T^ and M2), *Cloacimonadota* bacterium Cloa-SY6, *Ignavibacteriota* bacterium Igna-SY6, *Chloroflexota* bacterium Chflx-SY6, *Planctomycetota* bacterium Plnct-SY6 and *Mycoplasmatota* bacterium Izemo-BS could be key anaerobes involved in the turnover of “fresh” and labile OM (i.e., carbohydrates, proteins, lipids and nucleic acids) passaging through the deep sulfidic waters of the Black Sea.

### 2.3 Adaptation of novel microbial taxa to the *in*-*situ* conditions of deep sulfidic waters

Deep oceanic waters are characterized by high hydrostatic pressure and the microorganisms inhabiting them must have adapted to these harsh conditions. Despite this, our understanding of the mechanisms by which these microorganisms adapt remains limited, with only a few studies available^6,11,35–39^. To gain insight into their potential for adaptation, we investigated the growth of various bacterial strains, affiliated with the phyla *Fusobacteriota* (strain S5), *Bacillota* (strains A1^T^ and A2), *Spirochaetota* (strains M1^T^ and M2), *Bacteroidota* (strains B1^T^, S6), and *Desulfobacterota* (strains S3^T^, and S3-i) at hydrostatic pressures ranging from 0.1 to 60 MPa. We found that all strains were able to thrive at pressures up to 20 MPa, which represents the *in-situ* pressure conditions at 2,000 m depth, which is the maximum depth of the Black Sea. However, as the pressure increased beyond 20 MPa, the growth rates of the strains slowed down (Fig. 5), although they were still able to withstand a hydrostatic pressure of 50 MPa.

**Fig. 5:**
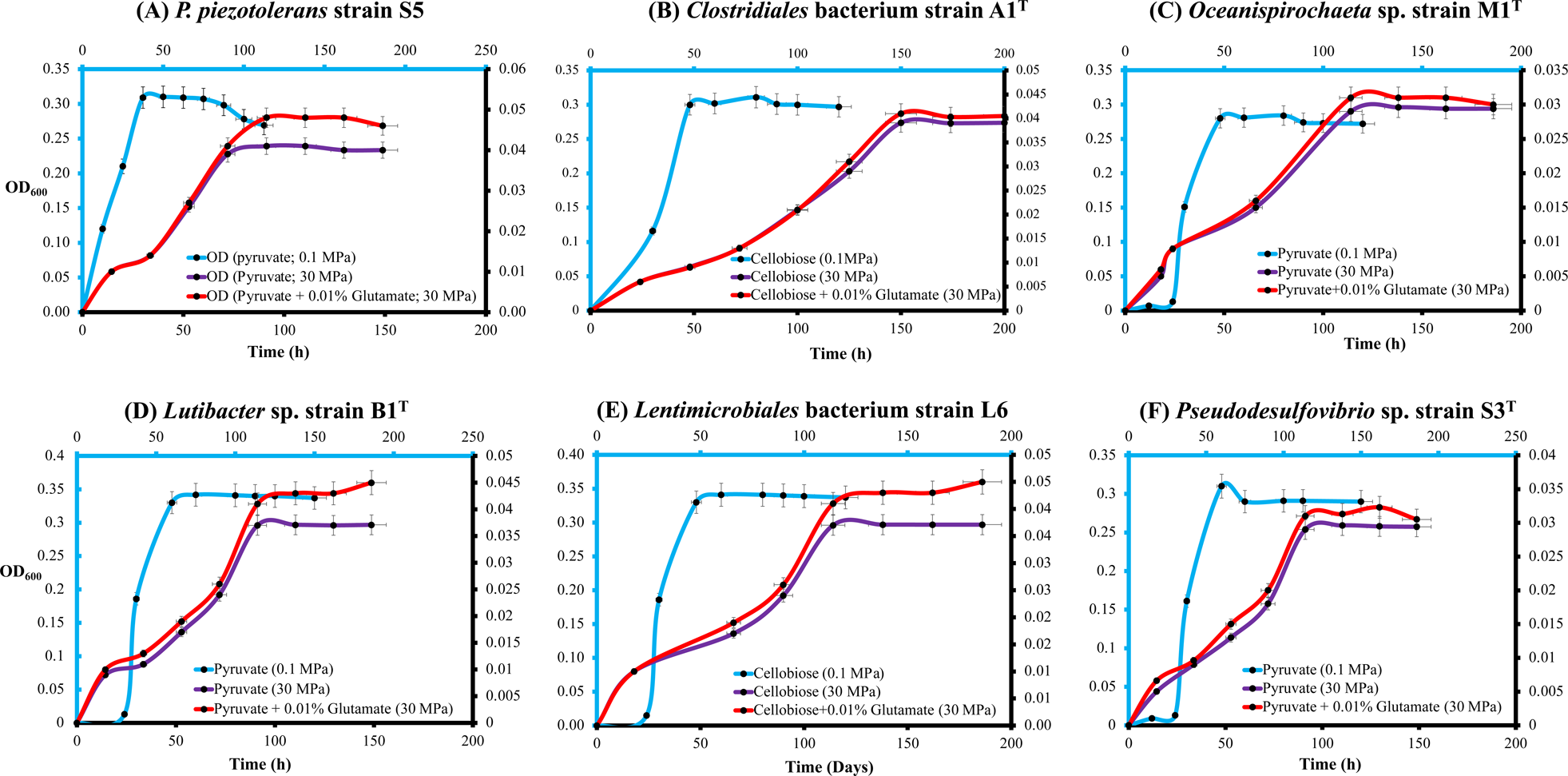
Growth curves of various novel taxa isolated from sulfidic waters of the Black Sea at optimal growth conditions (20 °C and 0.1 MPa), and at elevated hydrostatic pressure (30 MPa) with and without the piezolite glutamate (0.01%). **(A)** *Psychrilyobacter piezotolerans* strain S5, **(B)** *Clostridiales* bacterium strain A1^T^, **(C)** *Oceanispirochaeta* sp. strain M1^T^, **(D)** *Lutibacter* sp. strain B1^T^, **(E)** *Lentimicrobiales* bacterium strain L6, **(F)** *Pseudodesulfovibrio* sp. strain S3^T^. Pyruvate was the carbon substrate for growth in all experiments. Key: OD_600_ = optical density measured at 600 nm. Error bars refer to the standard error. The light, blue-colored x-axis and y-axis represent growth at 0.1 MPa, while the black-colored x-axis and y-axis denote growth at 30 MPa.

The growth of selected cultures (*P. piezotolerans* strain S5, *Oceanispirochaeta* sp. strain M1^T^, *Lutibacter* sp. strain B1^T^, *Lentimicrobiales* bacteria strain L6 and *Pseudodesulfovibrio* sp. strain S3^T^) at elevated hydrostatic pressure (>20 MPa) was improved when a piezolite (glutamate)^35^ was added to the growth medium (Fig. 5). Glutamate has been shown to promote growth at elevated hydrostatic pressure in *Desulfovibrio* spp.^35^ and *Psychrilyobacter* spp.^6^, as it is an important precursor for ectoine biosynthesis^40^, which helps to maintain osmotic balance under stressed conditions^41^. Therefore, glutamate is thought to be an important piezolite for adapting to elevated hydrostatic pressure. However, our experiments indicated that the piezolite effect of glutamate varied among different strains (Fig. 5). For instance, *Lentimicrobiales* bacterium strain L6 showed the maximum effect (as monitored by the increase of optical density, OD_600_; Fig. 5E), whereas the smallest effect was observed for strain M1^T^ (Fig. 5C). No significant effect was observed on the growth of *Clostridiales* bacterium strain A1^T^ at elevated hydrostatic pressure, although this bacterium possesses the genes encoding enzymes involved in the GS-GOGAT pathway (associated with piezolite production; Fig. S14). This suggests that bacteria related to strain A1^T^ might have different strategies to deal with elevated hydrostatic pressure. Recent observations have shown that piezotolerant members of the phylum *Bacteroidota* take up trimethylamine (TMA) through a TMA transporter, TmaT, and oxidize intracellular TMA into trimethylamine N-oxide (TMAO) using a TMA monooxygenase, MpTmm^36^. The TMAO produced accumulates in the cell, functioning as a piezolite, improving both growth and survival at elevated hydrostatic pressure. We examined the genomes of all pure cultures and MAGs of enrichment cultures (Table S6), but we could not detect genes coding for the TMA monooxygenase. Therefore, it is expected that these strains may harbor alternative mechanisms for dealing with elevated hydrostatic pressure conditions.

The deep waters of the Black Sea are also characterized by a high sulfide concentration (400 μM)^16^, resulting from the activity of sulfate-reducing microorganisms (SRM)^42^. Such high sulfide levels are toxic to most life forms and form a stress that microorganisms dwelling in the deep Black Sea waters must overcome. We observed that the pure cultures affiliated with the phyla *Spirochaetota* (strains M1^T^, M2 and S2), *Bacteroidota* (strains B1^T^, B2, S6, L6, M2P, and SYP), *Bacillota* (strains A1^T^ and A2), *Desulfobacterota* (strains S3^T^ and S3-i), and *Fusobacteriota* (strains SD5^T^ and BL5)^6^ were able to grow up to sulfide levels as high as 7.0, 7.0, 7.0, 9.0, and 32 mM, respectively (Fig. 6). In general, such a high level of sulfide tolerance is expected to be created through the detoxification of sulfide through enzymatic oxidation associated with the highly conserved sulfide:quinone oxidoreductase (sqr) pathway, and subsequent excretion of oxidized sulfur species (typically sulfate)^43–47^. However, we did not observe sulfide oxidation for these strains in culture.

**Fig. 6:**
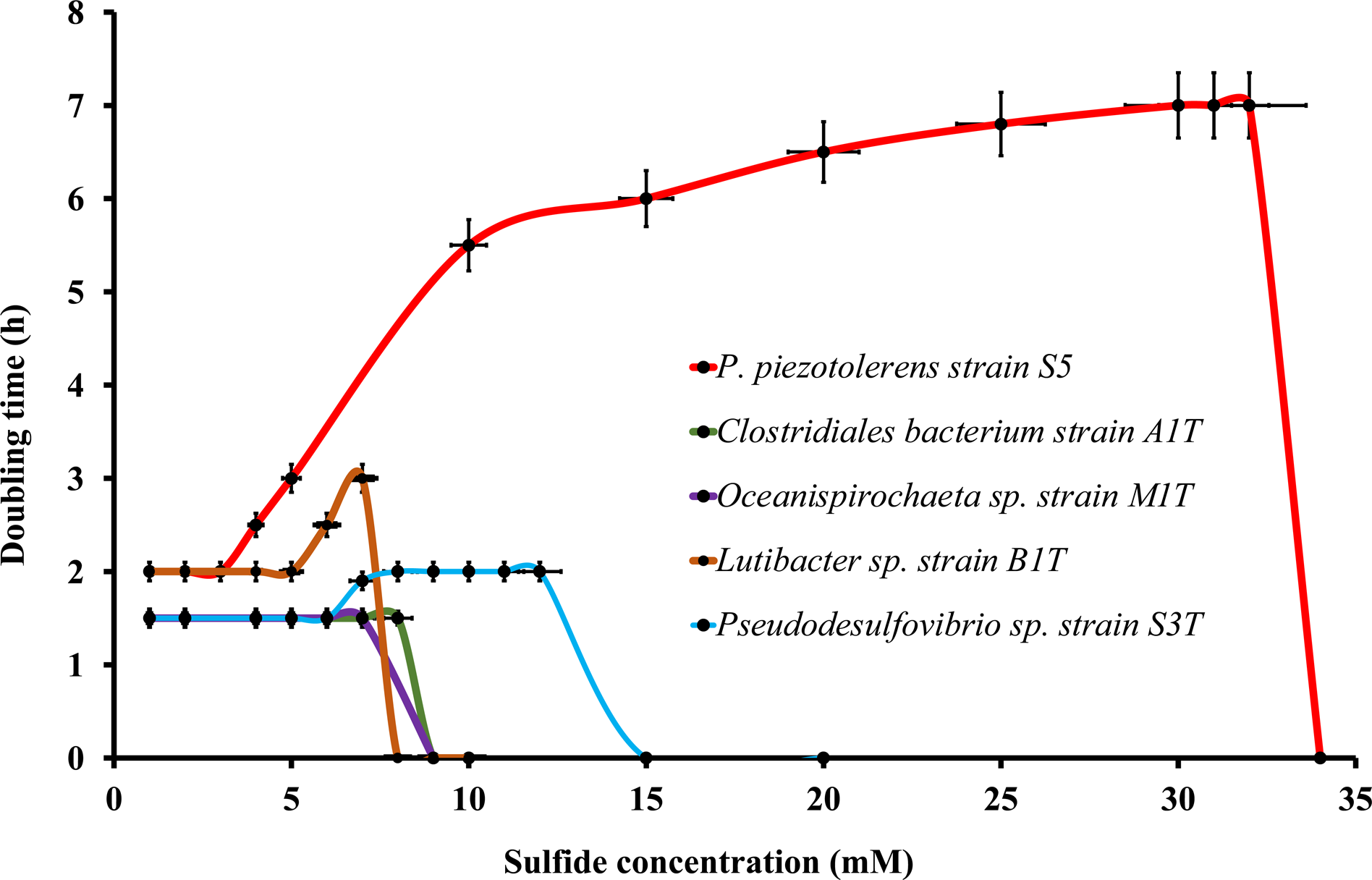
Doubling time of various novel taxa (*Psychrilyobacter piezotolerans* strain S5, *Clostridiales* bacteria strain A1^T^, *Oceanispirochaeta* sp. strain M1^T^, *Lutibacter* sp. strain B1^T^, *Lentimicrobiales* bacterium strain L6, *Pseudodesulfovibrio* sp. strain S3^T^) at increasing sulfide concentrations Key: OD_600_ = optical density measured at 600 nm. Error bars refer to the standard error.

Although the enrichment cultures of various bacteria, including *Cloacimonadota* bacterium Cloa-SY6, Chloroflexota bacterium Chflx-SY6, *Ignavibacteriota* bacterium Igna-SY6, and *Planctomycetota* bacterium Plnct-SY6, were observed to grow at higher sulfide concentrations (>2 mM; Fig. 1A), we could not find genes related to sqr in any of these taxa, except for *Ignavibacteriota* bacterium Igna-SY6. The MAG of *Ignavibacteriota* bacterium Igna-SY6 contained two sqr genes that might be involved in utilizing sulfide as an electron donor, ultimately helping to alleviate sulfide toxicity by forming sulfur/polysulfide. We detected genes encoding rhodanese (EC 2.8.1.1) in the novel taxa affiliated with *Fusobacteriota* (*P. piezotolerans* strain S5, SD5^T^ and BL5), *Spirochaetota* (*Oceanispirochaeta* sp. strains M1^T^ and M2; *Sphaerochaeta* sp. strain S2), *Bacteroidota* (*Lutibacter* sp. strains B1^T^, B2, *Lentimicrobiales* bacteria strains S6 and L6; *A. euxinus* strain M2P; *L. euxinus* strain SYP), *Bacillota* (*Clostridiales* bacteria A1^T^ and A2), and *Desulfobacterota* (*Pseudodesulfovibrio* sp. strains S3^T^ and S3-i). Rhodanese has been shown to be involved in sulfur trafficking and oxidation in bacteria^46,47^, indicating its role in sulfur metabolism, and potentially in alleviating sulfide toxicity. Moreover, a complete set of genes encoding the enzymes involved in the heterodisulfide reductase (HDR) system, acting as an elemental sulfur oxidation enzyme in the cytoplasmic space of bacteria and archaea^43–47^, was also identified in these strains, which might also be potentially involved in alleviating sulfidic toxicity. Nevertheless, further studies are required to gain deeper insights into the molecular mechanisms of sulfide detoxification in these strains.

### 2.4 Environmental distribution and ecological relevance of the isolated and enriched strains

We examined the distribution of isolated and enriched taxa by examining the pool of 16S rRNA gene amplicon sequences (fragments of 304 bp) detected in the sulfidic waters at 2,000 m. 16S rRNA gene sequences affiliated with *P. piezotolerans* strain S5, *Clostridiales* bacteria strains A1^T^ and A2, *Oceanispirochaeta* sp. strains M1^T^ and M2, *Lentimicrobiales* bacteria strains L6 and S6, *Lutibacter* sp. strains B1^T^ and B2, and *Mycoplasmatota* bacterium Izemo-BS were detected in relatively low abundance (i.e., representing <1.0% of the total bacterial+archaeal 16S rRNA gene reads), whereas sequences belong to *A. euxinus* strain M2P and *L. euxinus* strain SYP occurred in much higher abundance (∼9%; Fig. 1A), representing important community members. 16S rRNA gene amplicon sequences affiliated with the phyla *Cloacimonadota*, *Chloroflexota*, *Ignavibacteriota* and *Planctomycetota* were in the range of 1-3% in the deep sulfidic waters of the Black Sea. However, the 16S rRNA gene fragment sequences affiliated with *Cloacimonadota* bacterium Cloa-SY6, *Chloroflexota* bacterium Chflx-SY6, *Ignavibacteriota* bacterium Igna-SY6 and *Planctomycetota* bacterium Plnct-SY6 were less than 1% (Fig. 1D).

Taxa affiliated with *P. piezotolerans* strain S5, *Clostridiales* bacteria strains A1^T^ and A2, *Oceanispirochaeta* sp. strains M1^T^ and M2, *Lentimicrobiales* bacteria strains L6 and S6, *Lutibacter* sp. strains B1^T^, B2 and *Mycoplasmatota* bacterium Izemo-BS were detected low in abundance. However, their physiological, metabolic, and genomic properties indicated that they have potential to become abundant under copiotrophic (nutrient-rich) conditions occurring occasionally in the Black Sea^48–50^. For instance, *Clostridiales* bacteria strains A1^T^ and A2, *Oceanispirochaeta* sp. strains M1^T^ and M2, *A. euxinus* strain M2P, *L. euxinus* strain SYP, *Lutibacter* sp. strains B1^T^ and B2, *Pseudodesulfovibrio* sp. strain S3^T^ and S3-i and *M. ferrireducens* strain DF required a minimum of approximately 0.02 g L^−1^ of OM, such as yeast extract and tryptone, for their growth (Fig. 7), while their optimal growth occurred at around 5 g L^−1^ of OM (Fig. 5).

**Fig. 7:**
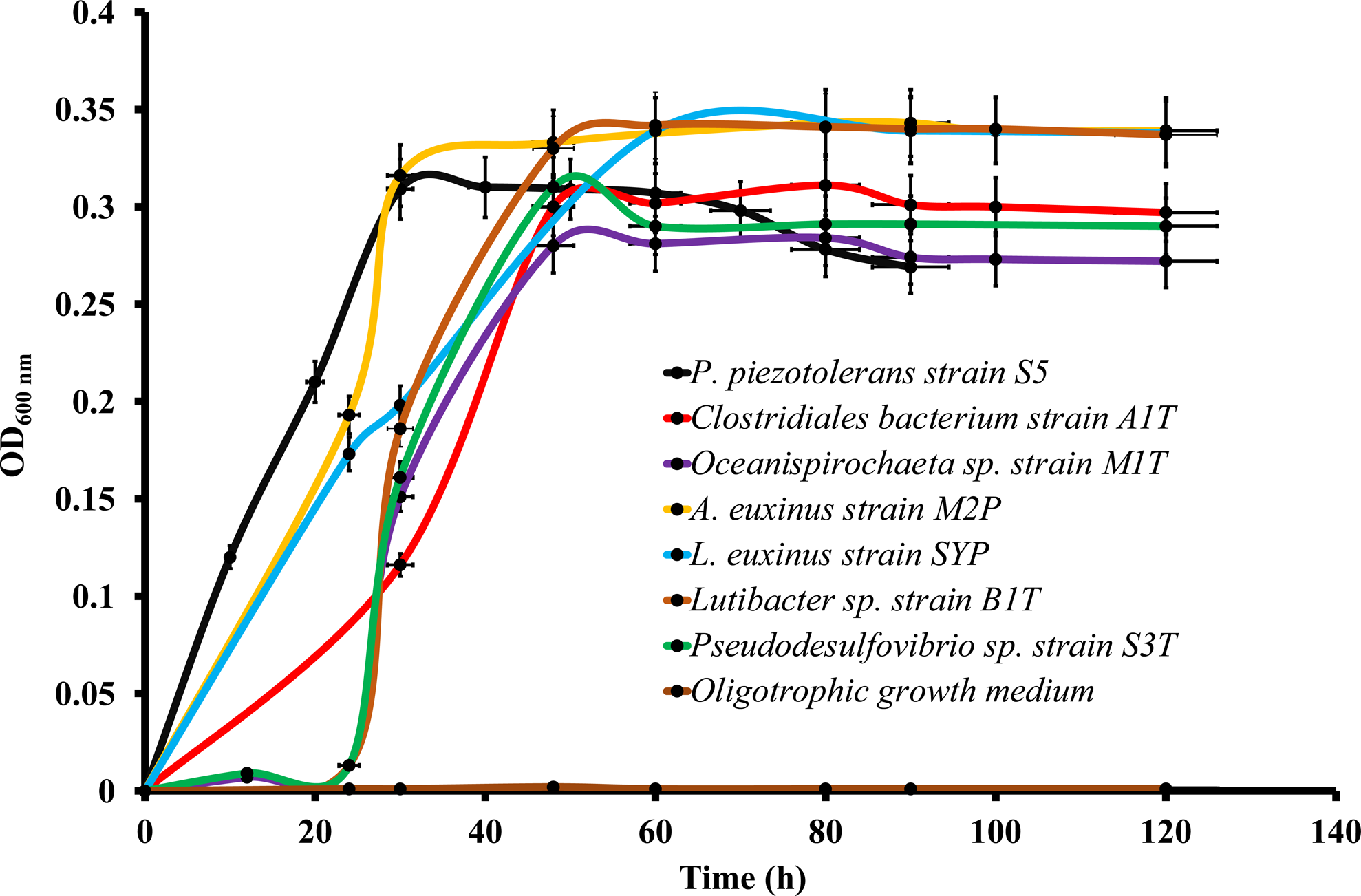
Growth of various novel taxa was observed at minimal (0.002 g. L^−1^) and optimal (5 g.L^−1^) concentrations of OM (yeast extract, tryptone and pyruvate/cellobiose). Pyruvate was used as the carbon source for the growth of all novel taxa, except for *Clostridiales* bacterium strain A1^T^ and *Lentimicrobiales* bacterium strain L6, which used cellobiose as the major carbon source. The optical density was measured at 600 nm (OD_600_), and error bars indicate the standard error.

Here, we further assessed the copiotrophic nature of these strains based on specific genomic signatures. These signatures have been suggested as a proxy for determining the ecological characteristics and trophic lifestyle of marine heterotrophic bacteria^51^. The genome sequences of these strains contain genes that encode the phosphotransferase system (PTS). This system is essential for regulating and transporting sugars, Na^+^ transporters, a wide array of other highly specific transporters, permeases, hydrolases, and motility and sensory systems. Copiotrophic bacteria widely present such transport systems and gene clusters, allowing them to locate and exploit transient microscale nutrient sources^51^. Therefore, we hypothesize that these strains may serve as “seed microbes” and become abundant (conditionally rare taxa) under copiotrophic conditions and play a significant role in the degradation of OM in the Black Sea.

## Conclusions

We enriched and isolated representatives of uncultivated microbial lineages that have the potential to break down OM under permanently sulfidic and high pressure conditions. These anaerobes belong to various physiological groups, including cellulose, chitin, and DNA degraders, fermenters, hydrogen-oxidizers, and sulfate reducers. We identified a range of genes encoding CAZymes, peptidases, nucleases, and lipases in the genomes of these novel piezotolerant anaerobes. However, not all cultures with such hydrolases exhibited hydrolytic properties, highlighting the need for context-dependent approaches to gain insights into their specific roles.

We found that piezolites, such as glutamate, could enhance growth under high hydrostatic pressure, but alternative and novel mechanisms likely exist to cope with elevated pressures. The permanently higher sulfide concentration in the deep waters of the Black Sea has resulted in the adaptation of microorganisms to an unusually high level of sulfide. Moreover, this study also uncovered a group of lipids that might potentially play an important role in the sustainable lifestyle of deep-sea anaerobes. However, more detailed analyses are required to gain deeper insights into their specific functions.

Overall, the novel anaerobes obtained in this study are adapted to high hydrostatic pressure and sulfide concentrations and could serve as important microorganisms for exploring the ecophysiological processes and adaptation strategies in deep sulfidic marine habitats.

## Materials and Methods

### Sampling sites, and 16S rRNA gene amplicon sequencing

A sampling survey was conducted to collect water from the Black Sea in 2017 and 2018 on board of the *R/V* Pelagia as previously described^6,11^. The study site was located at 42° 53.78’ N 30° 40.72’ E. A Sea-Bird SBE911C conductivity-temperature-depth (CTD) system equipped with a 24L×L12LL Niskin bottle rosette was used to collect water samples at a depth of 2,000Lm, while simultaneously measuring the conductivity, temperature, depth, and oxygen concentration. Water was immediately transferred into 2LL pressure bottles (*ca.*1.5-2.0Lbar), which was pressurized with N_2_, covered with aluminum foil to protect it from the light, and stored at 10 °C (*in situ* temperature) until the sample was used to start enrichment cultures. Water collected at 2,000Lm was filtered (2LL) through a 0.2 μm Sterivex™ filter for DNA analysis.

To analyze the total prokaryotic diversity, DNA was extracted from both the water column from the Black Sea and from the enrichment cultures themselves, and further amplified for 16S rRNA gene amplicon sequencing as described earlier^20^. 16S rRNA gene amplicon data were analyzed by the Cascabel pipeline^52^ including quality assessment by FastQC^53^, assembly of the paired-end reads with Pear^54^, library demultiplexing, OTU clustering and representative sequence selection (‘longest’ method) by diverse Qiime scripts^55^.

### Enrichment and isolation

Black Sea water from 2000 m was used for the subsequent enrichments in various growth media: Growth medium 1 (BS1) contained (g L^−1^, pH 7.0) cellulose (2.0); tryptone (2.0); yeast extract (1.0); CaCl_2_·2H_2_O (1.0), NaCl (20.0), MgCl_2_·6H_2_O (3.6), MgSO_4_·7H_2_O (4.3), KCl (0.5), Na_2_S.9H_2_O (100 mg L^−1^). Growth medium 2 (BS2) was prepared by diluting the medium BS1 for 10 times, except for NaCl and Na_2_S.9H_2_O (Fig. 1C-i). Growth medium 3 (BS3) was prepared by adding sterilized Black Sea water (100 ml L^−1^) in BS2 and adjusting the sulfide concentration to 400 µM to mimic the natural composition of the Black Sea. Growth medium 4 (BS4) was prepared by replacing the cellulose by sodium propionate (0.2g L^−1^). Growth medium 5 (BS5) was prepared by replacing the cellulose in the BS2 medium by chitin (0.2g L^−1^) with 400 mg L^−1^ of Na_2_S.9H_2_O and BS2 culture extract (50 mL L^−1^). Growth medium 6, 7, 8 and 9 (BS6-9) were prepared by replacing the cellulose in the BS2 medium by cellobiose (0.2 g L^−1^), glucose (0.2 g L^−1^), pyruvate (0.2 g L^−1^) and acetate (0.2 g L^−1^), respectively. The amino acids mixture medium contained (L^−1^) Na_2_HPO_4_ (5.5 g); KH_2_PO_4_ (3.4 g); MgSO_4_.7H_2_O (0.2 g); CaCl_2_.2H_2_O (0.06 g); FeSO_4_.7H_2_O (0.5 mg); vitamin solution (2 ml); and trace elements (1 ml)^6^ with different amino acids (lysine (4.5 g), leucine (13 mg), isoleucine (13 mg), valine (17 mg), threonine (19 mg), methionine (14 mg), proline (11 mg), arginine (17 mg), histidine (20 mg), phenylalanine (16 mg), cysteine (12 mg) and tryptophan (4 mg). The medium based on hydrolyzed product of cellulose was obtained by growing the *Psychrilyobacter piezotolerans* strain S5 in growth medium BS1 at 25°C at atmospheric pressure. All medium components were sterilized by autoclaving at 121 °C for 15 min except, amino acids stocks which were filter sterilized (0.2 µm) before adding into the medium. MgCl_2_⋅6H_2_O and CaCl_2_⋅2H_2_O were added from separately autoclaved stocks to prevent precipitation. After autoclaving the growth media containing OM (yeast extract, tryptone and carbon sources) and sulfide, the media was stored at 20 °C for about 4 weeks to allow the maximal sulfurization of the OM as described earlier^56^. All substrates were obtained from Sigma-Aldrich (Merck KGaA, Darmstadt, Germany). Enrichments were carried out in 110-mL serum bottles containing 50 mL of medium. Anoxic conditions were created by flushing the growth medium with ultrapure nitrogen. Serum bottles were sealed with butyl rubber stoppers and aluminum caps. The headspace of the serum bottles was re-flushed with ultrapure nitrogen by using an Anaerobic Gas Exchange System (GR Instruments, Netherlands) to create strictly anaerobic conditions. Enrichments at elevated hydrostatic pressure (i.e., 20 MPa) were kept in a high-pressure cultivation device as described earlier^11^. Samples for microbial community analysis by 16S rRNA gene amplicon sequencing were taken from the enrichments after 13 days of incubations.

### CARD-FISH of enrichment cultures

The presence of various bacterial and archaeal members was assessed in the enrichments by catalyzed reporter deposition fluorescence in situ hybridization (CARD-FISH) with specific HRP-labeled probes (Table S7). Subsamples of 1 mL from the various enrichments (Fig. 1B, C-ii) were diluted 20 times and about 1 mL was filtered on 0.22 μM 24 mm diameter polycarbonate discs. The CARD-FISH protocol was performed as described earlier^20^. To avoid cell loss during cell wall permeabilization, filters were dipped in low-gelling-point agarose (0.2% [wt/vol] in MQ water, dried face up on glass slides at room temperature, and subsequently dehydrated in 96% (vol/vol) ethanol for 1 min. For cell wall permeabilization, filters were incubated in a lysozyme solution (10 mg mL^−1^ in 0.05 M EDTA, 0.1 M Tris-HCl [pH 7.5]) at 37°C for 30 min. The sections were washed with MQ water, dehydrated with 96% ethanol, dried at room temperature, and subsequently stored in petri dishes at −20°C until further processing. Filters were mounted on microscopic slides with mounting medium containing DAPI and analyzed on an Axio Imager.M2, Carl Zeiss Microscopy GmbH with 100x magnification.

### Genome and metagenome sequencing and assembly

Genomic DNA from pure cultures was extracted by the method of Marmur (1961)^57^. Sequencing of the genome of all the pure cultures (Table S6) were performed at the CGEB-IMR (Dalhousie University, Canada), using an Illumina MiSeq (Illumina Inc) platform. Illumina MiSeq reads of all the pure cultures were assembled with SPAdes - v3.15.3^58^. For metagenome sequencing, unamplified DNA extracts from the cellulose enrichments were used to prepare TruSeq nano libraries which were further sequenced with Illumina Miseq (5 samples multiplexed per lane) at the Utrecht Sequencing Facility, generating 45 million 2L×L250Lbp paired end reads. All the Illumina Miseq reads of enrichments were assembled by metaSPAdes - v3.15.3^59^.

### Scaffold binning and assessment of MAG quality

Scaffolds were binned into draft genome sequences based on coverage profile across samples and tetra-nucleotide frequencies with MetaBAT v.2.3.0^60^. The ‘superspecific’ preset was used to minimize contamination. To increase sensitivity without losing specificity, MetaBAT was run with ensemble binning, which aims to combine highly similar bins. Quality of the MAGs was assessed based on absence and presence of lineage-specific marker gene sets after genome placement in a reference tree with CheckM v1.0.8^61^.

### Genome and MAG annotations, and phylogenetic analysis

Annotation of the assembled data was performed with the Prokka pipeline^62^ available on the DOE systems biology knowledgebase (KBase)^63^, the Rapid Annotation Using Subsystem Technology (RAST; http://rast.nmpdr.org/rast.cgi)^64^ and *PATRIC* (Pathosystems Resource Integration Center; https://www.patricbrc.org/)^65^. The BlastKOALA tool^66^ was used for the functional characterization. CRISPR arrays were identified with CRT (v. 1.1)^67^. Average nucleotide identity (ANI)^68^ and digital DNA-DNA hybridization (DDH)^69^ was performed by using the Kostas lab server (http://enve-mics.ce.gatech.edu/ani/) and the genome-to-genome distance calculator (http://ggdc.dsmz.de/), respectively. To identify carbohydrate degradation-related enzymes, we used the online dbCAN2 meta server^70^ to annotate protein sequences by HMMER search against the CAZy database, including glycoside hydrolases (GHs), carbohydrate esterases (CEs), glycosyl transferases (GTs), carbohydrate-binding modules (CBMs), polysaccharide lyases (PLs), and auxiliary activities (AAs)^70^. Additionally, eggNOG annotations were used as auxiliary results for GH identification^71^. Pfam 31.0^72^ was used as the reference database for the annotation of peptidases, aminotransferases, and transporters of oligopeptides and amino acids by HMMER v3.1b2 (cut-off e value: 1e^−10^, best hits reserved). In addition, nonredundant peptidase unit sequences included in MEROPS (Release 12.1) were used for peptidase annotation by Diamond Blastp v0.8.36.98 ^73^. Potential subcellular localizations of putative CAZymes, peptidases, and lipases were predicted by uploading FASTA amino acid sequences of genes into the PSORTb 3.0.3 database^74^.

### Morphological, physiological, and biochemical analyses

Morphological properties and other physiological tests such as growth at different temperatures, pH and NaCl were performed as described earlier^6,75,76^. Glucose fermentation products were analyzed as described previously^76^. Growth on organic substrates (D-glucose, glycerol, cellulose, chitin, cellobiose, sucrose, starch, pyruvate, acetate, L-glutamate and L-aspartic acid and L-lysine) was tested in a minimal medium described previously^6,75,76^. Fermentation of various carbon sources (glucose, lactate, glycerol, xylose, galactose, cellobiose, maltose, lactose, aspartate, threonine, glutamate, and lysine) were tested as described previously^75,76^. Growth at elevated hydrostatic pressure (0.1 MPa-60 MPa) was tested in pressure vessels under strict anaerobic conditions as described previously^11^. Sulfide tolerance tests were performed in liquid medium which contained (g L^−1^, pH 7.0) pyruvate (2.0); tryptone (2.0); yeast extract (1.0); CaCl_2_·2H_2_O (1.0), NaCl (20.0), MgCl_2_·6H_2_O (3.6), MgSO_4_·7H_2_O (4.3) and KCl (0.5). To examine the sulfide tolerance of strains A1^T^ and A2, pyruvate was replaced with cellobiose, while the rest of the medium remained the same. The sulfide solution was neutralized before adding to the medium as described earlier^77^. Chemolithoautotrophic and chemolithoheterotrophic growth was tested as described earlier^78,79^. Various biochemical tests (cellulose/starch/casein/chitin hydrolysis, oxidase, and catalase activity) were carried out as described previously^6,76,80^. Other enzymatic activities and biochemical test were determined with API 20 A kit (bioMerieux, France) according to the instructions of the manufacturer.

### Core lipid and intact polar lipid (IPL) analyses

The core lipid of all the pure cultures isolated in this study were analyzed on the culture grown in liquid medium with pyruvate as carbon source except *Clostridiales* bacteria strains A1^T^ and A2. Strains A1^T^ and A2 were grown at their optimal conditions by using cellobiose as carbon source. Cells were harvested by centrifugation (5,000 g for 10 min at 20 °C) from late exponential phase of growth and core lipids were released by either base^81^ or acid hydrolysis^82^ and analyzed by gas chromatography/mass spectrometry (GC/MS), as described previously^81,82^. Intact polar lipids (IPLs) were extracted from freeze-dried biomass using a modified Bligh-Dyer procedure and were analyzed by Ultra High Pressure Liquid Chromatography-High Resolution Mass Spectrometry (UHPLC-HRMS) and by high performance liquid chromatography-ion trap mass spectrometry (HPLC-ITMS) as described previously^83^.

### Taxonomic description of novel taxa

Based on the distinct phenotypic and (meta) genomic properties, here, we propose the following novel taxa affiliated with the phyla *Bacillota*, *Spirochaetota*, *Bacteroidota*, *Desulfobacterota*, *Cloacimonadota*, *Mycoplasmatota*, *Ignavibacteriota*, *Planctomycetota* and *Chloroflexota*. The classification of isolates/enrichment cultures based on (meta) genomes were also assessed using GTDB-Tk and by constructing phylogenomic trees.

### Description of *Pontosulfidobacter* gen. nov

*Pontosulfidibacter* (Pon.to.sul.fi.di.bac’ter. Gr. n. *pontos euxeinos*, the Black Sea. N.L. neut. n. *sulfidum*, sulfide; N.L. masc. n. *bacter*, a bacterium; N.L. masc. n. *Pontosulfidibacter*, a bacterium from sulfidic waters of the Black Sea.

Two bacterial strains (i.e., A1^T^, A2), were isolated using cellobiose as the primary carbon source through repeated streaking on agar medium. Both strains shared 90.3% 16S rRNA gene sequence similarity to their closest cultured relative, *Fusibacter paucivorans* SEBR 4211^T^ ^84^, belonging to the order *Clostridiales* of the phylum *Bacillota.* Based on phylogenomic analysis, strains A1^T^ and A2 were found to be closely related to members of an uncultivated clade of the order *Clostridiales* (*Fusibacteraceae* MAG BB-3; Figs. 3 and S2), known to be important OM degraders in deep sulfidic marine habitats^4,5^. Both strains are strictly anaerobic, Gram-positive, spindle-shaped, endospore-forming bacteria (Fig. S2C-D; for further details, refer to the supplementary information). They form muddy whitish colonies on agar medium and release terminal endospores after 4 days of growth.

### Description of *Pontosulfidobacter cellobiosiphilus* sp. nov

*P. cellobiosiphilus* (cel.lo.bi.o.si′phi.lus. N.L. neut. n. *cellobiosum* cellobiose; Gr. masc. adj. *philos,* friend, loving; N.L. masc. adj. *cellobiosiphilus* loving cellobiose).

In addition to the properties given in the genus description, the cells are motile and have a spindle vibrioid to curved shape (0.8-0.9×3.0-5.0 µm). They stain Gram-positive and are negative for catalase, oxidase, and chitinase activity but positive for caseinase activity. Denitrification is not possible. The optimal growth occurs between pH 6.5-8.8 (with an optimum at 7.0-8.0; refer to Table S3). They tolerate up to 5.5 % NaCl, with the best growth at 2-3 %. The optimal growth temperature is 20-22 °C (with a range of 4-35 °C). While growth on D-Glucose, D-mannitol, D-lactose, and D-saccharose is possible, cellobiose is the preferred carbon source. However, growth is not possible with cellulose, D-melezitose, D-raffinose, D-sorbitol, L-rhamnose, and D-trehalose. Additional vitamins are not required for growth. Sulfate is not used as a terminal electron acceptor, whereas thiosulfate and sulfur serve as terminal electron acceptors. Genes of 2C-methyl-D-erythritol 4-phosphate (MEP) pathway of isopentenyl pyrophosphate (IPP) biosynthesis are present in the genome C_14:0_, C_16:1_ω9C, C_16:0_, C_18:2_, C_18:1_ω7C and C_18:0_ are present as major fatty acids (>5%) with minor amounts (<5 and >1%) of *iso*-C_15:0,_ C_16:1_ω5C and C_18:1_ω9C (Table S44). G+C content is 42.8 %. Type strain is A1^T^ (=KCTC 15661^T^ = JCM 32480^T^).

### Description of *Oceanispirochaeta piezotolerans* sp. nov

*O. piezotolerans* (pie.zo.to′le.rans. Gr. v. *piezo* to press; L. part. adj. *tolerans* tolerating, N.L. part. adj. *piezotolerans* tolerating high hydrostatic pressure).

Two bacterial strains affiliated with the phylum *Spirochaetota* (strains M1^T^ and M2 and S2) were isolated by repeated streaking on a glucose medium (BS6; see material and methods for details; Fig. 1C-i). Both strains shared 96.2% 16S rRNA gene sequence similarity with *Oceanispirochaeta litoralis* DSM 2029^T^ (Fig. S3A) and phylogenomic analyses showed affiliation with an uncultivated lineage of the phylum *Spirochaetota* (Figs. 3 and S3B) involved in the OM degradation in sulfidic marine habitats^4^. Both strains were strictly anaerobic, helical shaped bacteria (Fig. S3C-D) belonging to a novel species within the phylum *Spirochaetota* according to distinct phenotypic, physiological, and biochemical differences from their closest phylogenetic neighbors (see supplementary information and Table S1). The bacterium forms muddy whitish colonies on the agar medium. It appears as helical-shaped Gram-negative cells, measuring 0.1-0.2 µm in width and 5-20 µm in length. The cells exhibit helical movement and are highly motile. Sphaeroplasts are commonly observed under the microscope during the stationary growth phase (4-5 days). It grows on a pH range of 6.5 to 8.8, with the optimum pH between 7.0 and 8.0. The bacterium requires at least 0.5% NaCl for growth and can tolerate up to 5.5%. The optimal growth temperature is 18-23 °C, with a range of 4-28 °C. Under optimal conditions (20 °C, 2% NaCl, pH 7.5), the doubling time is 1.5-2.0 h. The bacterium exhibits negative reactions for catalase, indole production, and hydrolysis of cellulose, chitin, and gelatin. It requires yeast extract for growth but does not need additional vitamins. It is an obligate chemoorganoheterotrophic bacteria and thrives on glucose and pyruvate but not on cellobiose and cellulose. Under strict anaerobic conditions, it ferments glucose for growth. However, it does not utilize sulfate, sulfite, thiosulfate, nitrate, fumarate, or elemental sulfur as terminal electron acceptors. Dominant fatty acids (>5%) are: *iso*-C_13:0_, C_14:0_, *iso-*C_15:0_ and C_16:0_ (Table S45). Cardiolipins and glycolipids are the major polar lipid classes, mostly with mixed-ether/ester cores (Table S46). G+C content is 42.8%. Type strain is M1^T^ (JCM 32615^T^= KCTC 15719^T^).

### Description of ‘*Candidatus* Sphaerochaeta piezotolerans*’* sp. nov

*S. piezotolerans* (pie.zo.to′le.rans. Gr. v. *piezo* to press; L. part. adj. *tolerans* tolerating, N.L. part. adj. *piezotolerans* tolerating high hydrostatic pressure).

A bacterial strain (S2) was isolated by repeatedly streaking on glucose medium (BS6; see material and methods for details; Fig. 1C-i). It belongs to the genus *Sphaerochaeta* in the phylum *Sphaerochaeta* and showed 99.1% 16S rRNA gene sequence similarity to *Sphaerochaeta halotolerans* 4-11^T^ (Fig. S3A). However, when compared to *S. halotolerans* 4-11^T^, its average nucleotide identity (ANI) and digital DNA-DNA hybridization (DDH) were found to be below the species cut-off (Table S2). Physiological (Table S3) and phylogenomic analyses (Fig. S3B) confirmed that strain S2 belongs to a novel species affiliated with the phylum *Spirochaetota* (see supplementary information). The cells of this culture were spherical in shape and stained Gram-negative (Fig. S3E). Due to the loss of viability during repeated subculturing in our laboratory conditions, it is referred to as candidatus status to describe this species. Cells of strain S2 are spherical shaped (0.8-0.9 μm in diameter) and Gram negative. Growth occurs in media with pH 6.5 to 8.8, and the optimum pH ranges between pH 7.0 and 8.0. They require at least 0.5 % NaCl for growth and could tolerate up to 5.5 %. Optimum growth occurs at 18-23 °C, with a range for growth of 4-28 °C. The doubling time under optimal conditions [20 °C, 2 % (w/v) NaCl, pH 7.5] are 1.5-2.0 h. Shows negative reactions for catalase, indole production and cellulose, chitin, and gelatin hydrolysis. Growth is possible from 0.1-50 MPa of hydrostatic pressure. Chemoorganoheterotrophic growth is possible. Sodium thioglycolate is required as reducing agent for growth. Growth is possible on glucose, pyruvate, and acetate. However, xylan, chitin and cellulose are not utilized for growth. Grow under strict anaerobic conditions by fermenting glucose. Sulfate, sulfite, thiosulfate, and nitrate are not utilized as terminal electron acceptor. The genomic G+C content is 46.87%.

### Description of *Lutibacter atrimaris* sp. nov

*L. atrimaris* (at.ri.ma’ris. L. adj. ater, black; mare, -is, sea; N.L. gen. n. *atrimaris*, of the Black Sea).

Two bacterial strains (B1^T^ and B2) were isolated on a pyruvate-containing medium (BS8; see M&M for details; Fig. 1C-i). They share 96.3% 16S rRNA gene sequence similarity with *Lutibacter profundi* LP1^T^ (Fig. S4A) and belong to an uncultivated clade (phylotype 10972)^7^ (*Lutibacter* ASV_02820)^5^ of the order *Flavobacteriales*, within the phylum *Bacteroidota*. These taxa play a significant role in OM degradation in deep sulfidic marine environments^4,5^ (see Fig. S5A). Both strains display motile and small rod-shaped cells (0.2-0.3×3.0-5.0 µm) that stain Gram-negative and are facultative anaerobic bacteria. They are negative for catalase, oxidase, and chitinase activity but positive for cellulase and amylase activity. Additionally, they produce indole but do not perform denitrification. The growth occurs between pH 6.5-9.0 (optimum 7.0-8.0), and they tolerate up to 5.5% NaCl, with optimum growth at 2-3%. The optimal growth temperature is 20-22 °C (range 4-35 °C). Their growth mode is chemoorganoheterotrophic. For growth, they utilize D-Glucose, D-mannitol, D-lactose, D-saccharose, D-maltose, salicin, D-xylose, L-arabinose, cellobiose, sucrose, and starch. However, they cannot grow with cellulose, D-melezitose, D-raffinose, D-sorbitol, L-rhamnose, and D-trehalose. The genome contains genes of the 2C-methyl-D-erythritol 4-phosphate (MEP) pathway and the mevalonate pathway of IPP biosynthesis *Iso*-C_15:0_, *iso*-C_15:1_ω11C, *iso*-β-OH C_15:0_ (Table S49) and glycine *iso*-β-OH C17:0 amide were present as major fatty acids (>5%) in strains B1^T^ and B2 along with minor amounts (<5 and >1%) of *iso-*C_13:0,_ *anteiso*-C_15:1_ω11C, *anteiso*-C_15:0_, C_15:0_, *iso*-C_17:0_, *iso*-α-OH C_15:0_, *iso*-β-OH C_17:0_, glycine *iso*-β-OH C_15:0_ amide, glycine *iso*-β-OH C_16:0_ amide, glycine β-OH *n-* C_16:0_ amide and glycine *anteiso*-β-OH C_17:0_ amide. MK-6 is the predominant menaquinone. Phosphatidylethanolamines, ornithines, flavolipins, capnines and glycine lipids are the major polar lipid groups with minor amounts (<3.5%) of glutamine lipids and phosphatidylcholines^81^. G+C content is 30.2 %. Type strain is B1^T^ (=KCTC 62862^T^ = JCM 32877^T^).

### Description of ‘*Candidatus* Bradybacterium’ gen. nov

*Bradyacterium* (Bra.dy.bac.te’ri.um. Gr. adj. *bradys*, slow; N.L. neut. n. *bacterium*, a bacterium; N.L. neut. n. *Bradybacterium*, a slow-growing bacterium).

Two bacterial strains, S6 and L6, were isolated using cellobiose as the primary carbon source through repeated streaking on agar medium. Both strains exhibited 88.7% 16S rRNA gene sequence similarity to *Lentimicrobium saccharophilum* TBC1^T^ (Fig. S4A). Phylogenomic analysis revealed their relation to an uncultivated lineage (phylotype 5079 of the family *Lentimicrobiaceae*) within the order *Bacteroidales* of the phylum *Bacteroidota* (Figs. 3 and S4B), known for its involvement in cyanobacterial necromass degradation in sulfidic marine habitats^7^. These strains were negative for catalase, oxidase, and chitinase activity but positive for cellulase and amylase activity. They possess a chemoorganoheterotrophic growth mode. The genome of both strains contains genes of the 2C-methyl-D-erythritol 4-phosphate (MEP) and mevalonate pathway of IPP biosynthesis.

### Description of ‘*Candidatus* Bradybacterium flavum’ sp. nov

*B. flavum* (fla’vum. L. neut. adj. *flavum*, yellow, referring to the colony colour).

In addition to the properties given in the genus description, this bacterium displays the following characteristics: it appears as motile and small rods measuring 0.2-0.3×3.0-5.0 µm. Visible colonies on agar medium take approximately 15-21 days to form. The cells stain Gram-negative and are strict anaerobes. They do not exhibit catalase, oxidase, or chitinase activity, but they show positive amylase and caseinase activity. Growth occurs within a pH range of 6.5-8.8, with the optimal pH being 7.0-8.0. It tolerates up to 5.5% NaCl, with the best growth observed at 2-3%. The optimum growth temperature is between 20-22 °C, and it can tolerate temperatures ranging from 4-30 °C. Chemoorganoheterotrophy is the sole growth mode. For growth, it utilizes D-glucose, cellobiose, sucrose, starch, and D-mannose. However, it cannot grow with cellulose, D-melezitose, D-raffinose, and D-trehalose. The genomic G+C content is 35.0%.

### Description of *Pseudodesulfovibrio turroides* sp. nov

*P. turroides* (tur.ro.i’des. L. n. *turris*, tower; L. suff. -*oides*, looking like; N.L. masc. adj. *turroides*, looking like a tower, referring to the unusual shape of the colonies).

Two bacterial strains (i.e., S3^T^, and S3-i) were isolated under strict anaerobic conditions by repeatedly streaking on a pyruvate containing medium (BS8; see M&M for details; Fig. 1C-i). Phylogenetic analysis of strains S3^T^ and S3-i (Fig. 1B) indicated their close affiliation with an uncultivated clade of the family *Desulfovibrionaceae* (Fig. S6A-B) while they shared 97.4% 16S rRNA gene sequence similarity with *Pseudodesulfovibrio indicus* J2^T^. Strains S3^T^ and S3-i formed orange colored and tower-shaped colonies on agar medium (after one month of incubation) when incubated at 18-25°C under strict anaerobic conditions (Fig. S6C-E). The orange colonies are quite surprising since there no orange-pigmented members of the family *Desulfovibrionaceae* have previously been reported. Moreover, to the best of our knowledge, there is no reported bacteria forming tower-shaped colonies. Nevertheless, cells of both strains were of typical vibrio shape and stained Gram-negative (Fig. S6C-E). Comparative analysis of both strains with their nearest phylogenetic neighbors (Table S5) indicated they belong to a novel species within the family *Desulfovibrionaceae* of the phylum *Desulfobacterota* (see supplementary information). The cells can tolerate short (30 min) exposure to oxygen. They are motile, curved rods, measuring 0.7-0.9 µm in width and 2.0-5.0 µm in length. They test negative for catalase and oxidase. Growth occurs under strictly anaerobic conditions at pH 6.5-8.8 (optimum 7.5-8.0). At least 0.5% (w/v) NaCl is necessary for growth, and up to 5.5% is tolerated. The optimum growth temperature is 18-23 °C, with a range of 4-30 °C. Pyruvate, acetate, L-Glutamate, aspartate, and fumarate are utilized for growth. Although chemolithoheterotrophic growth is possible, they preferably grow chemoheterotrophically. Sulfate serves as the terminal electron acceptor, and hydrogen is utilized as an energy source. Traces of yeast extract are required for growth, but added vitamins are not essential. Genes of 2C-methyl-D-erythritol 4-phosphate (MEP) pathway are present in the genome. *iso*-C_15:0_, *anteiso*-C_15:0_, C_16:0_, *iso*-C_17:0_, *iso*-C_17:1_ω8t and C_18:0_ are present as major fatty acids (>5%) with minor amounts (<5 and >1%%) of *iso*-C_16:1_ω7t, *iso*-C_16:0_, *iso*-C_17:1_ω8, *anteiso*-C_17:1_ω8, *anteiso*-C_17:0_, *iso*-C_19:1_ω8t, *iso*-C_19:0_ (Table S48). The genomic G+C content is 56.6 %. Type strain is S3^T^ (=KCTC 15635^T^ = JCM 32483^T^).

### Description of ‘*Candidatus* Atrimarinoplasma cellobiosiphila’

Atrimarinoplasma (Atri.ma.ri.no.plas’ma. N.L. n. atrum, black; N.L. n. mare, sea; N.L. neut. n. plasma, something formed; referring to a black marine bacterium).

cellobiosiphila (cel.lo.bi.o.si′phi.la. N.L. neut. n. *cellobiosum* cellobiose; N.L. fem. adj. *phila* from Gr. fem. adj. phileLJ friendly to, loving; N.L. fem. adj. *cellobiosiphila* loving cellobiose).

A cellobiose-medium amended with the antibiotic rifampicin (100 µg mL^−1^) was used to enrich strain Izemo-BS, a new member of the phylum *Mycoplasmatota*, and the enrichment level was found to be 22.2% based on 16S rRNA gene amplicon sequencing (Fig. 1A). The enrichment of this strain with rifampicin was not suspected as this antibiotic is mostly used for the enrichment of spirochetes due to their resistance to it^85^. Although the strain could not be isolated, an almost complete MAG (97.3% completeness with 1.3% contamination) was recovered and revealed that Izemo-BS is a new taxon at the family level of the *Mycoplasmatota* phylum (Fig. S10A-B; see supplementary information). 16S rRNA gene sequence (380 bp) of Izemo-BS shared only 85.59% similarity with *Acholeplasma morum* ATCC 33211^T^. It showed highest similarity (<84) with several uncultured 16S rRNA gene sequence of the members of phylum *Mycoplasmatota*. Within the MAG, a variety of transporters involved in the passage of phosphate, D-methionine, Fe^2+^, manganese, chromate, magnesium, cadmium, zinc, cobalt, biotin, maltose, lactose, L-arabinose, aspartate, glutamate, arginine, glutamine, vitamin B_12_, oligopeptides, and dipeptides are present. A variety of genes involved in nucleic acid degradation are also present. Cellobiose could be another important source for growth in addition to nucleic acids. resistant to rifampicin. The G+C content of the MAG is 31.1%.

### Description of ‘*Candidatus* Atrimarinobacter sulfidophilus’

Atrimarinobacter (Atri.ma.ri.no.bac’ter. N.L. n. atrum, black; N.L. n. mare, sea; N.L. masc. n. bacter; referring to a black marine bacterium).

sulfidophilus (sul.fi.di.phi′lus. N.L. n. *sulfidum* sulfide; N.L. masc. adj. *philus* from Gr. adj. *philos* loving; N.L. masc. adj. *sulfidiphilus* sulfide-loving).

Strain Cloa-SY6, which belongs to the *Cloacimonadota* phylum, was enriched in a medium with 0.2 g L^−1^ of propionate (BS4; see M & M for details; Fig. 2) from primary enrichments (cellulose with 4 mM of sulfide; Fig. 1B). An almost complete MAG (>98% completeness, contamination 1.1; see Table S6) was obtained, and the 16S rRNA gene sequence contained in the MAG revealed that it had only 83-84% 16S rRNA gene sequence similarity with its nearest relatives, *’Ca* Cloacamonas acidaminovorans’ and *’Ca* Syntrophosphaera thermopropionivorans’ (see Figs. 3 and S8A-B). A CARD-FISH probe was specifically designed to target the 16S rRNA sequence (see Table S7), which revealed that the level of enrichment was approximately 17%, and that the cell morphology of the strain was spherical to oval and had a diameter of 0.9-1.0 μm (see Figs. 2, S8C-D). When compared to closely related species, strain Cloa-SY6, it exhibited significant differences with respect to phylogenetic (Fig. S8A), phylogenomic (Fig. S8B), and physiological (Table S8) properties and was determined to be a new taxon at the class level of the phylum *Cloacimonadota* (see Fig. S8; see supplementary information).

16S rRNA gene sequence (1535 bp) of Cloa-SY6 shares 83.8% similarity with ‘*Candidatus* Syntrophosphaera thermopropionivorans’ of the phylum *Cloacimonadota*. The bacterium grows as oval to spherical shape with a diameter of 0.8-0.9 µm that may form chains. Sodium propionate is expected to be preferred carbon source for growth. Cells can tolerate high sulfide concentration (>2 mM) and are oligotrophic and psychrophilic. Within the MAG, a variety of transporters involved in the passage of phosphate, D-methionine, Fe^2+^, manganese, chromate, magnesium, cadmium, zinc, cobalt, biotin, riboflavin, fatty acids, dicarboxylic acids, lactate, glutamine, and vitamin B_12_ were detected. All the genes involved in the 2C-methyl-D-erythritol 4-phosphate (MEP) pathway of isoprenoid biosynthesis are present. The TCA cycle is incomplete with the steps from 2-oxoglutarate to fumarate and the conversion of malate to oxaloacetate missing. The G+C content of the MAG is 34.3.

### Description of ‘*Candidatus* Ponteuxinibacter sulfidophilus’

Ponteuxinibacter (Pont.eu.xi.ni.bac’ter. L. masc. n. and adj. Pontus Euxinus, Black Sea; L. masc. n. bacter, rod; N.L. masc. n. Ponteuxinibacter, rod-shaped bacterium from the Black Sea).

sulfidophilus (sul.fi.di.phi′lus. N.L. n. *sulfidum* sulfide; N.L. masc. adj. *philus* from Gr. adj. *philos* loving; N.L. masc. adj. *sulfidiphilus* sulfide-loving).

In the enrichment with 4 mM sulfide (Fig. 1A), the microbial community showed the presence of Igna-SY6, which was found to make up to 8% of the community, as confirmed by 16S rRNA gene amplicon analysis (Fig. 2). Using a specially designed CARD-FISH probe (Table S7), it was determined that the cell morphology of Igna-SY6 was rod-shaped, measuring 5-6 x 0.8-0.9 μm (Fig. S9C-D). The high-quality MAG obtained (>98% completeness; Table S6) indicated that Igna-SY6 was affiliated with the family *Ignavibacteriaceae* of the *Ignavibacteriota* phylum and was classified as a novel taxon (Fig. S9A-B; see supplementary information). 16S rRNA gene sequence of Igna-SY6 shares 88.0% (407 bp) similarity with *Ignavibacterium album* JCM 16511^T^ of the family *Ignavibacteriaceae* of the phylum *Ignavibacteriota*. The cell morphology of the bacterium is rod shaped with 4-5 μm in length and 0.5-0.6 μm in width (Table S14). Genome indicates the utilization of starch, cellulose, cellobiose, glucose and pyruvate. Cells can tolerate high sulfide concentration (>2 mM) and are oligotrophic and psychrophilic. Within the MAG, a variety of transporters involved in the passage of phosphate, Fe^2+^, manganese, chromate, magnesium, cadmium, zinc, cobalt, bicarbonate, biotin, glucose, hexuronate, L-arabinose, glutamate, putrescine, vitamin B_12_, and dipeptides were detected. The MAG contains all the genes involved in the 2C-methyl-D-erythritol 4-phosphate (MEP) pathway of isoprenoid biosynthesis. Transporter systems for phosphate, aspartate/glutamate/glutamine, and cystine are present. The G+C content of the MAG is 32.9.

### Description of ‘*Candidatus* Atrisphaera chitinolytica’

Atrisphaera (A.tri.sphae’ra. L. masc. adj. ater, black, dark; L. fem. n. sphaera, sphere; N.L. fem. n. Atrisphaera, a spherical-shaped bacterium from the Black Sea).

chitinolytica (chi.tin.o.lyt’i.ca. chem. term chitin, chitin, a polysaccharide; Gr. adj. lytos, soluble; M. L. fem, adj. chitinolytica, dissolving chitin).

Incubation using chitin in a diluted (0.2 g L^−1^) growth medium (BS5; see M&M for details) at 10 °C resulted in enrichment of a new member of the *Planctomycetota* phylum, named Plnct-SY6 (Fig. 2). A high degree of enrichment (40-50%) and a spherical-to-oval shaped cell morphology with a diameter of 1.0-1.2 μm was assessed by using CARD-FISH with a specific probe based on the 16S rRNA gene sequence (Table S7; Fig. S7C-D). Although isolation of Plnct-SY6 could not be achieved, a high-quality metagenome assembled genome (MAG) was recovered (Table S6), confirming that the isolate Plnct-SY6 represents a new taxon at the family level of the *Planctomycetota* phylum (shown in Fig. S7A-B; see supplementary information). Plnct-SY6 shared 91.25% (413 bp) 16S rRNA gene sequence similarity with *Limihaloglobus sulfuriphilus* SM-Chi-D1^T^ of the family *Sedimentisphaeraceae* of the phylum *Planctomycetota*. Cells are oval to spherical shape with a diameter of 0.8-0.9 µm. It was found to be positive for chitinolytic activity, and the genome sequence indicated its ability to utilize starch, cellulose, cellobiose, glucose, and pyruvate. Plnct-SY6 can tolerate high sulfide concentration (>2 mM) and is characterized as oligotrophic and psychrophilic. Within the MAG, a variety of transporters involved in the passage of phosphate, D-methionine, Fe^2+^, manganese, chromate, bicarbonate, magnesium, cadmium, zinc, cobalt, biotin, glucose, trehalose, L-lactate, putrescine, dicarboxylic acids, vitamin B_12_, oligopeptides, and dipeptides were detected. The MAG contains all the genes involved in the 2C-methyl-D-erythritol 4-phosphate (MEP) pathway of isoprenoid biosynthesis. The G+C content of the MAG is 41.3.

### Description of ‘*Candidatus* Euxinilinea sulfidophila’

Euxinilinea (Eu.xi.ni.li’ne.a. N.L. fem. n. Euxinus, pertaining to the Black Sea; L. fem. n. linea, a line, thread; referring to a bacterium isolated from the Black Sea, forming thread-like structures). sulfidophila (sul. fi. do’phi, la. Gr. adj. philus loving; M.L. adj. sulfidophila sulfide-loving.

A member (Chflx-SY6) of the phylum *Chloroflexota* was identified in the enrichments containing 4 mM of sulfide (Fig. 1B), albeit with a low relative abundance of 1% as by using 16S rRNA gene amplicon and CARD-FISH analyses. We were unable to increase its abundance in further enrichment experiments. Nonetheless, a high-quality MAG of Chflx-SY6 with an almost complete 16S rRNA gene sequence (1,509 bp) was recovered. The 16S rRNA gene sequence had a similarity of only 91.2% with its closest cultured relative, *Ornatilinea apprima* P3M-1^T^ ^86^, and <90.8% similarity with other cultivated members of the *Chloroflexota* phylum. (Fig. S9A). A comparative genomic analysis with its nearest phylogenetic neighbor revealed that Chflx-SY6 represents probably a novel genus within the *Chloroflexota* phylum (Table S9, and Fig. S9A-C; see supplementary information). Cells are rod shape with a length of 4.0-5.0 µm. Within the MAG, a variety of transporters involved in the passage of phosphate, D-methionine, Fe^2+^, manganese, magnesium, cadmium, nickel, calcium, potassium, L-cystine, zinc, cobalt, sulfate, biotin, maltose, L-lactate, L-arabinose, sialic acid, spermidine, putrescine, arginine, glutamine, vitamin B_12_, oligopeptides, and dipeptides were detected. The MAG contains all the genes involved in the mevalonate pathway of isoprenoid biosynthesis. Its genome also contains the complete gene repertoire of the Wood-Lunjdahl pathway (Fig. 4; Fig. S22), supporting its potential role in CO_2_ fixation in the deep sulfidic waters of the Black Sea. G+C content of the MAG is 43.1.

## Data Availability

16S rRNA gene amplicon sequencing data of the Black Sea water columns are deposited in NCBI as mentioned earlier^6^. 16S rRNA gene amplicon data of enrichment cultures are deposited under the NCBI bioproject accessions PRJNA681484, PRJNA596220 and PRJNA602617.

## Acknowledgements

The authors are thankful to Marianne Baas for helping in fermentation product (gas phase) analysis, to Sanne Vreugdenhil and Maartje Brouwer for their support with the molecular genetic analyses. Dr. Garret Smith (Radboud University, Nijmegen, The Netherlands) is greatly acknowledged for helping in phylogenomic analyses. We would like to thank the cruise leader, captain, crew, and scientific participants of the Black Sea expedition-2017 (64PE418) and 2018 on board of the *R/V* Pelagia for sampling and technical support. We thank Prof. Bernhard Schink (University in Konstanz, Germany) and Dr. Daan van Vliet (Wageningen University and Research) for their expert suggestions concerning the correct species epithet and Latin etymology. Johan van Heerwaarden, Edwin Keijzer and Roel Bakker (NMF, NIOZ) are greatly acknowledged for devising and helping with the “high pressure anaerobic cultivation” assembly. We acknowledge the Utrecht Sequencing Facility (USEQ) for providing sequencing service and data. USEQ is subsidized by the University Medical Center Utrecht and The Netherlands X-omics Initiative (NWO project 184.034.019).

## Author Contributions

SY, LV and JSSD designed the study. SY performed most of the laboratory work and data analysis; MK and NB performed the fatty acid and intact polar lipid analysis; JE, and WR sorted the protein sequences of nucleases, peptidases, and lipases; SY, LV and JSSD wrote the manuscript. All authors read and approved the final version of the manuscript.

## References

1. Karl, D. M. Microbial oceanography: paradigms, processes and promise. Nat. Rev. Microbiol. 5, 759–769 (2007).

2. Middelburg, J. J., Vlug, T., Jaco, F. & Nat, W. A. Organic matter mineralization in marine systems. Glob. Planet. Change. 8, 47–58 (1993).

3. Arndt, S. et al. Quantifying the degradation of organic matter in marine sediments: A review and synthesis. Earth-Sci. Rev. 123, 53–86 (2013).

4. Pelikan, C. et al. Anaerobic bacterial degradation of protein and lipid macromolecules in subarctic marine sediment. ISME J. 15, 833–847 (2021).

5. Wasmund, K. et al. Genomic insights into diverse bacterial taxa that degrade extracellular DNA in marine sediments. Nat. Microbiol. 6, 885–898 (2021).

6. Yadav, S., Koenen, M., Bale, N., Sinninghe Damsté, J. S. & Villanueva, L. The physiology and metabolic properties of a novel, low-abundance *Psychrilyobacter* species isolated from the anoxic Black Sea shed light on its ecological role. Env. Microbiol. Rep. 13, 899–910 (2021).

7. Müller, A. L. et al. Bacterial interactions during sequential degradation of cyanobacterial necromass in a sulfidic arctic marine sediment. Env. Microbiol. 20, 2927–2940 (2018).

8. Zheng, R. et al. Characterization of the first cultured free-living representative of Candidatus Izemoplasma uncovers its unique biology. ISME. J. 15, 2676–2691 (2021).

9. Zheng, R., Wang, C., Cai, R., Shan, Y. & Sun, C. Mechanisms of nucleic acid degradation and high hydrostatic pressure tolerance of a novel deep-sea wall-less bacterium. mBio (2023).

10. Skennerton, C. T. et al. Phylogenomic analysis of Candidatus ‘Izimaplasma’ species: free-living representatives from a Tenericutes clade found in methane seeps. ISME J. 10, 2679–2692 (2016).

11. Yadav, S. et al. Physiological, chemotaxonomic and genomic characterization of two novel piezotolerant bacteria of the family Marinifilaceae isolated from sulfidic waters of the Black Sea. Syst. Appl. Microbiol. 43, (2020).

12. Li, J., Dong, C., Lai, Q., Wang, G. & Shao, Z. Frequent Occurrence and Metabolic Versatility of Marinifilaceae Bacteria as Key Players in Organic Matter Mineralization in Global Deep Seas. mSystems 7:e0086422, (2022).

13. Suominen, S., Dombrowski, N., Sinninghe Damsté, J. S. & Villanueva, L. A diverse uncultivated microbial community is responsible for organic matter degradation in the Black Sea sulfidic zone. Env. Microbiol. 23, 2709–2728 (2021).

14. Suominen, S., Doorenspleet, K., Sinninghe Damsté, J. S. & Villanueva, L. Microbial community development on model particles in the deep sulfidic waters of the Black Sea. Env. Microbiol. 23, 2729–2746 (2021).

15. Suominen, S., Gomez-Saez, G. V., Dittmar, T., Sinninghe Damsté, J. S. & Villanueva, L. Interplay between microbial community composition and chemodiversity of dissolved organic matter throughout the Black Sea water column redox gradient. Limnol. Ocean. 67, 329–347 (2022).

16. Volkov, N., II & L.N. Hydrogen Sulfide in the Black Sea. 309–331 (2007).

17. Ducklow, H., Hansell, D. & Morgan, J. Dissolved organic carbon and nitrogen in the Western Black Sea. Mar. Chem. 105, 140–150 (2007).

18. Margolin, A., Gonnelli, M., Hansell, D. & Santinelli, C. Black Sea dissolved organic matter dynamics: Insights from optical analyses. Limnol. Ocean. 63, 1425–1443 (2018).

19. Hansell, D. A., Carlson, C. A., Repeta, D. J. & Schlitzer, R. Dissolved organic matter in the ocean: A controversy stimulates new insights. Oceanography 22, 202–211 (2009).

20. Villanueva, L. et al. Bridging the membrane lipid divide: bacteria of the FCB group superphylum have the potential to synthesize archaeal ether lipids. ISME J. 15, 168–182 (2021).

21. Pelletier, E. et al. Candidatus Cloacamonas acidaminovorans”: genome sequence reconstruction provides a first glimpse of a new bacterial division. J. Bacteriol. 190, 2572–2579 (2008).

22. Rotaru, A. E., Schauer, R., Probian, C., Mussmann, M. & Harder, J. Visualization of candidate division OP3 cocci in limonene-degrading methanogenic cultures. J. Microbiol. Biotechnol. 22, 457–461 (2012).

23. Hug, L. A. et al. Overview of organohalide-respiring bacteria and a proposal for a classification system for reductive dehalogenases. Philos. Trans. R. Soc. Lond. B Biol. Sci. 368, (2013).

24. Jochum, L. M. et al. Single-cell genomics reveals a diverse metabolic potential of uncultivated Desulfatiglans-related Deltaproteobacteria widely distributed in marine sediment. Front. Microbiol. 9, 1–16 (2018).

25. Hawley, A. K. et al. Diverse Marinimicrobia bacteria may mediate coupled biogeochemical cycles along eco-thermodynamic gradients. Nat. Commun. 8, (2017).

26. Dyksma, S. & Gallert, C. Candidatus Syntrophosphaera thermopropionivorans: a novel player in syntrophic propionate oxidation during anaerobic digestion. Env. Microbiol. Rep. 11, 558–570 (2019).

27. Ravin, N. V. et al. Genome Analysis of Fimbriiglobus ruber SP5T, a Planctomycete with Confirmed Chitinolytic Capability. Appl. Env. Microbiol. 84:e02645–17, (2018).

28. Rinke, C. et al. Insights into the phylogeny and coding potential of microbial dark matter. Nature 499, 431–437 (2013).

29. Johnson, L. A. & Hug, L. A. Cloacimonadota metabolisms include adaptations in engineered environments that are reflected in the evolutionary history of the phylum. Env. Microbiol. Rep. 14, 520–529 (2022).

30. Drickamer, K. & Taylor, M. E. Evolving views of protein glycosylation. Trends Biochem. Sci. 23, 321–324 (1998).

31. Bei, Q., Peng, J. & Liesack, W. Shedding light on the functional role of the Ignavibacteria in Italian rice field soil: A meta-genomic/transcriptomic analysis. Soil Biol. Biochem. 163, (2021).

32. Mazur, O. & Zimmer, J. Apo- and cellopentaose-bound structures of the bacterial cellulose synthase subunit BcsZ. J. Biol. Chem. 286, 17601–17606 (2011).

33. Ahmad, I. et al. BcsZ inhibits biofilm phenotypes and promotes virulence by blocking cellulose production in Salmonella enterica serovar Typhimurium. Microb. Cell Fact. 15, (2016).

34. Tomazini, A. Jr., et al. Analysis of carbohydrate-active enzymes in Thermogemmatispora sp. strain T81 reveals carbohydrate degradation ability. Can. J. Microbiol. 64, 992–1003 (2018).

35. Pradel, N. et al. The first genomic and proteomic characterization of a deep-sea sulfate reducer: insights into the piezophilic lifestyle of Desulfovibrio piezophilus. PLoS One 8:e55130, (2013).

36. Qin, Q. L. et al. Oxidation of trimethylamine to trimethylamine N-oxide facilitates high hydrostatic pressure tolerance in a generalist bacterial lineage. Sci. Adv. 7:eabf9941, (2021).

37. Allen, E. E., Facciotti, D. & Bartlett, D. H. Monounsaturated but not polyunsaturated fatty acids are required for growth of the deep-sea bacterium Photobacterium profundum SS9 at high pressure and low temperature. Appl. Env. Microbiol. 65, 1710–1720 (1999).

38. Grossi, V. et al. Hydrostatic pressure affects membrane and storage lipid compositions of the piezotolerant hydrocarbon-degrading Marinobacter hydrocarbonoclasticus strain #5. Env. Microbiol. 12, 2020–2033 (2010).

39. Amrani, A. et al. Transcriptomics reveal several gene expression patterns in the piezophile Desulfovibrio hydrothermalis in response to hydrostatic pressure. PLoS ONE 9:e106831, (2014).

40. Ono, H. et al. Characterization of biosynthetic enzymes for ectoine as a compatible solute in a moderately halophilic eubacterium, Halomonas elongata. J. Bacteriol. 181, 91–99 (1999).

41. Pastor, J. M. et al. Ectoines in cell stress protection: uses and biotechnological production. Biotechnol. Adv. 28, 782–801 (2010).

42. Leloup, J. et al. Diversity and abundance of sulfate-reducing microorganisms in the sulfate and methane zones of a marine sediment, Black Sea. Env. Microbiol. 9, 131–142 (2007).

43. Koch, T. & Dahl, C. A novel bacterial sulfur oxidation pathway provides a new link between the cycles of organic and inorganic sulfur compounds. ISME J. 12, 2479–2491 (2018).

44. Hildebrandt, T. M. & Grieshaber, M. K. Three enzymatic activities catalyze the oxidation of sulfide to thiosulfate in mammalian and invertebrate mitochondria. FEBS J. 275, 3352–61 (2008).

45. Shahak, Y. & Hauska, G. Sulfide oxidation from cyanobacteria to humans: sulfide-quinone oxidoreductase (SQR. in Sulfur Metabolism in Phototrophic Organisms (eds. Dahl, C., Knaff, D. & Leustek, T.) vol. 27 319–335 (Springer, 2008).

46. Dahl, C. Sulfur metabolism in phototrophic bacteria. in Modern Topics in the Phototrophic Prokaryotes (ed. Hallenbeck, P.) (Springer, 2017).

47. Xia, Y. et al. Sulfide production and oxidation by heterotrophic bacteria under aerobic conditions. ISME J. 11, 2754–2766 (2017).

48. Strokal, M. & Kroeze, C. Nitrogen and phosphorus inputs to the Black Sea in 1970-2050. Reg Env. Change 13, 179–192 (2013).

49. Panin, N. & Jipa, D. Danube river sediment input and its interaction with the northwestern Black Sea. Estuar. Coast Shelf Sci. 54, 551–562 (2002).

50. Lechner, A. et al. The Danube so colourful: a potpourri of plastic litter outnumbers fish larvae in Europe’s second largest river. Env. Pollut. 188, 177–181 (2014).

51. Lauro, F. M. et al. The genomic basis of trophic strategy in marine bacteria. Proc. Natl. Acad. Sci. USA. 106, 15527–15533 (2009).

52. Asbun, A. A. et al. Cascabel: A Scalable and Versatile Amplicon Sequence Data Analysis Pipeline Delivering Reproducible and Documented Results. Front. Genet. 11, (2020).

53. Andrews, S. FastQC: a quality control tool for high throughput sequence data [Internet. (2010).

54. Zhang, J., Kobert, K., Flouri, T. & Stamatakis, A. PEAR: a fast and accurate Illumina Paired-End reAdmergeR. Bioinformatics 30, 614–620 (2014).

55. Caporaso, J. G. et al. QIIME allows analysis of high-throughput community sequencing data. Nat. Methods 7, 335–336 (2010).

56. Pohlabeln, A. M., Gomez-Saez, G. V., Noriega-Ortega, B. E. & Dittmar, T. Experimental Evidence for Abiotic Sulfurization of Marine Dissolved Organic Matter. Front. Mar. Sci. 4, (2017).

57. Marmur, J. A. Procedure for the isolation of deoxyribonucleic acid from microorganisms. J. Mol. Biol. 3, 208–218 (1961).

58. Bankevich, A. et al. SPAdes: a new genome assembly algorithm and its applications to single-cell sequencing. J. Comput. Biol. 19, 455–477 (2012).

59. Nurk, S., Meleshko, D., Korobeynikov, A. & Pevzner, P. A. metaSPAdes: a new versatile metagenomic assembler. Genome Res. 27, 824–834 (2017).

60. Kang, D. D., Froula, J., Egan, R. & Wang, Z. MetaBAT, an efficient tool for accurately reconstructing single genomes from complex microbial communities. PeerJ. 3:e1165, (2015).

61. Parks, D. H., Imelfort, M., Skennerton, C. T., Hugenholtz, P. & Tyson, G. W. CheckM: assessing the quality of microbial genomes recovered from isolates, single cells, and metagenomes. Genome Res. 25, 1043–1055 (2015).

62. Seemann, T. Prokka: rapid prokaryotic genome annotation. Bioinformatics 30, 2068–2069 (2014).

63. Arkin, A. P. et al. KBase: The United States Department of Energy Systems Biology Knowledgebase. Nat. Biotechnol. 36, (2018).

64. Brettin, T., Davis, J. J., Disz, T. & Edwards, R. A. RASTtk: A modular and extensible implementation of the RAST algorithm for building custom annotation pipelines and annotating batches of genomes. Sci. Rep. 5, (2015).

65. Wattam, A. R. et al. Improvements to PATRIC, the all-bacterial Bioinformatics Database and Analysis Resource Center. Nucleic Acids Res. 45:D535–D542, (2017).

66. Kanehisa, M., Sato, Y. & Morishima, K. BlastKOALA and GhostKOALA: KEGG tools for functional characterization of genome and metagenome sequences. J. Mol. Biol. 428, 726–731 (2016).

67. Bland, C. et al. CRISPR recognition tool (CRT): a tool for automatic detection of clustered regularly interspaced palindromic repeats. BMC Bioinform. 8, (2007).

68. Goris, J. et al. DNA-DNA hybridization values and their relationship to whole-genome sequence similarities. Int. J. Syst. Evol. Microbiol. 57, 81–91 (2007).

69. Richter, M. & Rosselló-Móra, R. Shifting the genomic gold standard for the prokaryotic species definition. Proc. Natl. Acad. Sci. USA. 106, 19126–19131 (2009).

70. Zhang, H. et al. a meta server for automated carbohydrate-active enzyme annotation. Nucleic Acids Res. 46:W95–W101, (2018).

71. Huerta-Cepas, J. et al. Fast Genome-Wide Functional Annotation through Orthology Assignment by egg-NOG-Mapper. Mol. Biol. Evol. 34, 2115–22 (2017).

72. El-Gebali, S. et al. The Pfam protein families database in 2019. Nucleic Acids Res. 47:D427–D432, (2019).

73. Rawlings, N. D. et al. The MEROPS database of proteolytic enzymes, their substrates and inhibitors in 2017 and a comparison with peptidases in the PANTHER database. Nucleic Acids Res. 46:D624–D632, (2018).

74. Yu, N. Y. et al. PSORTb 3.0: improved protein subcellular localization prediction with refined localization subcategories and predictive capabilities for all prokaryotes. Bioinformatics 26, 1608–1615 (2010).

75. Shivani, Y., Subhash, Y., Ch, S. & ChV, R. Description of “Candidatus Marispirochaeta associata” and reclassification of Spirochaeta bajacaliforniensis, Spirochaeta smaragdinae and Spirochaeta sinaica to a new genus Sediminispirochaeta gen. in *nov. as Sediminispirochaeta bajacaliforniensis comb. nov., Sediminispirochaeta smaragdinae comb. nov. and Sediminispirochaeta sinaica com*b. nov. Int. J. Syst. Evol. Microbiol. 66, 5485–5492 (2016).

76. Shivani, Y., Subhash, Y., Ch, S. & ChV, R. Spirochaeta lutea sp. nov., isolated from marine habitats and emended description of the genus Spirochaeta. Syst. Appl. Microbiol. 38, 110–114 (2015).

77. Hansen, T. A. & Veldkamp, H. *Rhodopseudomonas sulfidophila*, nov. spec., a new species of the purple nonsulfur bacteria. Arc.h Microbiol. 92, 45–58 (1973).

78. Takai, K. et al. Isolation and physiological characterization of two novel, piezophilic, thermophilic chemolithoautotrophs from a deep-sea hydrothermal vent chimney. Env. Microbiol. 11, 1983–1997 (2009).

79. Miroshnichenko, M. L., et al. *Oceanithermus profundus* gen. nov., sp. nov., a thermophilic, microaerophilic, facultatively chemolithoheterotrophic bacterium from a deep-sea hydrothermal vent. Int. J. Syst. Evol. Microbiol. 53, 747–752 (2003).

80. Subhash, Y., Ch, S. & ChV, R. Flavobacterium aquaticum sp. nov., isolated from a water sample of a rice field. Int. J. Syst. Evol. Microbiol. 63, 3463–3469 (2013).

81. Bale, N. J. et al. Diagnostic amide products of amino lipids detected in the microaerophilic bacteria Lutibacter during routine fatty acid analysis using gas chromatography. Org. Geochem. 144, (2020).

82. Damsté, J. S. S. et al. Dominance of mixed ether/ester, intact polar membrane lipids in five species of the order Rubrobacterales: Another group of bacteria not obeying the “lipid divide”. Syst. Appl. Microbiol. 46, 126404 (2023).

83. DM, V., et al. Pontiella desulfatans gen. Nov Sp Nov Pontiella Sulfatireligans Sp Nov Two Mar. Anaerobes Pontiellaceae Fam. 8, (2020).

84. Ravot, G., et al. *Fusibacter paucivorans* gen. nov., sp. nov., an anaerobic, thiosulfate-reducing bacterium from an oil-producing well. Int. J. Syst. Bacteriol. 49, 1141–1147 (1999).

85. Leschine, S. & Canale-Parola, E. Rifampin-resistant RNA polymerase in spirochetes. FEMS Microbiol. Lett. 35, 199–204 (1986).

86. Podosokorskaya, O. A., Bonch-Osmolovskaya, E. A., Novikov, A. A., Kolganova, T. V. & Kublanov, I. Ornatilinea apprima gen. nov., sp. nov., a cellulolytic representative of the class Anaerolineae. Int. J. Syst. Evol. Microbiol. 63, 86–92 (2013).

87. Na, S. I. et al. Up-to-date bacterial core gene set and pipeline for phylogenomic tree reconstruction. J. Microbiol. 56, 280–285 (2018).

