## Supplementary Information for "Organic matter degradation in the deep, sulfidic waters of the Black Sea: Insights into the ecophysiology of novel anaerobic bacteria"

[2]Faculty of Geosciences. Department of Earth Sciences Utrecht University., P.O. Box 80.021, 3508 TA Utrecht, The Netherlands.

[3]Department of Microbiology, Radboud Institute for Biological and Environmental Sciences, Radboud University, Nijmegen, The Netherlands.

**Running title:** Cultivation and physiology of novel piezotolerant bacteria from the Black Sea.

***Author for correspondence:** Laura Villanueva

**Keywords:** Black Sea; piezotolerant; sulfidic waters; organic matters; *Clostridiales*; *Marinifilaceae*; *Planctomycetota*, *Cloacimonadota*; *Ignavibacteriota*; *Desulfobacterota*.

**Microbial community analysis of the enrichments**

Firstly, we performed 16S rRNA gene amplicon sequencing to determine the diversity of the collected sulfidic water from 2,000 m depth. This contained 16S rRNA gene sequences affiliated to the phyla *Fusobacteriota*, *Cloacimonadota*, *Planctomycetota*, *Chloroflexota*, *Desulfobacterota*, *Bacteroidota*, *Marinimicrobia*, and *Omnitrophica* (Fig. 1A-i), in good agreement with the microbial diversity previously reported[1–4]. Next, we used different treatments to enrich specific microbes. The growth medium amended with simpler carbon sources like acetate (Fig. 1A-ii) and pyruvate (Fig. 1A-iii) mostly supported enrichment of members of the phyla *Bacteroidota* (55.4 and 84.4%, respectively) and *Desulfobacterota* (41.0 and 11.7%, respectively), while amino acid mixtures supported *Bacteroidota* (79.0%) and alphaproteobacterial members (18.9%; Fig. 1A-iv). A significant increase in the relative abundance of *Cloacimonadota* (~17%) and *Planctomycetota* (~30-42%) was observed in the growth media amended with propionate and chitin, respectively (Fig. 2).

The growth medium containing cellulose as the major carbon source promoted a much greater range of microbial members affiliated with *Bacteroidota*, *Clostridiales*, *Alphaproteobacteria*, *Desulfobacterota*, *Epsilonproteobacteria*, *Spirochaetota*, *Ignavibacteriota*, and *Cloacimonadota* (Fig. 1A-vii-x) (BS1; see M & M for details). Growth of such a wide variety of microbial members was expected, given that the gradual hydrolysis of cellulose gently generates a myriad of carbon sources that function as substrates for diverse physiological groups of microorganisms. Moreover, the gentle release of carbon sources in the medium is advantageous since most of the uncultivated microorganisms are not ready to cope with the sudden exposure of carbon sources[5–7]. Application of a diluted cellulose medium (BS2) incubated at a lower temperature (10°C) and higher sulfide concentration (>2 mM) promoted the growth of an even broader range of bacteria, including *Ignavibacteriota*, *Cloacimonadota*, *Chloroflexota*, and *Planctomycetota* (Fig. 1A-ix). Further relative increases in the abundance of members of *Ignavibacteriota*, *Cloacimonadota*, and *Planctomycetota* (Fig. 1A-x) were observed in the cellulose enrichments incubated at 10°C at even higher sulfide concentrations (4 mM; BS3 medium). An increase in the abundance of members of these phyla might be possible due to their adaptation to *in-situ* conditions.

**Physiology and metabolic properties of the *Psychrilyobacter piezotolerans* strain S5**

*Psychrilyobacter* sp. strain S5 was isolated by repeatedly streaking on cellulose amended medium (see M&M for details; Fig. 1B-i), and it was found to have a 100% 16S rRNA gene sequence similarity with the *Psychrilyobacter piezotolerans* strains SD5$^T$ and BL5 of the phylum *Fusobacteriota* previously isolated from the Black Sea[1] (Fig. S1A). The strain S5 was initially rod-shaped and Gram-negative in growth medium containing 1 mM sulfide (Fig. S1C), but at higher sulfide concentrations (>20 mM) its cell morphology transformed into a spherical shape, as previously described[1]. While strain S5 shared similar physiology and metabolic properties with *P. piezotolerans* strains SD5$^T$ and BL5[1], it differs from them in its inability to use chitin, differences in optimal growth temperature range (18-23 °C) and lipid profile (Table S42-S43; by relatively lower abundance of hydroxy fatty acids). Based on phenotypic and genotypic analyses this culture could represent as an additional strain of the *Psychrilyobacter piezotolerans*.

**Genome characteristics, physiology, and metabolic properties of *Clostridiales* bacteria strains A1$^T$ and A2**

The genomes of *Clostridiales* bacteria strains A1$^T$ and A2 were 4,111,109 bp and 4,116,947 bp in length, respectively. These genomes were found to be almost complete and free of contamination (Table S6). The G+C mol% of the A1$^T$ and A2 strains were 42.88% and 42.87%, respectively. Annotation of the genomes revealed that strain A1$^T$ had 3,889 coding sequences, while A2 had 3,901. No CRISPR repeats were identified in either genome. Based on a BLAST search analysis of the complete 16S rRNA gene sequences (1,541 bp), strains A1$^T$ and A2 showed 90.3% similarity with *Fusibacter paucivorans* SEBR 4211$^T$ of the order *Clostridiales* of the phylum *Bacillota*. The genomes of both strains contained a complete set of genes encoding enzymes of the Embden-Meyerhof glycolytic pathway (Fig. S14). The pentose phosphate pathway was represented only by a non-oxidative branch, as the transaldolase gene was not found. The tricarboxylic acid cycle was incomplete due to the lack of malate dehydrogenase and was likely only used for biosynthetic purposes. Interestingly, an upper TCA cycle metabolite 2-oxoglutarate was a key intermediate in the biosynthesis of some amino acids. Both strains possessed all the genes involved in the 2-C-methyl-D-erythritol4-phosphate/1-deoxy-D-xylulose 5-phosphate (MEP/DOXP) pathway of isoprenoid biosynthesis (Fig. S14). Hydrogen was a major fermentation product in both strains and may be produced by [FeFe]-hydrogenase, which is known to couple the oxidation of reduced ferredoxin to the evolution of $H_2$ during carbohydrate and protein fermentation[8]. The reversible electron transfer between NADP(H) and ferredoxin could be performed by ferredoxin-NADP (+) reductase. The presence of V-type ATPase indicates that strains A1$^T$ and A2 may rely on substrate-level phosphorylation for ATP production.

Both strains (A1$^T$ and A2) were strict anaerobic bacteria and did not show catalase and oxidase activity, which is consistent with their anaerobic lifestyle. However, they were able to tolerate air exposure for about an hour, indicating some level of aerotolerance. This property was supported by the presence of genes coding for superoxide reductase, superoxide dismutase, rubredoxin, and rubrerythrin that might help in oxygen detoxification. Our observation suggests that aerotolerance in these strains could be an adaptation to neutralize oxygen exposure in the upper water column of the Black Sea. The strains could use thiosulfate and sulfur as terminal electron acceptors, but not sulfate. They were unable to oxidize sulfide but were able to grow at sulfide concentrations up to 7 mM at pH 7.0, which is 17.5-fold higher than the natural concentration. Under strict anaerobic conditions, both strains were able to grow at hydrostatic pressures up to 50 MPa. However, growth at higher pressures (>20 MPa) was slower and the doubling time was extended to 8-10 hours at 30 MPa. This observation suggests that these strains are adapted to the extreme conditions of the deep sulfidic waters of the Black Sea.

**Genome characteristics, physiology, and metabolic properties of novel species of the phylum *Spirochaetota* (strains M1$^T$, M2 and S2)**

*Oceanispirochaeta* sp. strains M1$^T$ and M2 have similar genome sizes of 5.88 and 5.87 Mbp, respectively, with genome completeness of 99.73% and contamination levels of 3.73% and 4.13%, respectively (Table S6). Both strains have a G+C content of 42.89% and 56 tRNA and 98 CRISPR repeats. Strain M1$^T$ has 5440 CDS with 2333 functional proteins, while strain M2 has 5495 CDS with 2,120 functional proteins. *Sphaerochaeta* sp. strain S2 has a genome size of 3.1 Mbp with a genome completeness of 98% and 1.2% contamination (Table S6). It has a G+C content of 46.87%, 43 tRNA, 6 rRNA, and 14 CRISPR repeats, with 3368 genes, of which 1,589 have functional assignments.

All three strains preferentially fermented glucose for their growth (Table S1, S3), which was further supported by the presence of relevant genes in their genomes (Fig. S18). While the strains possessed enzymes for the hydrolysis of polysaccharides like starch (glucoamylase), they could not utilize starch or cellobiose for their growth (Table S1). *Sphaerochaeta* sp. strain S2 stopped growing during repeated subculturing under laboratory conditions; therefore, we present the relevant findings of organic matter degradation for this strain under the Candidatus option.

The genomes of all three strains contained a complete set of genes encoding enzymes of the Embden-Meyerhof glycolytic pathway (Fig. S18). However, the pentose phosphate pathway is incomplete, lacking the transaldolase gene and only having a non-oxidative branch. The tricarboxylic acid cycle is also incomplete due to the absence of the malate dehydrogenase gene, and it is likely used only for biosynthetic purposes. The preferred carbon source for all three strains was pyruvate, as evidenced by their ability to utilize it for growth and the presence of pyruvate-formate lyase and pyruvate-ferredoxin oxidoreductase genes in their genomes (Fig. S18). Lactate dehydrogenase was also present, indicating that lactate could be produced by fermentation, along with acetate. The cytoplasmic [FeFe] group A hydrogenase can couple the oxidation of reduced ferredoxin generated by pyruvate-ferredoxin oxidoreductase to the reduction of protons to hydrogen during fermentative growth. V-type ATPase was found, indicating that these strains rely on substrate-level phosphorylation for ATP production. They can hydrolyze protein (casein) and ferment aspartate and glutamate for growth, as supported by the presence of peptidase and amino acid fermentation genes in their genomes (Fig. S18). However, aerobic, or anaerobic respiration is not possible due to the absence of related genes in the genomes. Hydrogen is a major fermentation product in all three strains, likely produced by [FeFe]-hydrogenase, which is known to couple the oxidation of reduced ferredoxin to the evolution of hydrogen during carbohydrate and protein fermentation[8].

All three strains were found to be Gram-negative bacteria (Fig. S3) and exhibited negative reactions for oxidase and catalase activities. They were able to tolerate oxygen exposure for up to 30 minutes, and the presence of superoxide reductase genes in strains $M1^T$ and M2 supported this observation. Strain S2, on the other hand, contained catalase and superoxide dismutase genes in addition to the aforementioned genes (Fig. S18). All three strains also contained genes encoding rubredoxin and rubrerythrin, which could act as oxygen scavengers. None of the strains could utilize sulfate, sulfite, thiosulfate, nitrate, or elemental sulfur as electron acceptors, as evidenced by the absence of the corresponding genes in their genomes. Sulfide oxidation was not possible in any of the strains. However, they were able to grow at sulfide concentrations of up to 7 mM at pH 7.0, which was 17.5-fold the natural concentration. Under strict anaerobic conditions, all three strains were able to grow at hydrostatic pressures of up to 50 MPa. However, they exhibited weak growth at elevated hydrostatic pressures (>20 MPa), and their doubling time was extended to 8-10 hours at 30 MPa (strain M1$^{T}$; Fig. 4C).

**Genome characteristics, physiology, and metabolic properties of novel members of the phylum *Bacteroidota* (strains S6, L6, B1$^{T}$, B2, M2P and SYP)**

We have isolated six bacterial strains (S6, L6, B1$^{T}$, B2, M2P and SYP) affiliated with phylum *Bacteroidota* from cellulose enrichments. The genome size of *Lentimicrobium* sp. strain S6 was 5,722,269 bp with a genome completeness of 98.92%, 2.24% contamination, and a G+C content of 35.04%. In comparison, *Lentimicrobium* sp. strain L6 had a smaller genome size of 5,144,386 bp with a genome completeness of 96.24%, 1.70% contamination, and a G+C content of 35.03%. Based on a BLAST search analysis of the complete 16S rRNA gene sequences, strains S6 and L6 showed 88.7% similarity with *Lentimicrobium saccharophilum* TBC1$^{T}$, which belongs to the family *Lentimicrobiaceae* of the phylum *Bacteroidota*. Due to the extremely slow growth of strains S6 and L6, we could delay the deposition of these strains to recognized culture collections. However, due to the high relevance of the findings reported here for OM degradation, we chose to use the *Candidatus* option for strain S6.

*Lutibacter* sp. strain B1$^{T}$ had a genome size of 3,428,913 bp with a genome completeness of 97.65%, 1.10% contamination, and a lower G+C content of 30.27%. It had 3110 genes and 1929 proteins with functional assignments, along with 1,251 hypothetical proteins, 39 tRNA, and 4 rRNA. Interestingly, it also had 31 CRISPR repeats. *Lutibacter* sp. strain B2 had a slightly smaller genome size of 3,240,965 bp with a genome completeness of 97.16%, 0.35% contamination, and a G+C content of 30.83%. It had 3,155 CDS and 1,759 proteins with functional assignments, along with 1396 hypothetical proteins and 86 tRNA. One copy of 16S rRNA gene sequences was identified in the genome of strains B1$^T$ and B2. 16S rRNA gene sequence obtained from the draft genome sequence was identical to the one retrieved with the 16S rRNA gene PCR on the DNA extracted from the strain B1$^T$ and B2. 16S rRNA gene sequence similarity between strains B1$^T$ and B2 was 100%. Strains B1$^T$ and B2 showed 96.25% 16S rRNA gene sequence similarity with *Lutibacter profundi* LP1$^T$ and <95.8% with other members of the genus *Lutibacter*.

*Ancylomarina euxinus* strain M3P had a genome size of 4,351,206 bp with a genome completeness of 100%, no contamination, and a higher G+C content of 35.8%. It have 3,643 genes and 1,855 proteins with functional assignments, along with 1788 hypothetical proteins, 85 tRNA, and 8 rRNA. Like *Lutibacter* sp. strain B2, it have 22 CRISPR repeats.

The genomes of all six strains contain the genes involved in both the 2-C-methyl-D-erythritol 4-phosphate/1-deoxy-D-xylulose 5-phosphate (MEP/DOXP) and mevalonate pathways of isoprenoid biosynthesis (Fig S25). Growth of all six strains was observed in medium amended with either or both fosmidomycin (a pathway inhibitor of the MEP pathway) and simvastatin (a pathway inhibitor of the mevalonate pathway), indicating that both pathways are functional for isoprenoid biosynthesis. Under a microscope, flagellar motility was observed, and all six strains grew chemoheterotrophically. Fermentation of various mono- and disaccharides (such as glucose, galactose, xylose, and maltose) as well as pyruvate, lactate, and glycerol were observed (Table S8-S9, S12). Genes for central metabolic pathways, including glycolysis, the TCA cycle, and the reductive TCA cycle, were found in the genomes, supporting the capability of the strains to ferment glucose. However, autotrophic growth was not observed in physiological tests using $H_2/CO_2$ as an electron donor and carbon source. Propionate was likely formed via the methylmalonyl-CoA pathway. The ability to use pyruvate as a substrate was confirmed by the presence of a pyruvate-formate lyase, which yields acetate and formate, and a pyruvate-ferredoxin oxidoreductase, which yields acetate, $CO_2$, and $H_2$ (Fig. S25). Furthermore, fermentation of lactate and maltose was confirmed by the presence of genes encoding their respective degradation pathways.

The physiological behavior toward oxygen differed among the strains. Strains B1$^T$ and B2 preferentially grew under microaerophilic conditions, while strains S6, L6, M2P, and SYP grew well under strict anaerobic conditions. Moreover, the genomes of strains B1$^T$ and B2 encoded a cbb3-type cytochrome c oxidase, which is comprised of three subunits and cytochrome c oxidase. These enzymes function as terminal oxygen reductases and have been suggested to be associated with microaerophilic growth[9,10]. Genes for oxygen detoxification (i.e., superoxide dismutase, catalase, and cytochrome c551 peroxidase) were identified in the genomes of all six strains tested (Fig S25). Reduction of sulfate to sulfide, nitrate to nitrite or nitrite was not observed in the presence of glucose in all the tested strains. All the isolated strains utilized glucose, pyruvate, acetate, threonine, lysine, glutamate, and aspartate (Table S4-S10, S12) for growth. However, no growth was observed on cellulose, cellobiose, and chitin. No strains used sulfate, sulfite, thiosulfate, nitrate, fumarate, and elemental sulfur as an electron acceptor. They were also able to grow at sulfide concentrations up to 9 mM at pH 7.0, which was 22.5-fold the natural concentration. All strains were able to grow up to 50 MPa of hydrostatic pressure (Fig. 5D-5E), but they grew weakly when the hydrostatic pressure was increased beyond 20 MPa.

**Genome characteristics, physiology, and metabolic properties of novel members of the phylum *Desulfobacterota* (strains S3$^T$ and S3-i)**

We have isolated two bacterial strains (*Pseudodesulfovibrio* sp. strains S3$^T$, and S3-i;) affiliated with phylum *Desulfobacterota*. Whole genome sequencing of strains S3$^T$ and S3-i yielded genomes of 3,706,001 bp and 3,705,660 bp in length, respectively after assembly (Table S6). The G+C mol% of strains S3$^T$ and S3-i was 56.6 and 56.60%, respectively. Annotation of the genome of strain S3$^T$ and S3-i indicated 3,612 and 3,607 coding sequences, respectively. One copy of 16S rRNA gene sequences was identified in the genome of strains S3$^T$ and S3-i. 16S rRNA gene sequence obtained from the draft genome sequence was identical to the one retrieved with the 16S rRNA gene PCR on the DNA extracted from the strain S3$^T$ and S3-i. Based on BLAST search analysis of the 16S rRNA gene sequences, strains S3$^T$ and S3-i showed 97.4% similarity with *Pseudodesulfovibrio indicus* J2$^T$ of the family *Desulfovibrionaceae* of the phylum *Desulfobacterota*.

Strains S3$^T$ and S3-i were sulfate-reducing bacteria grew on a limited number of organic substrates, with lactate, pyruvate, and acetate (Table S5). In support of such activities, secreted glycoside hydrolases were not detected in the genomes of these strains. The glycolytic pathway probably operates in the direction of gluconeogenesis, as indicated by the presence of genes coding phosphoenolpyruvate synthase and fructose-1,6-bisphosphatase which specifically perform the reverse reactions (Fig. S24). Pyruvate could be reversibly decarboxylated to acetyl-CoA by pyruvate: ferredoxin oxidoreductase. Furthermore, conversion of acetyl-CoA to acetate with the production of ATP can be performed by acetyl-CoA synthetase. Oxidation of lactate to pyruvate was probably facilitated by putative lactate dehydrogenases (LDH). The tricarboxylic acid (TCA) cycle in strains S3$^T$ and S3-i are incomplete, lacking citrate synthase and succinyl-CoA synthetase. This finding is consistent with the observed inability of strains S3$^T$ and S3-i to oxidize organic substrates completely. For example, growth of strain S3$^T$ and S3-i with formate was only possible in the presence of acetate as an auxiliary carbon source, consistent with the absence of known pathways for autotrophic C1 fixation. In particular, the Wood-Ljungdahl (reductive acetyl-CoA) pathway, frequently used by autotrophic sulfate reducers, is incomplete, lacking carbon monoxide dehydrogenase/acetyl CoA synthase.

The genomes of $S3^T$ and S3-i contain all genes necessary for dissimilatory sulfate reduction (Fig. S24). Another linkage of sulfate-reduction enzymes to the membrane is enabled by the sulfite reductase-associated electron transfer complex DsrMKJOP with subunits DsrM and DsrP containing transmembrane domains. Four hydrogenases of the [NiFe]-family and one formate dehydrogenase are encoded by the strains $S3^T$ and S3-i genomes. Consistently, like other members of the family *Desulfovibrionaceae,* $S3^T$ and S3-i grew with $H_2$ as an energy source in the presence of acetate as a carbon source. The first hydrogenase is encoded by eight-gene operon. The [NiFe] uptake hydrogenase, could oxidize $H_2$, by donating the electrons to the quinone pool via the third cytochrome *b* subunits linking them to the cytoplasmic membrane. The electron transfer from this soluble periplasmic complex to the cytoplasmic membrane may be facilitated by a pool of *c*-type cytochromes present in the periplasm[11,12]. This electron transport pathway probably ends at the membrane linked Hmc complex[13]. Cytochrome c, HmcA, may accept electrons from periplasmic cytochromes. Genes coding for cytoplasmic hydrogen: heterodisulfide oxidoreductase, consisting of CoB-CoM heterodisulfide reductase (HdrACB) and methyl viologen-reducing hydrogenase (MvhDGA) were also detected which catalyzes the endergonic reduction of ferredoxin and the exergonic reduction of heterodisulfide, coupled to $H_2$ oxidation by electron bifurcation involving HdrA[14].

Genomes of strains $S3^T$ and S3-i also contained four subunits of soluble cytoplasmic hydrogenase. These hydrogenases are bidirectional and can re-oxidize the cofactors by using protons as electron acceptors[15]. The presence of a NADPH-binding motif suggests that this hydrogenase can use NADPH in hydrogen turnover reactions. The presence of formate dehydrogenase explains the observed ability of strains S3$^T$ and S3-i to use formate as an electron donor. The presence of a N-terminal Tat signal peptide in FdhA suggests that it is in the periplasm. Like the periplasmic uptake hydrogenase, formate dehydrogenase lacks a membrane subunit, and electron transfer to the membrane is probably performed via the periplasmic cytochromes and Hmc complex. Genomes of strains S3$^T$ and S3-i also contain several other membrane-linked oxidoreductases that can contribute to the generation of transmembrane ion gradient and/or the use of alternative electron acceptors to sulfate. Two putative complexes similar to the bacterial NADH:quinone oxidoreductase are present. Both comprise the subunits like NuoA, B, C, D, H, I, J, K, L, M, and N, while the genes for the subunits NuoEFG that form the NADH dehydrogenase module are missing, indicating that NADH is likely not an electron donor. The first cluster is linked to genes coding for two subunits of CISM oxidoreductases of the Psr/Psh family[16]: the molybdopterin-binding catalytic subunit A and the iron-sulfur electron transfer subunit B. It is possible that such an arrangement indicates coupling of transmembrane proton transfer, performed by the core subunits of NADH: quinone oxidoreductase, with the oxidation or reduction of sulfur compounds. Oxidation of pyruvate produces reduced ferredoxin that could provide electrons to this oxidoreductase. Genes encoding molybdopterin family oxidoreductase consisting of all three subunits: A, B and membrane subunit C of the NrfD family. The catalytic A subunit was predicted to contain a N-terminal Tat signal peptide and is phylogenetically related to thiosulfate or polysulfide reductases. The presence of putative thiosulfate reductase, capable of producing sulfide and sulfite from thiosulfate, explains the ability of strains S3$^T$ and S3-i to use thiosulfate as an electron acceptor.

Despite the observed inability of strains $S3^T$ and S3-i to grow by nitrite reduction, its genome contains cytochrome *c* nitrite reductase[17], comprising large NrfA and small NrfH subunits with five and four hemes, respectively. The presence of an N-terminal signal peptide in the large subunit suggests that this complex faces the periplasmic side of the membrane. The physiological role of nitrite reductase could be detoxification of nitrite, which is known as an inhibitor of sulfate-reducing organisms[18]. A similar function in detoxification of oxygen could be assigned to the cytochrome *bd* ubiquinol oxidase. Furthermore, all three strains were able to grow at sulfide concentrations up to 9 mM at pH 7.0, which was 22.5 -fold of the *in-situ* environmental conditions. All strains were able to grow up to 50 MPa of hydrostatic pressure. A significant improvement in the growth was observed in the glutamate amended medium at elevated hydrostatic pressure (30 MPa; Fig. 5F).

**Genome characteristics and metabolic properties of *Mycoplasmatota* bacterium strain Izemo-BS**

The MAG of '*Ca* Atrimarinoplasma cellobiosiphila' strain Izemo-BS was 2,431,619 base pairs in size and has a genome completeness of 97.33%. The MAG has a low contamination rate of 1.3% and the G+C content was 31.1%. The genome contains 2,233 protein-coding genes, and out of these, 930 have functional assignments, while 1,303 are hypothetical proteins. The MAG contains 33 tRNA genes that are responsible for translating genetic information into proteins. Various genes e.g., endoglucanase, alpha-amylase (EC 3.2.1.1), amylomaltase, beta-glycosyl hydrolase, glycosyltransferase was detected in Izemo-BS which are involved in the hydrolysis of polysaccharides (cellulose and starch) supporting the capabilities to obtain energy by hydrolyzing the polysaccharides. Furthermore, the genes responsible for the degradation of cellobiose (cellobiose phosphorylase) was also identified. Various genes involve in the glucose fermentation (Fig. S16) are also present indication fermenting lifestyles in the sulfidic waters. Various genes encoding endonuclease, exonuclease, and extracellular ribonuclease are also detected (Fig. S16) in the MAG which further which is in line with the earlier report[19,20]. Genes involved in the glycolysis were also present, however, complete absence of electron transfer chain involved in the anaerobic respiration indicates that Izemo-BS obtain their energy through the degradation of DNA and sugars. Largely, the various genes detected in genome indicated that Izemo-BS might be actively involved in the degradation of DNA and simpler carbon sources like cellobiose; a component of the organic matter sinking from upper oxic zones of Black Sea.

#### Genome characteristics and metabolic properties of *Cloacimonadota* bacterium strain Cloa-SY6

The MAG of the *Cloacimonadota* bacterium strain Cloa-SY6 was 3,336,172 bp in size. The genome is estimated to be 98.84% complete with low contamination (1.1%). The G+C content of the genome is 34.31%. The genome contains 2,981 protein-coding genes, of which 1,111 have functional annotations and 1,870 are hypothetical proteins. There are 184 clustered regularly interspaced short palindromic repeats (CRISPRs) in the genome, which suggests that the bacterium may have the ability to defend against viral attacks. The genome also encodes 55 transfer RNA genes a complete set of ribosomal RNA gene operon. The 16S rRNA gene sequence (1535 bp) is identical to that obtained from 16S rRNA gene amplicon analysis. The Cloa-SY6 strain has 82.72%, 83.8% and 81.9% 16S rRNA gene similarities with '*Ca* Cloacamonas acidaminovorans' strain Evry, '*Ca* Syntrophosphaera thermopropionivorans', and bin40 (16S rRNA gene sequence recovered from the bin40; *Cloacimonadota* MAG obtained from the Black Sea) respectively (Fig. S8A). Comparative analysis of Cloa-SY6 along with '*Ca* Cloacamonas acidaminovorans', '*Ca* Syntrophosphaera thermopropionivorans', and other MAGs obtained from the Black Sea (bin40, bin80, bin85 and bin108) showed distinct differences (Table S8). The genome sequence of Cloa-SY6 is relatively larger than that of '*Ca* Cloacamonas acidaminovorans', '*Ca* Cloacamonas acidaminovorans', *'Ca* Syntrophosphaera thermopropionivorans', and other MAGs obtained from the Black Sea (bin40, bin80, bin85 and bin108)[21] indicating their independent nature with respect to various biosynthetic pathways (Fig. S21).

Members of the phylum *Cloacimonadota* have been frequently observed in both engineered and natural habitats and are mostly known for their involvement in sugar transformation[21–30]. However, the MAG of Cloa-SY6 contained a limited number of 64 CAZymes, most of which are GTs (n=45) involved in the initiation and elongation of glycan chains[31]. Only 11 CAZymes were classified as GHs, indicating limited carbohydrate utilization capability. While the increased abundance in the propionate medium suggests a preference for propionate, like *'Ca* Syntrophosphaera thermopropionivorans'. Moreover, genome of Cloa-SY6 encoded most of the genes involved in the amino acid biosynthesis (Fig. S21) which further reflects their independent nature. In contrast, *Cloacimonadota* MAGs (bin40, bin80, bin85 and bin108) obtained earlier[21] from the Black Sea lacks several genes involved in the amino acid biosynthesis.

Cloa-SY6 encodes both a rudimentary respiratory and fermentative pathway for energy generation. In the absence of a canonical electron transport chain (ETC) for generating a membrane potential, we assume that Rnf electron transport complexes are likely sources of a transmembrane ion gradient[32] in Cloa-SY6. Various enzymes related to anaerobic lifestyles were detected in the MAG of Cloa-SY6, including ribonucleoside triphosphate reductase, ferredoxin oxidoreductases, and radical S-adenosylmethionine-dependent proteins, indicating that Cloa-SY6 is well adapted to the permanently anoxic conditions of the Black Sea. However, the presence of genes related to microaerophilic growth (such as superoxide reductase; EC 1.15.1.2), ruberythrin, and thioredoxin reductase (EC 1.8.1.9) in Cloa-SY6 suggests an adaptation to survive in the suboxic zones of the Black Sea.

Cloa-SY6 differed from previously reported MAGs due to its relatively larger genome size, presence of various genes involved in amino acid biosynthesis, and lack of many CAZymes. Additionally, it exhibits low 16S rRNA gene sequence similarity and have distinct phylogenomic differences (Fig. S8A-B), which suggests that Cloa-SY6 belongs to a novel taxon at the order level within the phylum *Cloacimonadota*.

#### Genome characteristics and metabolic properties of *Planctomycetota* bacterium strain Plnct-SY6

We recovered a high-quality metagenome-assembled genome (MAG) affiliated with this phylum, as shown in Table S3. The MAG was 5,418,934 base pairs in size and had a genome completeness of 97.66%, indicating that most of the expected genes were present in the genome. The genome had a low contamination rate of 1.14%. The G+C content of the genome was relatively high at 41.35%. The MAG contained 4,562 protein-coding genes, out of which 1,332 had functional assignments, while 3,230 were hypothetical proteins whose function was not yet known. The genome also contained 47 tRNA genes that were responsible for translating genetic information into proteins. A complete 23S rRNA gene was present in the genome sequence, showing 83.8% similarity with *Phycisphaerae* bacterium ST-NAGAB-D1 of the order *Sedimentisphaerales* in the phylum *Planctomycetota*. Phylogenetic analysis confirmed their affiliation with an uncultivated clade of this phylum thriving in deep marine habitats (Fig. S7A). Additionally, the genome of the '*Ca* Atrisphaera chitinolytica' strain Plnct-SY6 had 79 CRISPR repeats, which are commonly found in bacterial genomes and play a crucial role in the immune system of bacteria against phage infection.

The genes encoding metabolic pathways common for chemoorganotrophic bacteria, such as glycolysis, the citrate cycle, the pentose-phosphate pathway, and oxidative phosphorylation are present. The Plnct-SY6 has the genomic potential for synthesis of all amino acids. Two fructosetype sugar-specific subunits of the phosphotransferase system could be found in Plnct-SY6. The survey for genes related to cell division revealed that the FtsZ-encoding gene was absent, while two copies of the gene coding for DNA translocase FtsK were present in the genome of Plnct-SY6. The gene encoding the key enzyme for synthesis of N-methylated phosphorus-free ornithine membrane lipids, N-methyltransferase (OlsG), is identified in the genome of Plnct-SY6 which might help their growth under phosphate limiting conditions. Genomic analysis revealed the presence of the major components of electron transfer chain i.e., proton-translocating NADH-dehydrogenase complexes, a membrane-bound succinate dehydrogenase/ fumarate reductase, isoprenoid quinones, and a $F_0F_1$-type bacterial ATP synthase.

**Genome characteristics and metabolic properties of *Ignavibacteriota* bacterium strain Igna-SY6**

The size of '*Ca* Ponteuxinibacter sulfidophilus' strain Igna-SY6 was 4,731,990 base pairs, with a genome completeness of 100%. The G+C content of the genome was 32.95%. The genome contained 3,922 protein-coding genes, out of which 1,809 had functional assignments, while 2,113 were hypothetical proteins whose function was not yet known. The genome also contained 41 tRNA genes responsible for translating genetic information into proteins. Additionally, the MAG of '*Ca* Ponteuxinibacter sulfidophilus' strain Igna-SY6 has 112 CRISPR repeats, which are commonly found in bacterial genomes and play a crucial role in the immune system of bacteria against phages and other invading elements.

The Igna-SY6 genome encodes a complete set of genes for glycolysis, the TCA cycle, and gluconeogenesis which indicates that it can grow with glucose or other oligosaccharides as sole carbon source (Fig. S22). The genome also includes genes for glycogen synthase and glycogen phosphorylase, which suggests that glycogen is its major storage compound. Genes for polyhydroxyalkanoate (PHA) synthesis and degradation were not detected, hence, it is assumed that Igna-SY6 probably produces acetate and L-lactate as the main products when growing fermentatively. Common pathways for fermentative production of propionate (via methylmalonyl-CoA carboxyltransferase), ethanol (via alcohol dehydrogenase), and formate (via pyruvate formate lyase) are present which indicate the fermentative growth mode. Igna-SY6 possesses genes encoding two $CO_2$ fixing enzymes pyruvate: ferredoxin oxidoreductase (PFOR) and 2-oxoglutarate: ferredoxin oxidoreductase (OFOR), which are essential for autotrophic $CO_2$ fixation in green sulfur bacteria[33]. Because the glyoxylate cycle is not present, PFOR is probably essential for the assimilation of acetate by carboxylation of acetyl-CoA to pyruvate. The Igna-SY6 genome encode citrate lyase, which is a key enzyme required for autotrophic $CO_2$ fixation by the reverse TCA cycle[34]. This enzyme catalyzes the cleavage of citrate to acetate and oxaloacetate, and it is involved in citrate fermentation in some organisms[35]. The operation of the reverse TCA cycle for $CO_2$ assimilation would also depend upon the availability of electron sources to produce reduced ferredoxin. The genome of Igna-SY6 includes genes necessary to take advantage of some potential electron sources and to produce reduced ferredoxin from them. Thus, the gene repertoire Igna-SY6 shows that the organism can probably grow mixotrophically. The Igna-SY6 genome encodes different ferredoxins and a variety of electron transfer complexes, including the RNF (Na+-translocating ferredoxin:NAD+ oxidoreductase) complex[36], two type-1 NADH dehydrogenase complexes, and alternative complex III (ACIII). The presence of such a broad array of electron transfer complexes likely reflects an ability of Igna-SY6 to utilize the different electron carriers used by various redox enzymes as well as the various terminal electron acceptors that might be available *in-situ* conditions of the Black Sea. Genome sequence of Igna-SY6 does not encodes the photosynthetic apparatus to produce the reduced ferredoxins required for carbon fixation by the reverse TCA cycle.

The Igna-SY6 has three different oxygen-dependent terminal oxidases, including cbb3-type heme-copper cytochrome c oxidases and two different cytochrome bd-quinol oxidases (Fig. S25). Both types of terminal oxidases could participate in aerobic respiration and/or protection against reactive oxygen species[37]. The cbb3 cytochrome oxidase and cytochrome bd-quinol oxidase typically have much higher affinity for $O_2$ than the caa3 cytochrome oxidase[38], and because of this, they are frequently involved in protecting anaerobes from reactive oxygen species. The presence of all three types of complexes in Igna-SY6 strongly suggests that Igna-SY6 experiences varying $O_2$ concentrations in-situ conditions of the Black Sea. The presence of these different terminal oxidases in Igna-SY6 would confer not only the ability to respire under oxic conditions but also the ability to protect oxygen-sensitive enzymes such as hydrogenase under microoxic conditions. 16S rRNA gene amplicon data analysis indicated that members of the phylum *Ignavibacteriota* are also present in the suboxic zone of the Black Sea. Genome of Igna-SY6 has genes encoding both catalase and superoxide dismutase, which protect organisms exposed to oxygen from reactive oxygen species. The genome of Igna-SY6 also encodes an oxygen-dependent protoporphyrinogen oxidase for heme biosynthesis, catabolic enzymes pyruvate dehydrogenase and 2-oxoglutarate dehydrogenase, which are typically found in aerobes. The presence of these genes in Igna-SY6 might be an adaptation to survive under oxic conditions. The genome of Igna-SY6 also encode two different [FeFe]-hydrogenases. Fe-only hydrogenases are often associated with $H_2$ evolution[15]. Thus, it appears likely that Igna-SY6 could use these enzymes to establish redox balance during fermentation The genome encodes a sulfide-quinone oxidoreductases which indicates that Igna-SY6 I can use sulfide as an electron donor which ultimately helps in the sulfide detoxification by formation of sulfur/polysulfide. The genome of Igna-SY6 is missing key genes involved in the biosynthetic pathways for several amino acids.

Hence, it expected that it might obtains them from its environment. Here, we detected them in the growth media containing yeast extract and tryptone which are rich in amino acids and oligopeptides which confirms their involvement in amino acid utilization.

**Genome characteristics and metabolic properties of *Chloroflexota* bacterium strain Chflx-SY6**

The *Chloroflexota* bacterium strain Chflx-SY6 has a genome size of 4,951,662 base pairs. The genome completeness is 99.09%, and the percentage of contamination is 7.27%. The G+C content of the genome is 43.12%. The genome contains 4,488 coding DNA sequences (CDS) and 1,705 proteins with functional assignments. Additionally, there are 2,783 hypothetical proteins, 40 tRNA, 1 rRNA, and 38 CRISPR repeats. The genome also has the mevalonate pathway of isoprenoid biosynthesis.

Potential for aerobic respiration via cytochrome C oxidase and the tricarboxylic acid (TCA) cycle were also identified in Chflx-SY6 (Fig. S23). However, we could not detect Chflx-SY6 in the enrichment cultures grown under aerobic or microaerophilic conditions which suggest that Chflx-SY6 could be a strict anaerobe. We identified a total of 146 genes coding for carbohydrate active enzymes in the genome sequence (Table S35). Genes coding for the hydrolysis of cellulose, xylan, and starch were present in the genome sequence, however, their low abundance in the cellulose medium indicated that such polysaccharides could not be preferred carbon sources for growth. All the genes involved in the Wood Ljungdahl (W-L) pathway were present in the genome sequence which could be involved in the carbon dioxide fixation.

Genomic analysis revealed the presence of the electron transfer chain i.e., proton-translocating NADH-dehydrogenase complexes, isoprenoid quinones, and a F0F1-type bacterial ATP synthase. Their presence indicates that Chflx-SY6 in the sulfidic waters have the potential to conserve energy via sugar fermentation/gluconeogenesis[39], the potentially reversible W-L pathway, pyruvate ferredoxin oxidoreductase, ATP synthase and NADH-quinone oxidoreductase (Fig. S23). Such energy metabolisms are also reported in the earlier studies from anoxic site indicate fermentation and acetogenesis as potential metabolisms in the subseafloor *Chloroflexaeota*[40,41]. Pyruvate ferredoxin oxidoreductase is also present which may provide a link between the W-L pathway and other anabolic pathways in Chflx-SY6 as suggested previously[19]. Genes coding ferredoxin and flavodoxin are likely involved as electron carriers in cellular redox reactions[42]. Peptide transporters and peptidases were also present in the genome sequence which indicate that Chflx-SY6 might be utilizing traces of proteinaceous components present in the sulfidic waters of the Black Sea.

**Membrane lipid characterization of the novel microbial taxa**

Membrane lipids are crucial components of various organisms, serving a vital role in the adaptation of marine microorganisms to their environment[1,43,44]. They can be unique to specific species, groups of organisms, or ecological processes[45,46]. We, therefore, analyzed the lipid profiles of pure cultures, excluding strains S2, S6, and L6, due to difficulties in generating sufficient biomass.

The fatty acid profile of *P. piezotolerans* strain S5 consisted $C_{16:0}$, $C_{16:1\omega7}$, $C_{14:0}$, $\beta$-OH $C_{12:0}$, and $\beta$-OH $C_{16:0}$ (Table S42). Major polar head groups identified were phosphatidylethanolamine (PE), phosphatidylglycerol (PG), cardiolipins, and lyso-PE. Cardiolipins were found to be crucial for *Psychrilyobacter* spp. as they contribute to the ability of bacteria to withstand high pressures[1]. This is consistent with earlier findings that reported the importance of cardiolipins in the physiology of bacteria living under extreme conditions[47].

$C_{14:0}$, $C_{16:1\omega9C}$, $C_{16:0}$, $C_{18:2}$, $C_{18:1\omega7c}$, and $C_{18:0}$ were major (>5%) core lipids of the *Clostridiales* bacterium strain A1$^{T}$ (Table S44). Strain A1$^{T}$ differed from its closest phylogenetic neighbor with respect to the absence of $C_{18:2}$ phylogenetic neighbor (*Fusibacter paucivorans* DSM 12116$^T$). Hence, it is likely that this core lipid could be an important signature for the *Clostridiales* bacterium strain A1$^T$ in the deep sulfidic waters of the Black Sea.

*Iso*-$C_{13:0}$, $C_{14:0}$, *iso*-$C_{15:0}$, and $C_{16:0}$ were present as major core lipids in *Oceanispirochaeta* sp. strains M1$^T$ and M2 along with a variety of minor core lipids (Tables S45-S46). Additionally, several diphosphatidylglycerols (DPGs, also known as cardiolipins) were detected with a mixed acyl/ether glycerol (AEG) core (Table S45-S46). The presence of alcohols and monoalkyl glycerol ethers in these strains might be related with their adaptation to the elevated hydrostatic pressure conditions of the deep sulfidic waters. However, further study is required to unravel the role and distribution of such complex and unusual lipids in bacteria.

*Ancylomarina* sp. strain M2P and *Labilibaculum* sp. strain SYP, contained *iso*-$C_{15:0}$ as the most abundant core lipid, accounting for 33.6% and 34.4% respectively, followed by *iso*-$C_{17:1}\omega 8$, *iso*-$C_{15:1}\omega 8$, and other fatty acids in smaller relative abundances (Table S47). In a previous study on *Ancylomarina* and *Labilibaculum* spp., we observed that these species produced a wide range of phosphate-free polar headgroups in their intact polar lipids, such as ornithine lipids (OLs), capnine lipids (CpL) (i.e., sulfur-containing lipids), flavolipins (FL), and glycine lipids (GlyL)[4]. Marine microorganisms are known to use phosphate-free head groups as a survival strategy to cope with phosphate-limiting conditions[48,49]. They can substitute e.g. aminolipids in place of phospholipids in their cell membranes, which allows them to conserve phosphorus, a critical nutrient that is often scarce in marine environments[50]. These lipids can replace phospholipids in the cell membrane without compromising membrane integrity and may even enhance membrane stability and resistance to environmental stressors[50,51]. Furthermore, OLs have been shown to enhance the uptake of phosphorus and other nutrients, potentially providing a competitive advantage to bacteria under P-limited conditions[4,48,51]. Moreover, OLs containing cell membranes are less susceptible to antibiotics and antimicrobial peptides[51] which could ultimately protect against bacterial predators (antibiotic producing microorganisms).

*Iso*-$C_{15:0}$, *anteiso*-$C_{15:0}$, $C_{16:0}$, *iso*-$C_{17:0}$ and $C_{18:0}$ fatty acids were present as major core lipids (>5%) in *Pseudodesulfovibrio* sp. strain S3$^T$ (Table S48). Other notable fatty acids present in these strains included, *iso*-$C_{17:1\omega8}$, *anteiso*-$C_{17:0}$, *iso*-cy$C_{18:0}$, and *iso*-$C_{19:1\omega8}$. The presence of these fatty acids might play important role in their adaptation to elevated hydrostatic pressure since the presence of these fatty acids can impact the fluidity of the cell membrane by impacting the melting temperature[1,4,43,44].

*Iso*-$C_{15:0}$ was the predominant fatty acid in *Lutibacter* sp. strains B1$^T$, B2[52], and *Lutibacter profundi* DSM 100437$^T$ (this study), with relative abundances of 29%, 27.6%, and 28.1%, respectively. Other fatty acids include the *iso*-$C_{13:0}$ and *iso*-$C_{15:1\omega11c}$, which were also present in relatively high abundances. Hydroxy fatty acids were also present in significant amounts, with *iso*-$C_{15:0}$ *β*-OH FA being the most abundant in all three strains (Table S49). Glycine *β*-OH fatty acid amides were present in smaller amounts, with glycine *iso*-*β*-OH $C_{17:0}$ amide being the most abundant in strains B1$^T$ and B2. The high proportion of amino acid lipids in *Lutibacter* sp. strains B1$^T$ and B2 could be due to their adaptation to the conditions of the Black Sea's deep sulfidic waters.

Overall, the lipid results suggest that the novel taxa isolated in this study have adapted to their environment by incorporating specific types of core lipids, hydroxy fatty acids, and glycerol-free amino lipids such as GlyLs into their cell membranes. However, further study is required to better constrain the role of these lipids in adapting to the triple extreme conditions [(i.e., recalcitrant nutrient condition (sulfurized OM), elevated hydrostatic pressure, and higher sulfide concentrations)] of the deep sulfidic waters of the Black Sea.

**References:**


1. Yadav, S., Koenen, M., Bale, N., Sinninghe Damsté, J. S. & Villanueva, L. The physiology and metabolic properties of a novel, low-abundance Psychrilyobacter species isolated from the anoxic Black Sea shed light on its ecological role. *Env. Microbiol. Rep.* **13**, 899–910 (2021).
2. Suominen, S., Dombrowski, N., Sinninghe Damsté, J. S. & Villanueva, L. A diverse uncultivated microbial community is responsible for organic matter degradation in the Black Sea sulfidic zone. *Env. Microbiol.* **23**, 2709–2728 (2021).
3. Suominen, S., Doorenspleet, K., Sinninghe Damsté, J. S. & Villanueva, L. Microbial community development on model particles in the deep sulfidic waters of the Black Sea. *Env. Microbiol.* **23**, 2729–2746 (2021).
4. Yadav, S. *et al.* Physiological, chemotaxonomic and genomic characterization of two novel piezotolerant bacteria of the family Marinifilaceae isolated from sulfidic waters of the Black Sea. *Syst. Appl. Microbiol.* **43**, (2020).
5. Davis, K. E., Joseph, S. J. & Janssen, P. H. Effects of growth medium, inoculum size, and incubation time on culturability and isolation of soil bacteria. *Appl. Env. Microbiol.* **71**, 826–34 (2005).
6. Connon, S. A. & Giovannoni, S. J. High-throughput methods for culturing microorganisms in very-low-nutrient media yield diverse new marine isolates. *Appl Env. Microbiol.* **68**, 3878–3885 (2002).

7. Zengler, K. *et al.* Cultivating the uncultured. *Proc. Natl. Acad. Sci. USA.* **99**, 15681–15686 (2002).

8. Greening, C. *et al.* Genomic and metagenomic surveys of hydrogenase distribution indicate $H_2$ is a widely utilized energy source for microbial growth and survival. *ISME J.* **10**, 761–777 (2016).

9. Ramel, F. *et al.* Growth of the Obligate Anaerobe Desulfovibrio vulgaris Hildenborough under Continuous Low Oxygen Concentration Sparging: Impact of the Membrane Bound Oxygen Reductases. *PLoS ONE* **10**, 0123455 (2015).

10. Mardanov, A. V. *et al.* Genomic insights into a new acidophilic, copper-resistant Desulfosporosinus isolate from the oxidized tailings area of an abandoned gold mine. *FEMS Microbiol. Ecol.* **92**, 111 (2016).

11. Pereira, I. A. C., Romão, C. V., Xavier, A. V., LeGall, J. & Teixeira, M. Electron transfer between hydrogenases and mono and multiheme cytochromes in Desulfovibrio spp. *J. Biol. Inorg. Chem.* **3**, 494–498 (1998).

12. Matias, P. M., Pereira, I. A., Soares, C. M. & Carrondo, M. A. Sulphate respiration from hydrogen in Desulfovibrio bacteria: a structural biology overview. *Prog. Biophys. Mol. Biol.* **89**, 292–329 (2005).

13. Rossi, M. *et al.* The hmc operon of Desulfovibrio vulgaris subsp. vulgaris Hildenborough encodes a potential transmembrane redox protein complex. *J. Bacteriol.* **175**, 4699–4711 (1993).

14. Thauer, R. K., Kaster, A. K., Seedorf, H., Buckel, W. & Hedderich, R. Methanogenic archaea: ecologically relevant differences in energy conservation. *Nat. Rev. Microbiol.* **6**, 579–591 (2008).

15. Vignais, P. M. & Billoud, B. Occurrence, classification, and biological function of hydrogenases: an overview. *Chem. Rev.* **107**, 4206–4272 (2007).

16. Rothery, R. A., Workun, G. J. & Weiner, J. H. The prokaryotic complex iron-sulfur molybdoenzyme family. *Biochim. Biophys. Acta.* **1778**, 1897–1929 (2008).

17. Rodrigues, M. L., Oliveira, T. F., Pereira, I. A. & Archer, M. X-ray structure of the membrane-bound cytochrome c quinol dehydrogenase NrfH reveals novel haem coordination. *EMBO J.* **25**, 5951–5960 (2006).

18. Greene, E. A., Hubert, C., Nemati, M., Jenneman, G. E. & Voordouw, G. Nitrite reductase activity of sulfate-reducing bacteria prevents their inhibition by nitrate-reducing, sulfide-oxidizing bacteria. *Env. Microbiol.* **5**, 607–617 (2003).

19. Wasmund, K. *et al.* Genomic insights into diverse bacterial taxa that degrade extracellular DNA in marine sediments. *Nat. Microbiol.* **6**, 885–898 (2021).

20. Zheng, R. *et al.* Characterization of the first cultured free-living representative of Candidatus Izemoplasma uncovers its unique biology. *ISME J.* **15**, 2676–2691 (2021).

21. Villanueva, L. *et al.* Bridging the membrane lipid divide: bacteria of the FCB group superphylum have the potential to synthesize archaeal ether lipids. *ISME J.* **15**, 168–182 (2021).

22. Johnson, L. A. & Hug, L. A. Cloacimonadota metabolisms include adaptations in engineered environments that are reflected in the evolutionary history of the phylum. *Env. Microbiol. Rep.* **14**, 520–529 (2022).

23. Chouari, R. *et al.* Novel predominant archaeal and bacterial groups revealed by molecular analysis of an anaerobic sludge digester. *Env. Microbiol.* **7**, 1104–1115 (2005).

24. Solli, L., Håvelsrud, O. E., Horn, S. J. & Rike, A. G. A metagenomic study of the microbial communities in four parallel biogas reactors. *Biotechnol. Biofuels* **7**, (2014).

25. Ahlert, S., Zimmermann, R., Ebling, J. & König, H. Analysis of propionate-degrading consortia from agricultural biogas plants. *Microbiology* **5**, 1027–1037 (2016).

26. Westerholm, M. *et al.* Microbial community dynamics linked to enhanced substrate availability and biogas production of electrokinetically pre-treated waste activated sludge. *Bioresour. Technol.* **218**, 761–770 (2016).

27. Calusinska, M. *et al.* A year of monitoring 20 mesophilic full-scale bioreactors reveals the existence of stable but different core microbiomes in bio-waste and wastewater anaerobic digestion systems. *Biotechnol. Biofuels* **11**, (2018).

28. Jankowska, E., Duber, A., Chwialkowska, J., Stodolny, M. & Oleskowicz-Popiel, P. Conversion of organic waste into volatile fatty acids - the influence of process operating parameters. *Chem. Eng. J.* **345**, 395–403 (2018).

29. Theuerl, S., Klang, J., Heiermann, M. & Vrieze, J. Marker microbiome clusters are determined by operational parameters and specific key taxa combinations in anaerobic digestion. *Bioresour. Technol.* **263**, 128–135 (2018).

30. S, S. Y. *et al.* Sulfide level in municipal sludge digesters affects microbial community response to long-chain fatty acid loads. *Biotechnol. Biofuels* **12**, (2019).

31. Drickamer, K. & Taylor, M. E. Evolving views of protein glycosylation. *Trends Biochem. Sci.* **23**, 321–324 (1998).

32. Hess, V. *et al.* Occurrence of ferredoxin: NAD+ oxidoreductase activity and its ion specificity in several gram-positive and gram-negative bacteria. *PeerJ.* **4:e1515**, (2016).

33. Feng, X., Tang, K. H., Blankenship, R. E. & Tang, Y. J. Metabolic flux analysis of the mixotrophic metabolisms in the green sulfur bacterium Chlorobaculum tepidum. *J. Biol. Chem.* **285**, 39544–39550 (2010).

34. Wahlund, T. M. & Tabita, F. R. The reductive tricarboxylic acid cycle of carbon dioxide assimilation: initial studies and purification of ATP-citrate lyase from the green sulfur bacterium Chlorobium tepidum. *J. Bacteriol.* **179**, 4859–4867 (1997).

35. Meyer, M., Dimroth, P. & Bott, M. Catabolite repression of the citrate fermentation genes in Klebsiella pneumoniae: evidence for involvement of the cyclic AMP receptor protein. *J. Bacteriol.* **183**, 5248–5256 (2001).

36. Biegel, E. & Müller, V. Bacterial Na+-translocating ferredoxin:NAD+ oxidoreductase. *Proc. Natl. Acad. Sci. USA.* **107**, 18138–18142 (2010).

37. García-Horsman, J. A., Barquera, B., Rumbley, J., Ma, J. & Gennis, R. B. The superfamily of heme-copper respiratory oxidases. *J. Bacteriol.* **176**, 5587–5600 (1994).

38. van der Oost, J. *et al.* The heme-copper oxidase family consists of three distinct types of terminal oxidases and is related to nitric oxide reductase. *FEMS Microbiol. Lett.* **121**, 1–9 (1994).

39. Seshadri, R. *et al.* Genome sequence of the PCE-dechlorinating bacterium Dehalococcoides ethenogenes. *Science* **307**, 105–108 (2005).

40. Sewell, H. L., Kaster, A. K. & Spormann, A. M. Homoacetogenesis in Deep-Sea Chloroflexi, as Inferred by Single-Cell Genomics, Provides a Link to Reductive Dehalogenation in Terrestrial Dehalococcoidetes. *mBio.* **8:e02022-17**, (2017).

41. Kaster, A. K., Mayer-Blackwell, K., Pasarelli, B. & Spormann, A. M. Single cell genomic study of Dehalococcoidetes species from deep-sea sediments of the Peruvian Margin. *ISME J.* **8**, 1831–1842 (2014).

42. Buckel, W. & Thauer, R. K. Flavin-Based Electron Bifurcation, A New Mechanism of Biological Energy Coupling. *Chem. Rev.* **118**, 3862–3886 (2018).

43. Ernst, R., Ejsing, C. S. & Antonny, B. Homeoviscous adaptation and the regulation of membrane lipids. *J. Mol. Biol.* **428**, 4776–4791 (2016).

44. Nichols, D. S. *et al.* Cold adaptation in the Antarctic Archaeon Methanococcoides burtonii involves membrane lipid unsaturation. *J. Bacteriol.* **186**, 8508–8515 (2004).

45. Rush, D. & Sinninghe Damsté, J. S. Lipids as paleomarkers to constrain the marine nitrogen cycle. *Env. Microbiol.* **19**, 2119–2132 (2017).

46. Bauersachs, T. *et al.* Distribution of heterocyst glycolipids in cyanobacteria. *Phytochem.* **70**, 2034–2039 (2009).

47. Zhang, Y. M. & Rock, C. O. Membrane lipid homeostasis in bacteria. *Nat. Rev. Microbiol.* **6**, 222–233 (2008).

48. Carini, P. *et al.* SAR11 lipid renovation in response to phosphate starvation. *Proc. Natl. Acad. Sci. USA.* **112**, 7767–7772 (2015).

49. Vences-Guzmán, M. Á., Geiger, O. & Sohlenkamp, C. Ornithine lipids and their structural modifications: from A to E and beyond. *FEMS Microbiol. Lett.* **335**, 1–10 (2012).

50. Sebastián, M. *et al.* Lipid remodelling is a widespread strategy in marine heterotrophic bacteria upon phosphorus deficiency. *ISME J.* **10**, 968–978 (2016).

51. Kim, S. K. *et al.* Bacterial ornithine lipid, a surrogate membrane lipid under phosphate-limiting conditions, plays important roles in bacterial persistence and interaction with host. *Env. Microbiol.* **20**, 3992–4008 (2018).

52. Bale, N. J. *et al.* Diagnostic amide products of amino lipids detected in the microaerophilic bacteria Lutibacter during routine fatty acid analysis using gas chromatography. *Org. Geochem.* **144**, (2020).

### Supplementary Figures

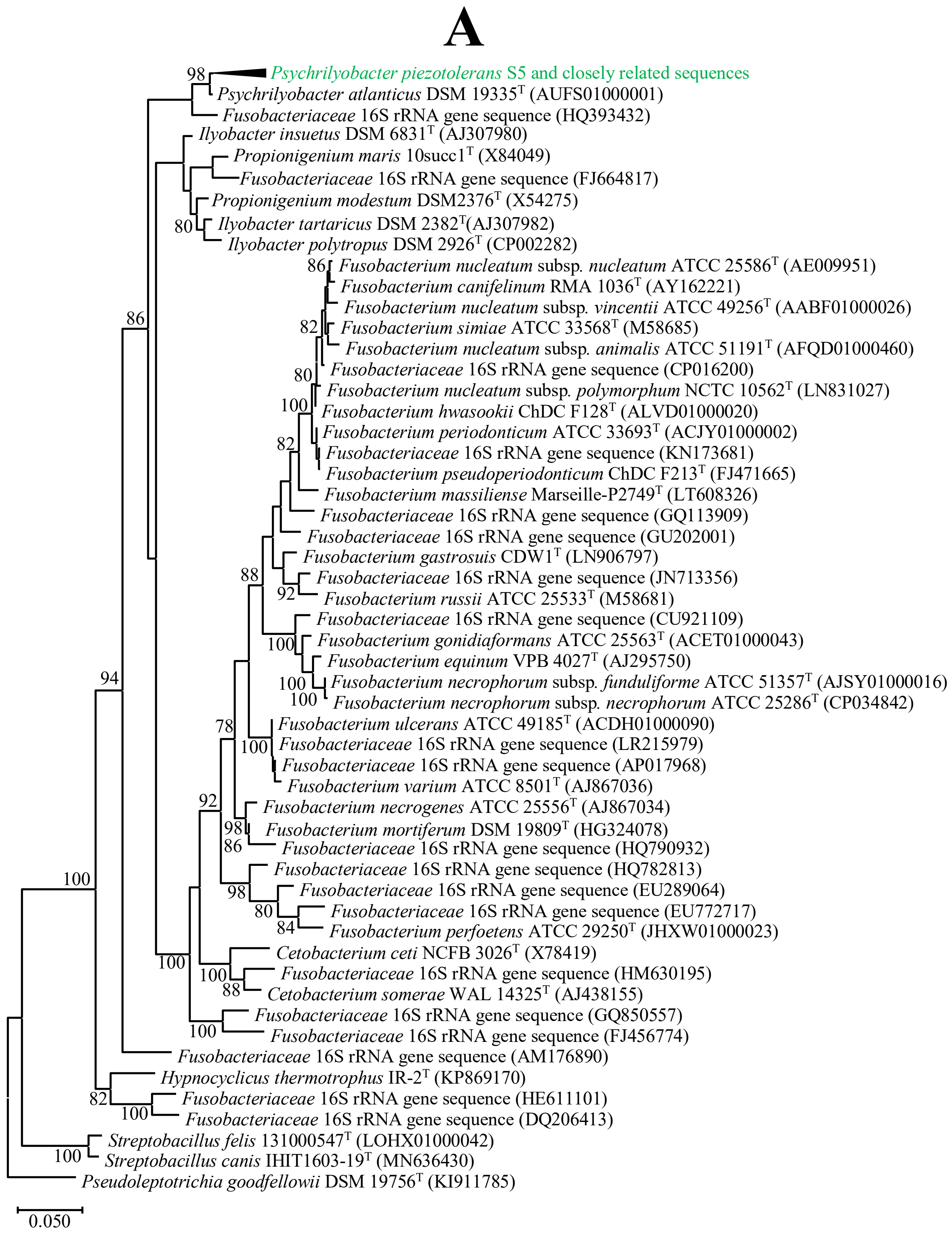


**Fig. S1: (A)** Phylogenetic tree based on 16S rRNA gene sequences showing the relationship of strains S5, SD5$^T$ and BL5 within the phylum *Fusobacteriota*. The tree was reconstructed by the maximum-likelihood method using MEGAX software and was rooted by using the 16S rRNA gene sequence of *Pseudoleptotrichia goodfellowii* DSM 19756$^T$ (KI911785) as the outgroup. Numbers at nodes represent bootstrap value (percentages, based on 1000 resamplings). GenBank accession numbers for 16S rRNA gene sequences are shown between parentheses. Bar, indicated 5 nucleotide substitutions per 100 nucleotides.

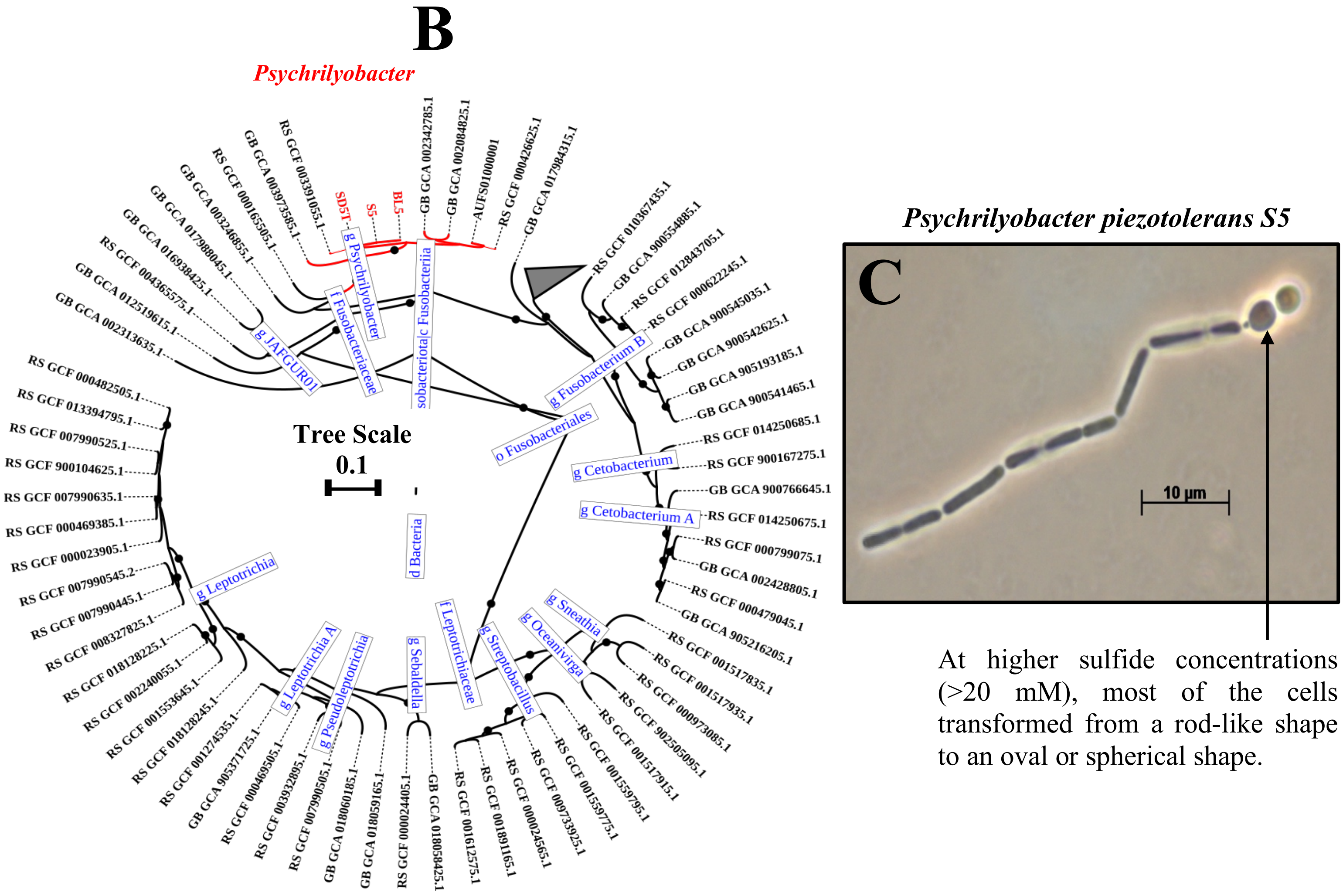


**Fig. S1: (B)** Subtrees of the GTDB-Tk 2.1.1 phylogenomic tree showing the affiliation of *Psychrilyobacter piezotolerans* strain S5, SD5$^{T}$ and BL5 (shown in red color) with other closely related members of the phylum *Fusobacteriota*. Class names are indicated by a leading "c_," order names by "o_," family names by "f_," and genus names by "g_,". The genome sequence accession numbers of the closely related taxa are shown. Black circle at nodes represents bootstrap value (100). The length of the bar indicates ten nucleotide substitutions per 100 nucleotides. **(C)** Cell morphology (phase contrast micrograph) of the *Psychrilyobacter piezotolerans* strain S5 grown at optimal growth conditions. Most of the cells appear as rod shape at optimal growth conditions while spherical cells are observed at increasing sulfide concentration.

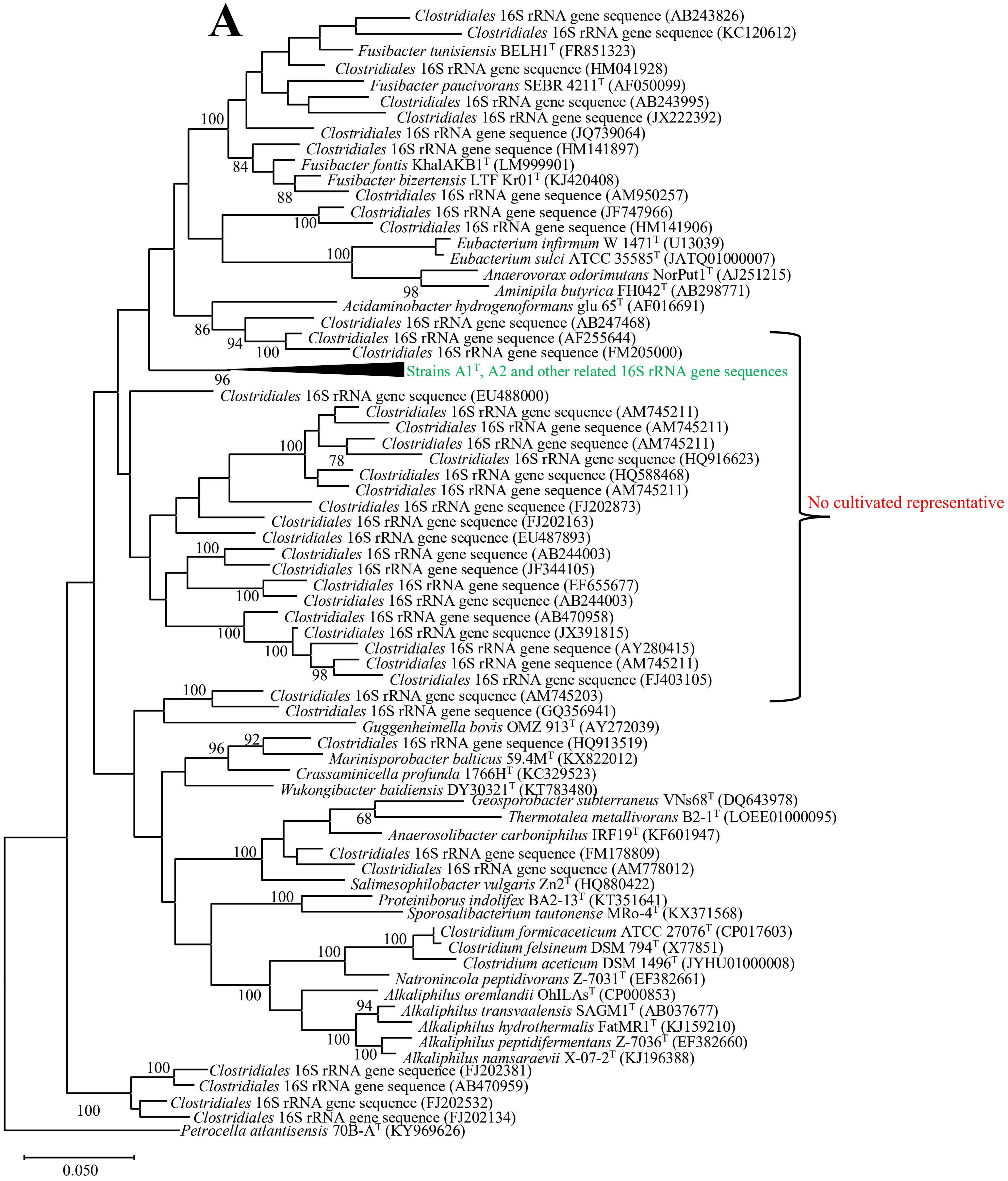
A
Clostridiales 16S rRNA gene sequence (AB243826)
Clostridiales 16S rRNA gene sequence (KC120612)
Fusibacter tunisiensis BELH1$^{T}$ (FR851323)
Clostridiales 16S rRNA gene sequence (HM041928)
Fusibacter paucivorans SEBR 4211$^{T}$ (AF050099)
Clostridiales 16S rRNA gene sequence (AB243995)
Clostridiales 16S rRNA gene sequence (JX222392)
100
Clostridiales 16S rRNA gene sequence (JQ739064)
Clostridiales 16S rRNA gene sequence (HM141897)
84
Fusibacter fontis KhalAKB1$^{T}$ (LM999901)
Fusibacter bizertensis LTF Kr01$^{T}$ (KJ420408)
88
Clostridiales 16S rRNA gene sequence (AM950257)
Clostridiales 16S rRNA gene sequence (JF747966)
100
Clostridiales 16S rRNA gene sequence (HM141906)
Eubacterium infirmum W 1471$^{T}$ (U13039)
100
Eubacterium sulci ATCC 35585$^{T}$ (JATQ01000007)
Anaerovorax odorimutans NorPut1$^{T}$ (AJ251215)
98
Aminipila butyrica FH042$^{T}$ (AB298771)
Acidaminobacter hydrogenoformans glu 65$^{T}$ (AF016691)
Clostridiales 16S rRNA gene sequence (AB247468)
86
Clostridiales 16S rRNA gene sequence (AF255644)
94
100
Clostridiales 16S rRNA gene sequence (FM205000)
Strains A1$^{T}$, A2 and other related 16S rRNA gene sequences
96
Clostridiales 16S rRNA gene sequence (EU488000)
Clostridiales 16S rRNA gene sequence (AM745211)
Clostridiales 16S rRNA gene sequence (AM745211)
100
Clostridiales 16S rRNA gene sequence (AM745211)
78
Clostridiales 16S rRNA gene sequence (HQ916623)
Clostridiales 16S rRNA gene sequence (HQ588468)
Clostridiales 16S rRNA gene sequence (AM745211)
No cultivated representative
Clostridiales 16S rRNA gene sequence (FJ202873)
Clostridiales 16S rRNA gene sequence (FJ202163)
Clostridiales 16S rRNA gene sequence (EU487893)
100
Clostridiales 16S rRNA gene sequence (AB244003)
Clostridiales 16S rRNA gene sequence (JF344105)
Clostridiales 16S rRNA gene sequence (EF655677)
100
Clostridiales 16S rRNA gene sequence (AB244003)
Clostridiales 16S rRNA gene sequence (AB470958)
100
Clostridiales 16S rRNA gene sequence (JX391815)
100
Clostridiales 16S rRNA gene sequence (AY280415)
Clostridiales 16S rRNA gene sequence (AM745211)
98
Clostridiales 16S rRNA gene sequence (FJ403105)
100
Clostridiales 16S rRNA gene sequence (AM745203)
Clostridiales 16S rRNA gene sequence (GQ356941)
Guggenheimella bovis OMZ 913$^{T}$ (AY272039)
92
96
Clostridiales 16S rRNA gene sequence (HQ913519)
Marinisporobacter balticus 59.4M$^{T}$ (KX822012)
Crassaminicella profunda 1766H$^{T}$ (KC329523)
Wukongibacter baidiensis DY30321$^{T}$ (KT783480)
Geosporobacter subterraneus VNs68$^{T}$ (DQ643978)
68
Thermotalea metallivorans B2-1$^{T}$ (LOEE01000095)
Anaerosolibacter carboniphilus IRF19$^{T}$ (KF601947)
100
Clostridiales 16S rRNA gene sequence (FM178809)
Clostridiales 16S rRNA gene sequence (AM778012)
Salimesophilobacter vulgaris Zn2$^{T}$ (HQ880422)
100
Proteiniborus indolifex BA2-13$^{T}$ (KT351641)
Sporosalibacterium tautonense MRo-4$^{T}$ (KX371568)
100
Clostridium formicaceticum ATCC 27076$^{T}$ (CP017603)
100
Clostridium felsineum DSM 794$^{T}$ (X77851)
Clostridium aceticum DSM 1496$^{T}$ (JYHU01000008)
Natronincola peptidivorans Z-7031$^{T}$ (EF382661)
100
Alkaliphilus oremlandii OhILAs$^{T}$ (CP000853)
94
Alkaliphilus transvaalensis SAGM1$^{T}$ (AB037677)
Alkaliphilus hydrothermalis FatMR1$^{T}$ (KJ159210)
100
Alkaliphilus peptidifermentans Z-7036$^{T}$ (EF382660)
100
Alkaliphilus namsaraevii X-07-2$^{T}$ (KJ196388)
100
Clostridiales 16S rRNA gene sequence (FJ202381)
Clostridiales 16S rRNA gene sequence (AB470959)
Clostridiales 16S rRNA gene sequence (FJ202532)
100
Clostridiales 16S rRNA gene sequence (FJ202134)
Petrocella atlantisensis 70B-A$^{T}$ (KY969626)
0.050

**Fig. S2: (A)** Phylogenetic tree based on 16S rRNA gene sequences showing the relationship of *Clostridiales* bacteria strains $A1^T$ and A2 within the phylum *Bacillota*. The tree was reconstructed by the maximum-likelihood method using MEGAX software and was rooted by using the 16S rRNA gene sequence of *Petrocella atlantisensis* 70B-$A^T$ (KY969626) as the outgroup. Numbers at nodes represent bootstrap value (percentages, based on 1000 resamplings). GenBank accession numbers for 16S rRNA gene sequences are shown in parentheses. Bar, 5 nucleotide substitutions per 100 nucleotides.

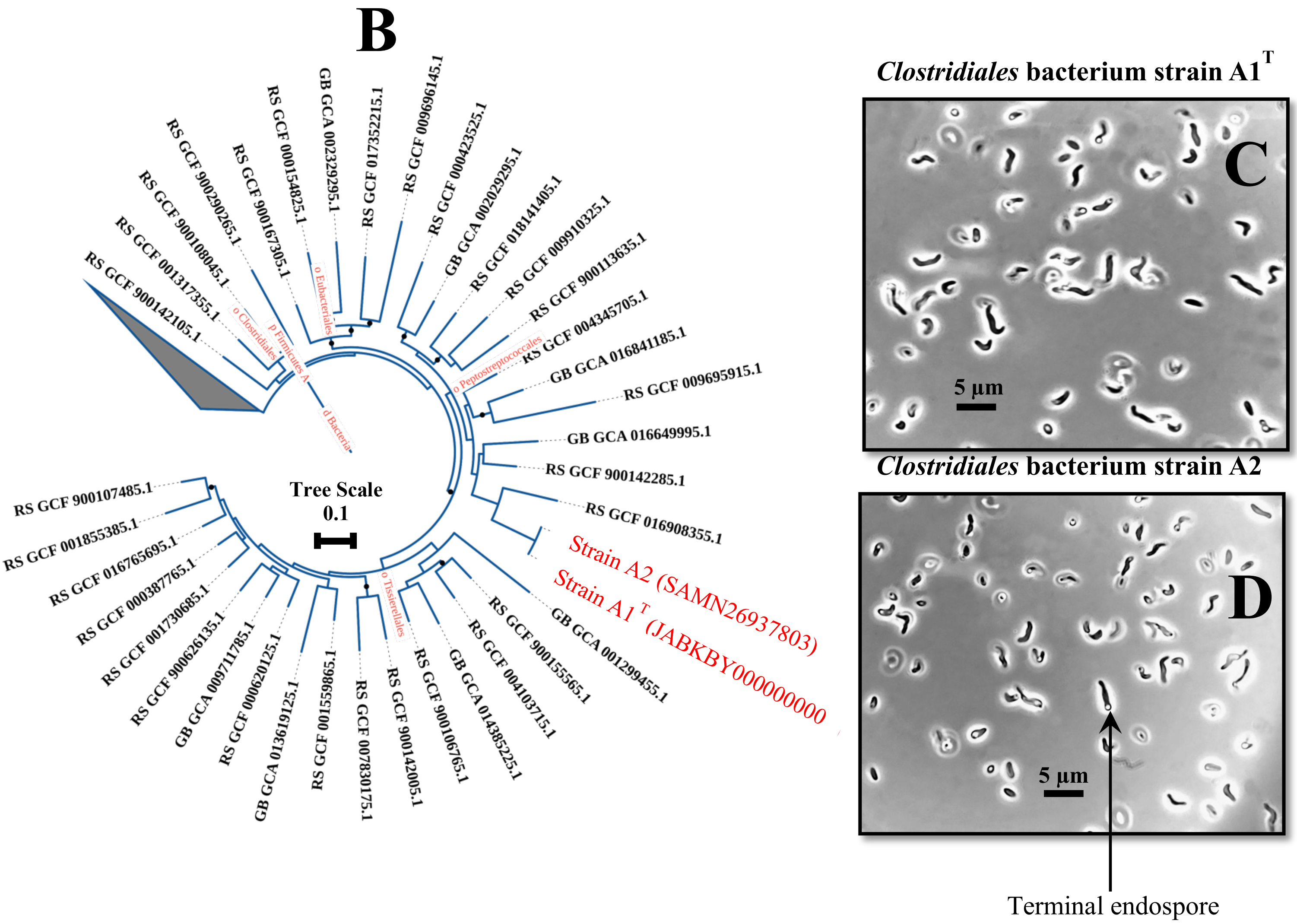


**Fig. S2: (B)** Subtrees of the GTDB-Tk 2.1.1 phylogenomic tree showing the affiliation of *Clostridiales* bacteria strains A1$^{T}$, and A2 (shown in red color) with other closely related members of the phylum *Bacillota*. Class names are indicated by a leading "c_," order names by "o_," family names by "f_," and genus names by "g_,". The genome sequence accession numbers of the closely related taxa are shown. Black circle at nodes represents bootstrap value (100). The length of the bar indicates ten nucleotide substitutions per 100 nucleotides. **(C)** Cell morphology (phase contrast micrograph) of the *Clostridiales* bacteria strains A1$^{T}$ **(C)**, and **(D)** A2 grown at optimal growth conditions. Terminal endospores are observed under phase contrast micrograph after 5 days of incubation at optimal growth conditions.

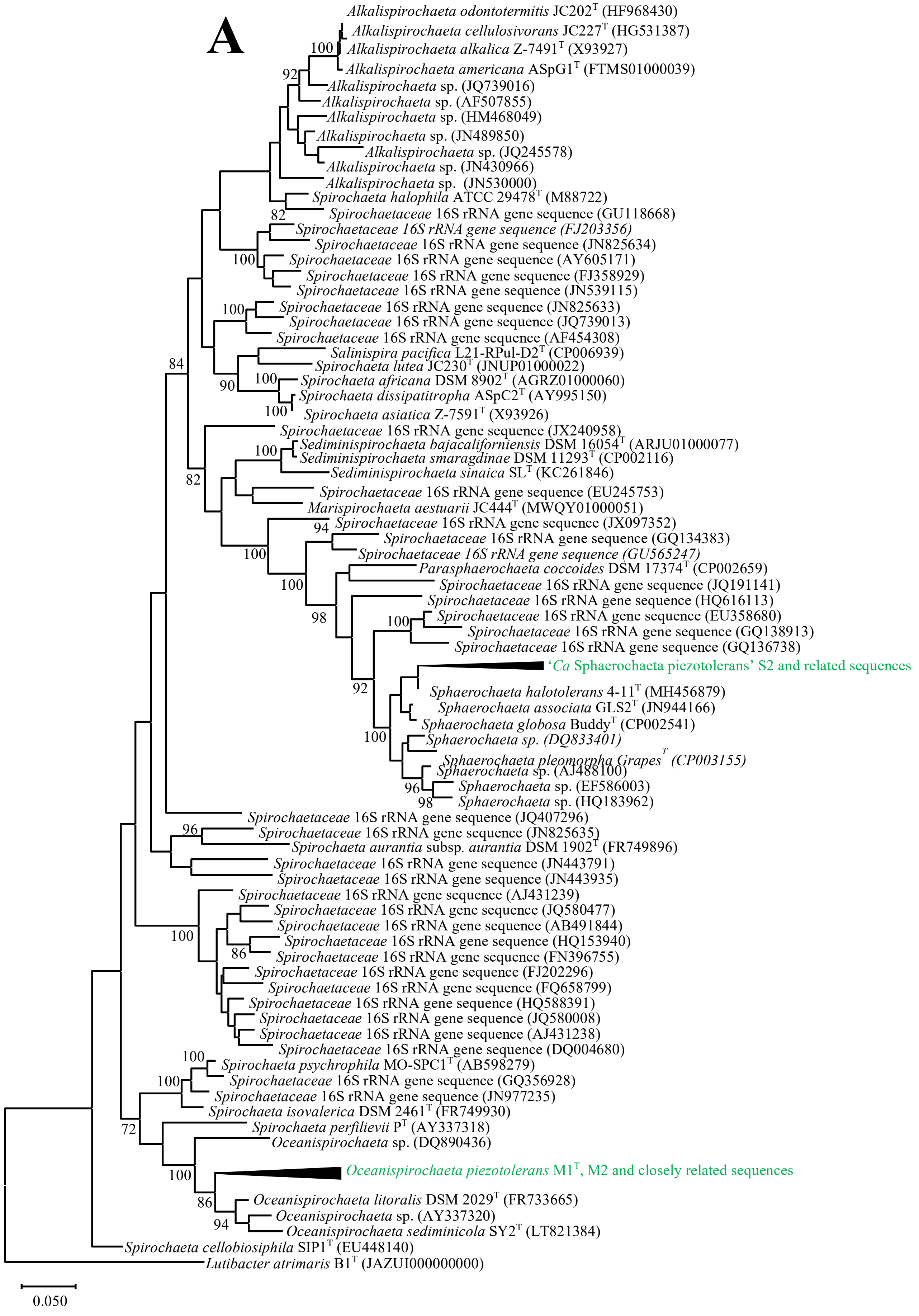
A
Alkalispirochaeta odontotermitis JC202T (HF968430)
Alkalispirochaeta cellulosivorans JC227T (HG531387)
Alkalispirochaeta alkalica Z-7491T (X93927)
Alkalispirochaeta americana ASpG1T (FTMS01000039)
Alkalispirochaeta sp. (JQ739016)
Alkalispirochaeta sp. (AF507855)
Alkalispirochaeta sp. (HM468049)
Alkalispirochaeta sp. (JN489850)
Alkalispirochaeta sp. (JQ245578)
Alkalispirochaeta sp. (JN430966)
Alkalispirochaeta sp. (JN530000)
Spirochaeta halophila ATCC 29478T (M88722)
Spirochaetaceae 16S rRNA gene sequence (GU118668)
Spirochaetaceae 16S rRNA gene sequence (FJ203356)
Spirochaetaceae 16S rRNA gene sequence (JN825634)
Spirochaetaceae 16S rRNA gene sequence (AY605171)
Spirochaetaceae 16S rRNA gene sequence (FJ358929)
Spirochaetaceae 16S rRNA gene sequence (JN539115)
Spirochaetaceae 16S rRNA gene sequence (JN825633)
Spirochaetaceae 16S rRNA gene sequence (JQ739013)
Spirochaetaceae 16S rRNA gene sequence (AF454308)
Salinispira pacifica L21-RPul-D2T (CP006939)
Spirochaeta lutea JC230T (JNUP01000022)
Spirochaeta africana DSM 8902T (AGRZ01000060)
Spirochaeta dissipatitropha ASpC2T (AY995150)
Spirochaeta asiatica Z-7591T (X93926)
Spirochaetaceae 16S rRNA gene sequence (JX240958)
Sediminispirochaeta bajacaliforniensis DSM 16054T (ARJU01000077)
Sediminispirochaeta smaragdinae DSM 11293T (CP002116)
Sediminispirochaeta sinaica SLT (KC261846)
Spirochaetaceae 16S rRNA gene sequence (EU245753)
Marispirochaeta aestuarii JC444T (MWQY01000051)
Spirochaetaceae 16S rRNA gene sequence (JX097352)
Spirochaetaceae 16S rRNA gene sequence (GQ134383)
Spirochaetaceae 16S rRNA gene sequence (GU565247)
Parasphaerochaeta coccoides DSM 17374T (CP002659)
Spirochaetaceae 16S rRNA gene sequence (JQ191141)
Spirochaetaceae 16S rRNA gene sequence (HQ616113)
Spirochaetaceae 16S rRNA gene sequence (EU358680)
Spirochaetaceae 16S rRNA gene sequence (GQ138913)
Spirochaetaceae 16S rRNA gene sequence (GQ136738)
'Ca Sphaerochaeta piezotolerans' S2 and related sequences
Sphaerochaeta halotolerans 4-11T (MH456879)
Sphaerochaeta associata GLS2T (JN944166)
Sphaerochaeta globosa BuddyT (CP002541)
Sphaerochaeta sp. (DQ833401)
Sphaerochaeta pleomorpha GrapesT (CP003155)
Sphaerochaeta sp. (AJ488100)
Sphaerochaeta sp. (EF586003)
Sphaerochaeta sp. (HQ183962)
Spirochaetaceae 16S rRNA gene sequence (JQ407296)
Spirochaetaceae 16S rRNA gene sequence (JN825635)
Spirochaeta aurantia subsp. aurantia DSM 1902T (FR749896)
Spirochaetaceae 16S rRNA gene sequence (JN443791)
Spirochaetaceae 16S rRNA gene sequence (JN443935)
Spirochaetaceae 16S rRNA gene sequence (AJ431239)
Spirochaetaceae 16S rRNA gene sequence (JQ580477)
Spirochaetaceae 16S rRNA gene sequence (AB491844)
Spirochaetaceae 16S rRNA gene sequence (HQ153940)
Spirochaetaceae 16S rRNA gene sequence (FN396755)
Spirochaetaceae 16S rRNA gene sequence (FJ202296)
Spirochaetaceae 16S rRNA gene sequence (FQ658799)
Spirochaetaceae 16S rRNA gene sequence (HQ588391)
Spirochaetaceae 16S rRNA gene sequence (JQ580008)
Spirochaetaceae 16S rRNA gene sequence (AJ431238)
Spirochaetaceae 16S rRNA gene sequence (DQ004680)
Spirochaeta psychrophila MO-SPC1T (AB598279)
Spirochaetaceae 16S rRNA gene sequence (GQ356928)
Spirochaetaceae 16S rRNA gene sequence (JN977235)
Spirochaeta isovalerica DSM 2461T (FR749930)
Spirochaeta perfilievii PT (AY337318)
Oceanispirochaeta sp. (DQ890436)
Oceanispirochaeta piezotolerans M1T, M2 and closely related sequences
Oceanispirochaeta litoralis DSM 2029T (FR733665)
Oceanispirochaeta sp. (AY337320)
Oceanispirochaeta sediminicola SY2T (LT821384)
Spirochaeta cellobiosiphila SIP1T (EU448140)
Lutibacter atrimaris B1T (JAZUI000000000)
0.050

**Fig. S3: (A)** Phylogenetic tree based on 16S rRNA gene sequences showing the relationship of *Oceanispirochaeta* sp. strains M1$^{T}$, M2 and ‘Ca Sphaerochaeta piezotolerans’ strain S2 within the phylum *Spirochaetota*. The tree was reconstructed by the maximum-likelihood method using MEGA X software and was rooted by using the 16S rRNA gene sequence of *Lutibacter atrimaris* B1$^{T}$ (JAZUI000000000) as the outgroup. Numbers at nodes represent bootstrap value (percentages, based on 1000 resamplings). GenBank accession numbers for 16S rRNA gene sequences are shown in parentheses. Bar, 5 nucleotide substitutions per 100 nucleotides.

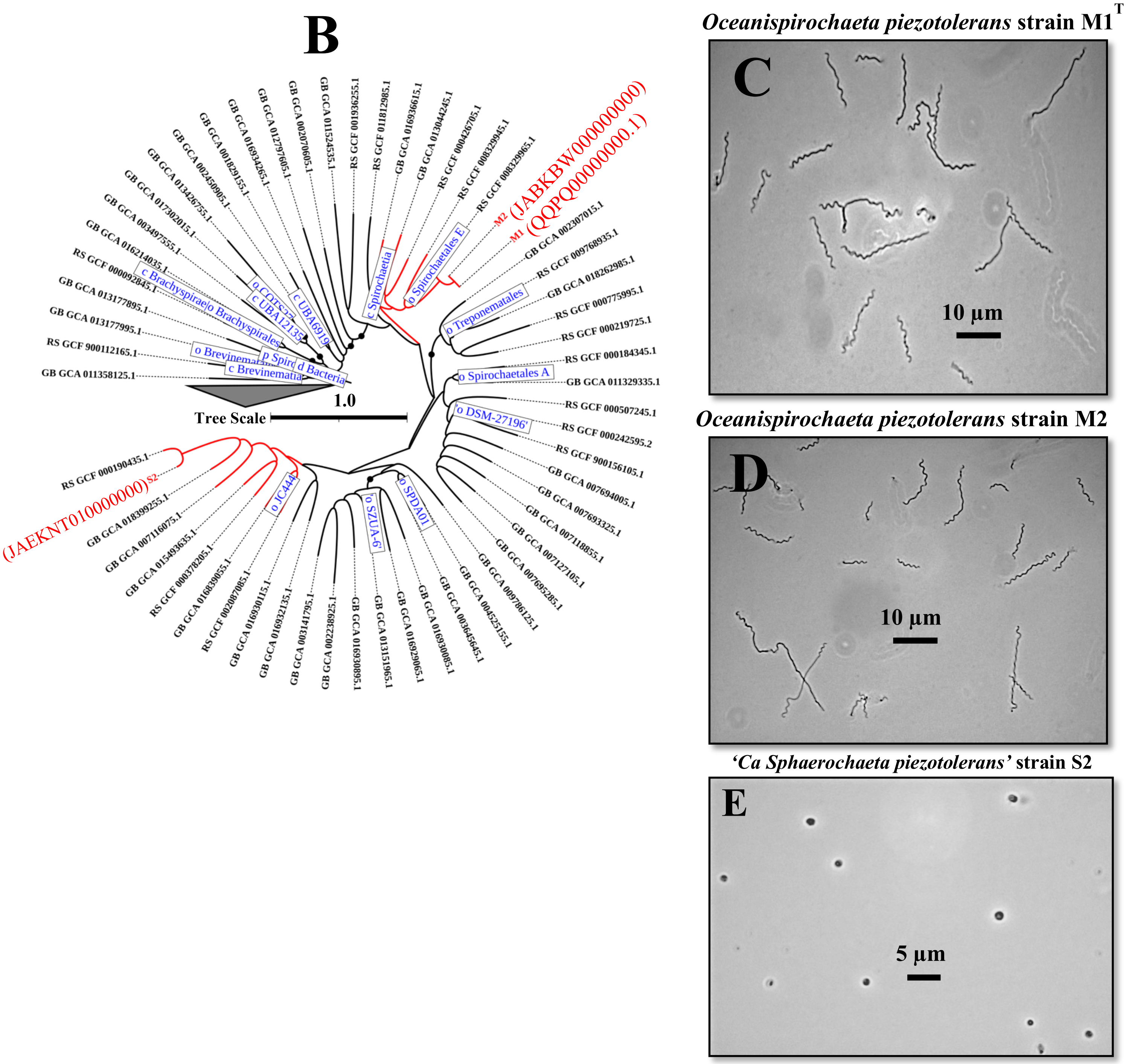


**Fig. S3: (B)** Subtrees of the GTDB-Tk 2.1.1 phylogenomic tree showing the affiliation of *Oceanispirochaeta piezotolerans* strains M1$^{T}$, M2 and '*Ca* Sphaerochaeta piezotolerans' strain S2 (shown in red color) with other closely related members of the phylum *Spirochaetota*. Class names are indicated by a leading "c_," order names by "o_," family names by "f_," and genus names by "g_,". The genome sequence accession numbers of the closely related taxa are shown. Black circle at nodes represents bootstrap value (100). The length of the bar indicates 100 nucleotide substitutions per 100 nucleotides. **(C)** Cell morphology (phase contrast micrograph) of the *Oceanispirochaeta piezotolerans* strains **(C)** M1$^{T}$, **(D)** M2 and **(E)** '*Ca* Sphaerochaeta piezotolerans' strain S2 grown at optimal growth conditions.

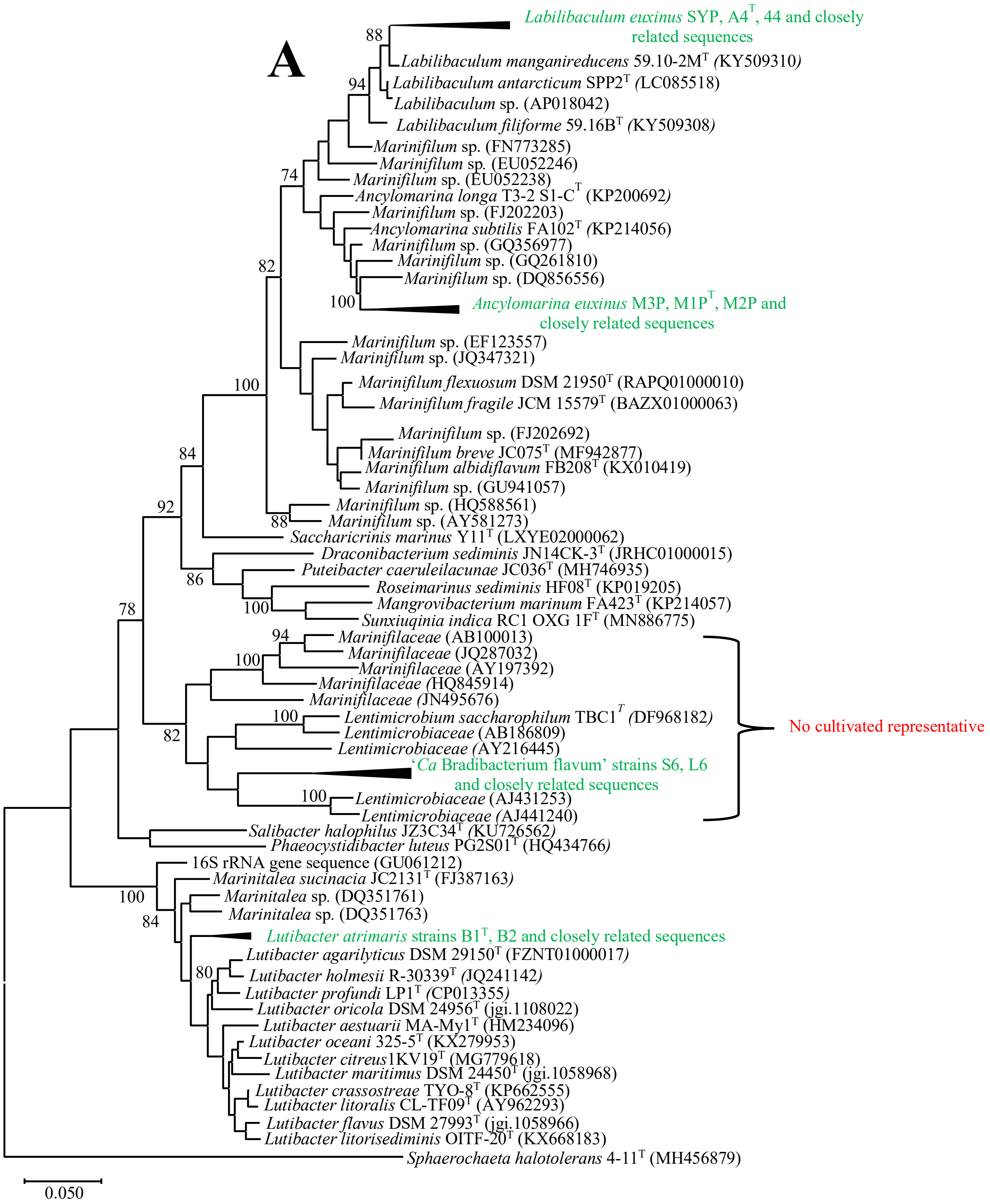


**Fig. S4: (A)** Phylogenetic tree based on 16S rRNA gene sequences showing the relationship of members of the phylum *Bacteroidota* (*Lutibacter* sp. strains B1$^T$, B2; 'Ca Bradibacterium flavum' strain S6, L6; *Ancylomarina euxinus* M2P; *Labilibaculum euxinus* SYP) isolated from deep sulfidic waters of the Black Sea. The tree was reconstructed by the maximum-likelihood method using MEGA X software and was rooted by using the 16S rRNA gene sequence of *Sphaerochaeta halotolerans* 4-11$^{T}$ (MH456879) as the outgroup. Numbers at nodes represent bootstrap value (percentages, based on 1000 resamplings). GenBank accession numbers for 16S rRNA gene sequences are shown in parentheses. Bar, 5 nucleotide substitutions per 100 nucleotides.

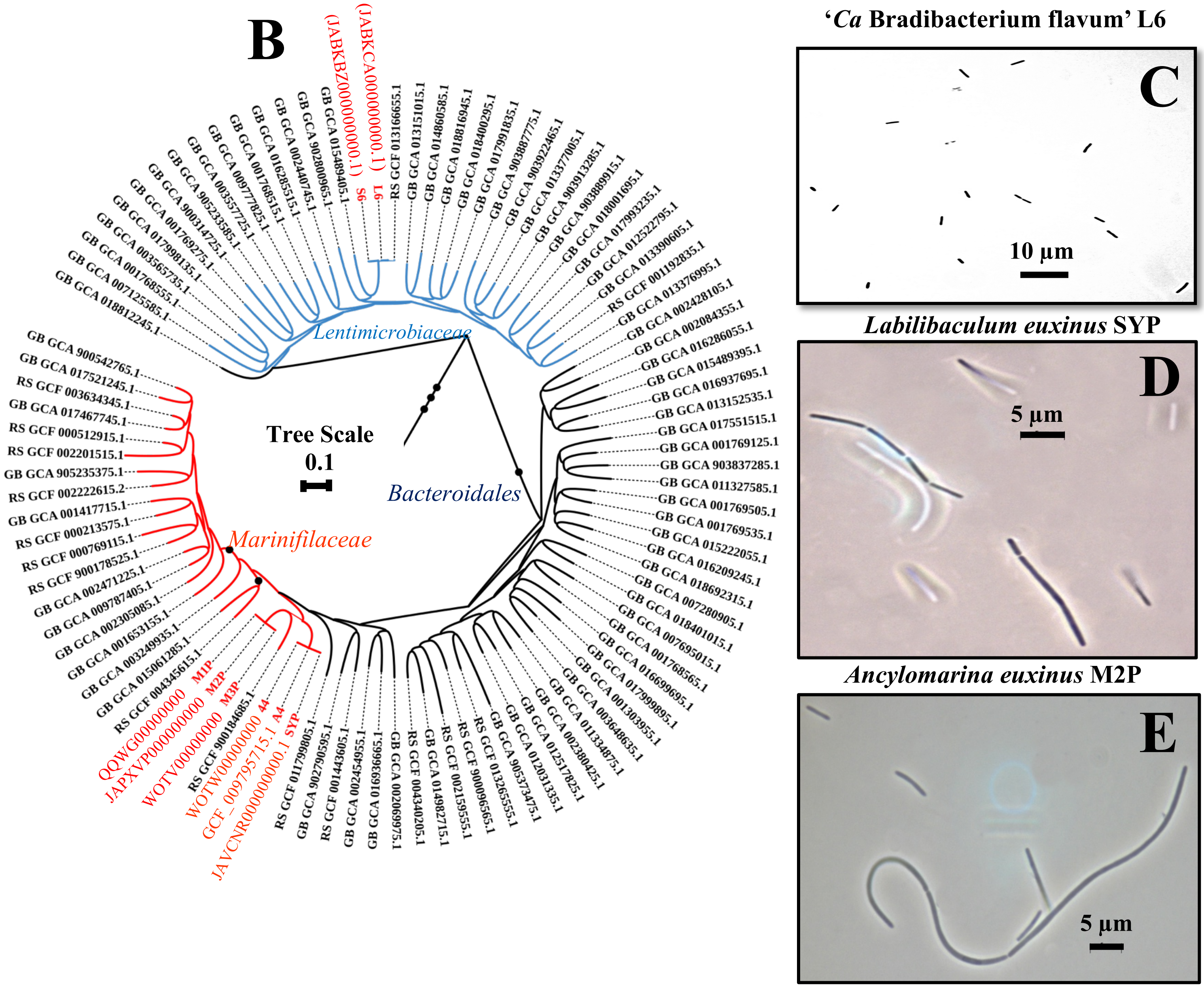


**Fig. S4: (B)** Subtrees of the GTDB-Tk 2.1.1 phylogenomic tree showing the affiliation of various members of the phylum *Bacteroidota* (‘Ca Bradibacterium flavum’ strains S6, L6; *Ancylomarina euxinus* strains M2P, M1P$^{T}$ and M3P; *Labilibaculum euxinus* strain SYP, A4$^{T}$ and 44 (shown in red color) with other closely related members of the phylum *Bacteroidota*. The genome sequence accession numbers of the closely related taxa are shown. Black circle at nodes represents bootstrap value (100). The length of the bar indicates ten nucleotide substitutions per 100 nucleotides. **(C)** Cell morphology (phase contrast micrograph) of the ‘*Ca* Bradibacterium flavum’ strain S6 **(D)** *Ancylomarina euxinus* strains M2P and **(E)** *Labilibaculum euxinus* strain SYP grown at optimal growth conditions.

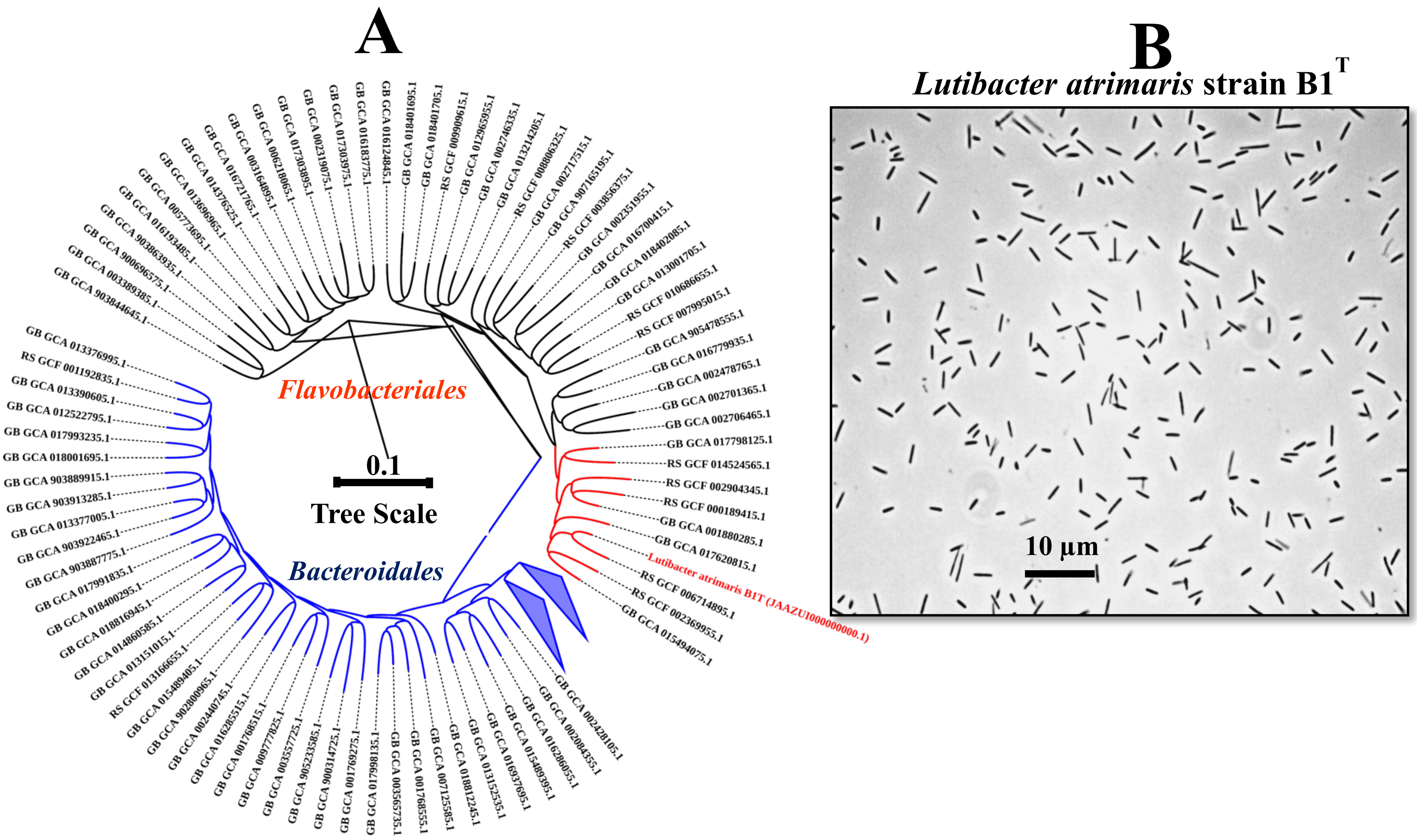


**Fig. S5: (A)** Subtrees of the GTDB-Tk 2.1.1 phylogenomic tree showing the affiliation of *Lutibacter atrimaris* strain B1$^{T}$ (shown in red color) with other closely related members of the phylum *Bacteroidota*. The genome sequence accession numbers of the closely related taxa are shown. The length of the bar indicates ten nucleotide substitutions per 100 nucleotides. **(B)** Cell morphology (phase contrast micrograph) of the *Lutibacter atrimaris* strain B1$^{T}$ grown at optimal growth conditions.

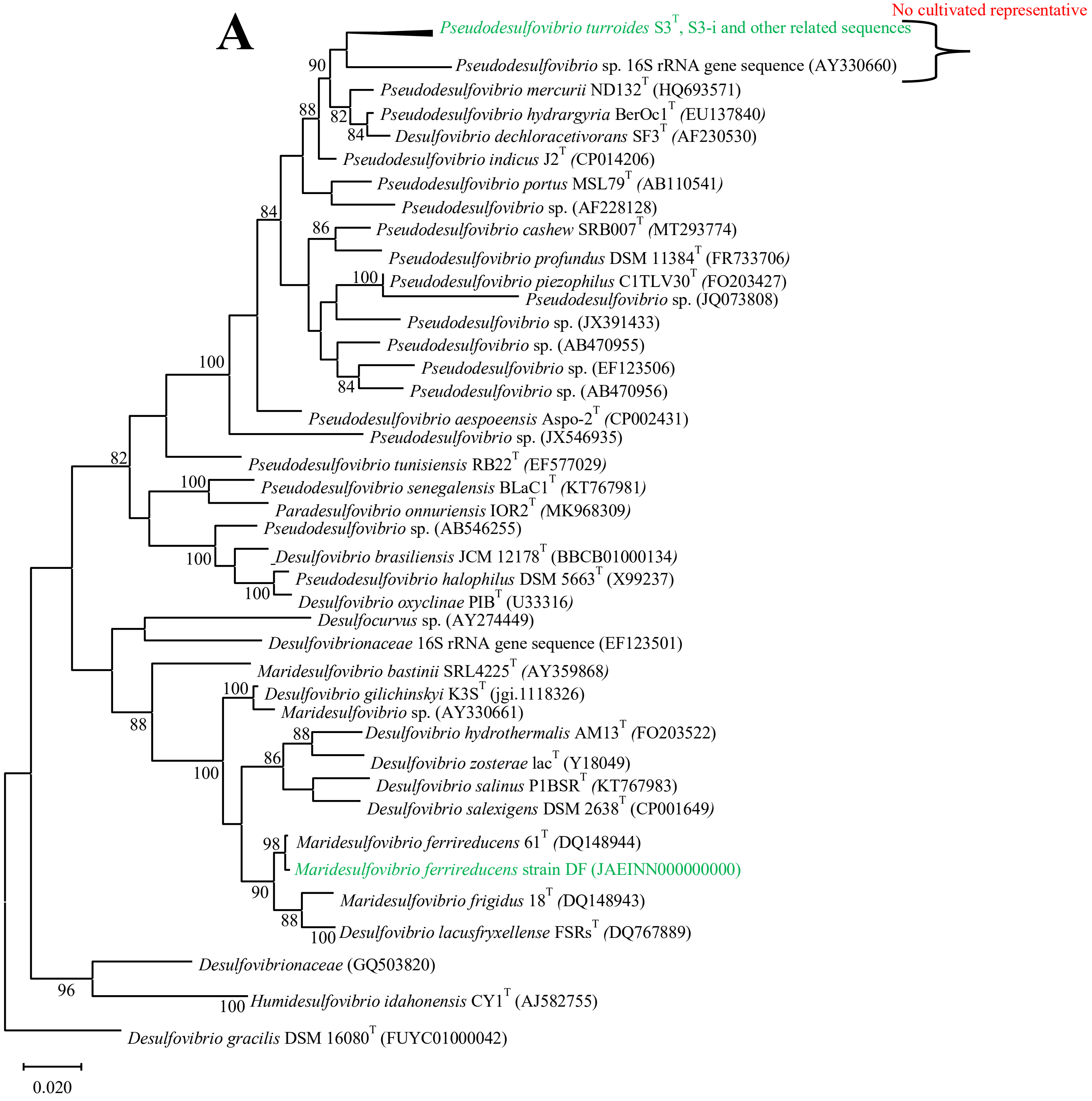


**Fig. S6: (A)** Phylogenetic tree based on 16S rRNA gene sequences showing the relationship of *Pseudodesulfovibrio* sp. strains S3$^T$, S3-i and other closely related members of the phylum *Desulfobacterota*. The tree was reconstructed by the maximum-likelihood method using MEGA X software and was rooted by using the 16S rRNA gene sequence of *Desulfovibrio gracilis* DSM 16080$^T$ (FUYC01000042) as the outgroup. Numbers at nodes represent bootstrap value (percentages, based on 1000 resamplings). GenBank accession numbers for 16S rRNA gene sequences are shown in parentheses. Bar, 2 nucleotide substitutions per 100 nucleotides.

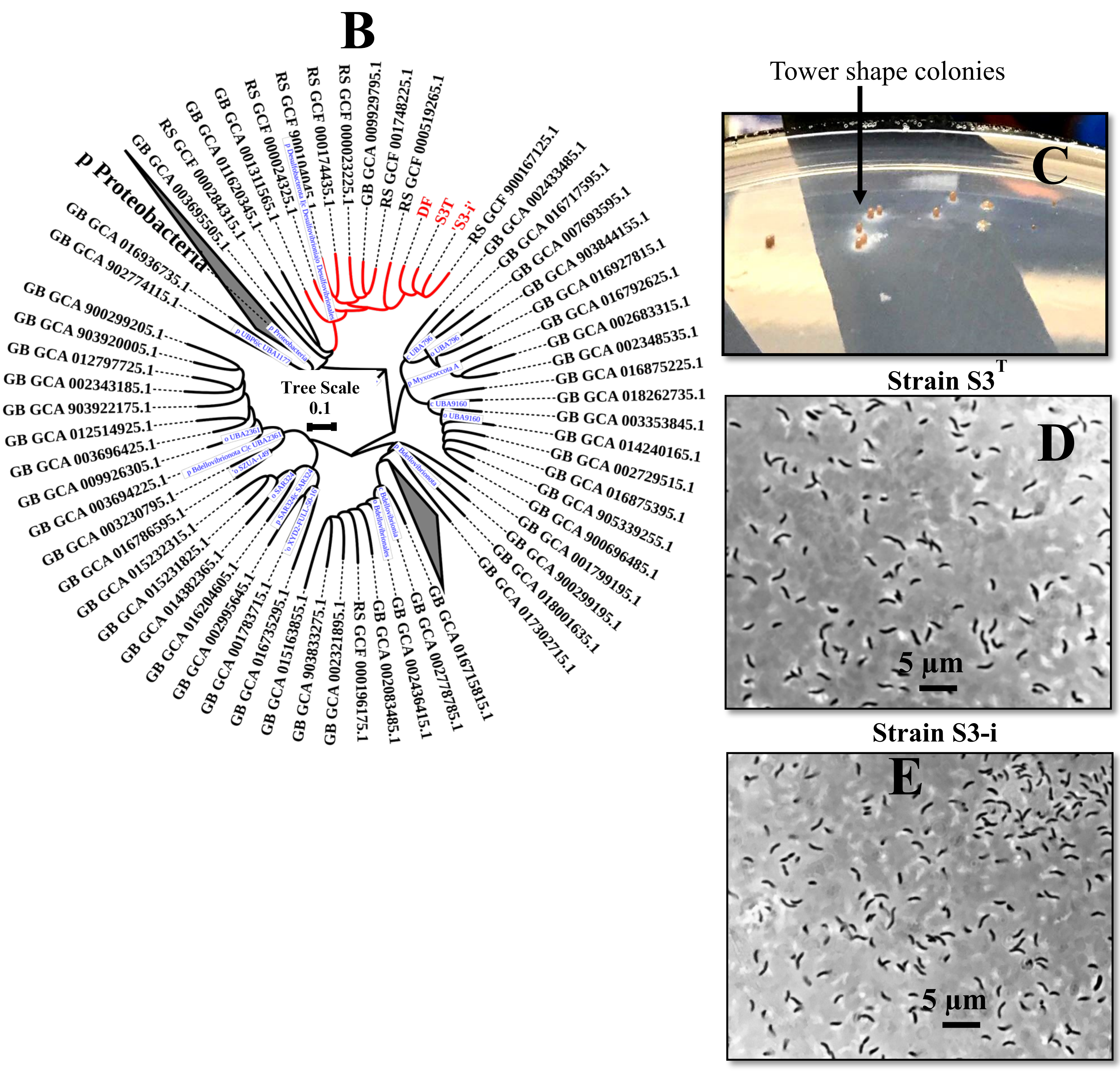


**Fig. S6: (B)** Subtrees of the GTDB-Tk 2.1.1 phylogenomic tree showing the affiliation of *Pseudodesulfovibrio turroides* strains S3$^T$, S3-i and *Maridesulfovibrio ferrireducens* strain DF (shown in red color) with other closely related members of the phylum *Desulfobacterota*. Class names are indicated by a leading "c_," order names by "o_," family names by "f_," and genus names by "g_,". The genome sequence accession numbers of the closely related taxa are shown. Black circle at nodes represents bootstrap value (100). The length of the bar indicates ten nucleotide substitutions per 100 nucleotides. **(C)** Orange colored tower shape colony morphology of strain S3$^T$ grown on agar medium (1.8%) after 1 month of incubation at 20 °C under strict anaerobic conditions at atmospheric pressure. **(D)** Cell morphology of *Pseudodesulfovibrio turroides* strains S3$^T$ **(E)** *Pseudodesulfovibrio turroides* strains S3-i grown at optimal growth conditions.

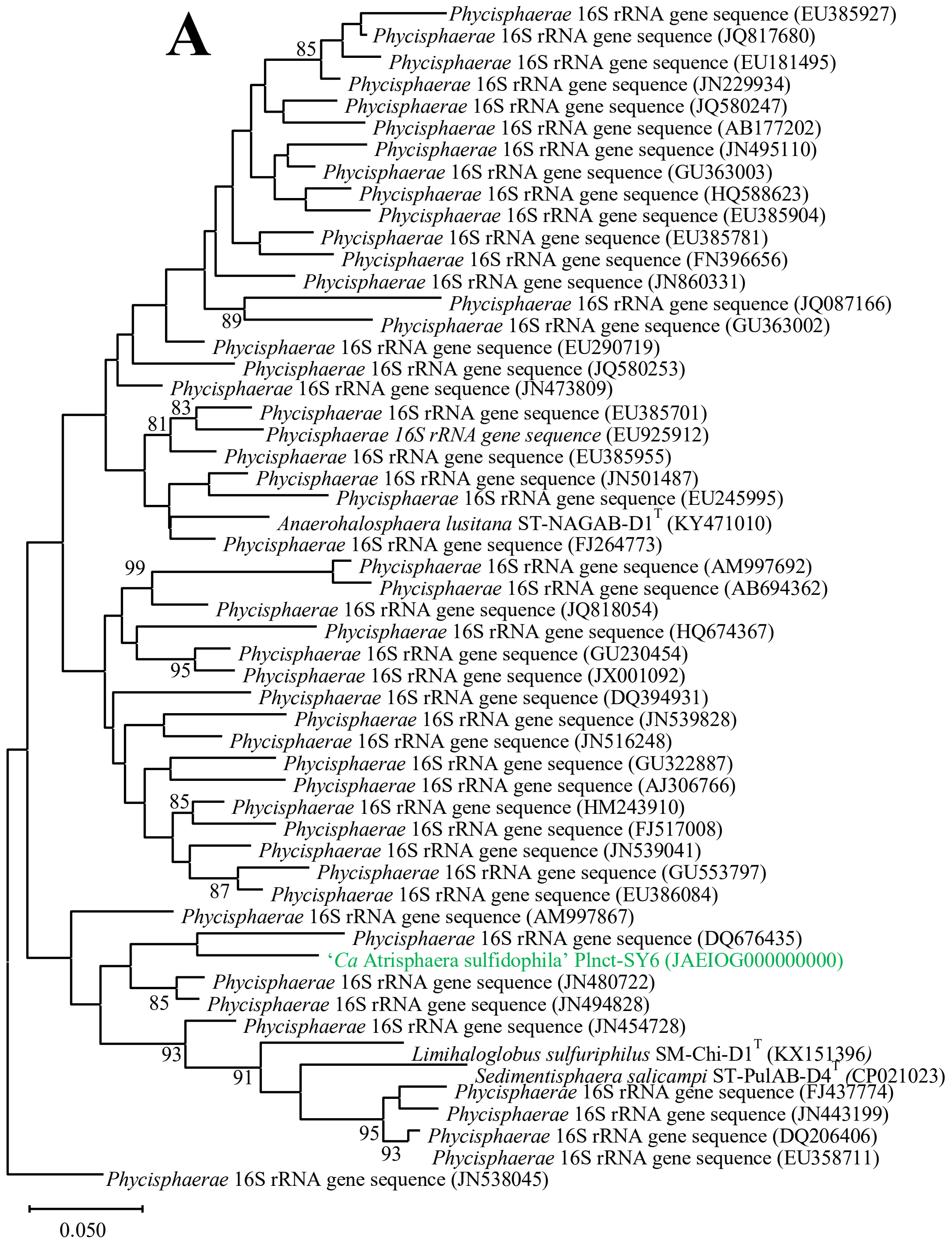


**Fig. S7: (A)** Phylogenetic tree based on 16S rRNA gene sequences showing the relationship of *'Ca* Atrisphaera sulfidophila' Plnct-SY6 and other closely related members of the phylum *Planctomycetota*. The tree was reconstructed by the maximum-likelihood method using MEGA X software and was rooted by using *Phycisphaerae* 16S rRNA gene sequence (JN538045) as the outgroup. Numbers at nodes represent bootstrap value (percentages, based on 1000 resamplings). GenBank accession numbers for 16S rRNA gene sequences are shown in parentheses. Bar, 5 nucleotide substitutions per 100 nucleotides.

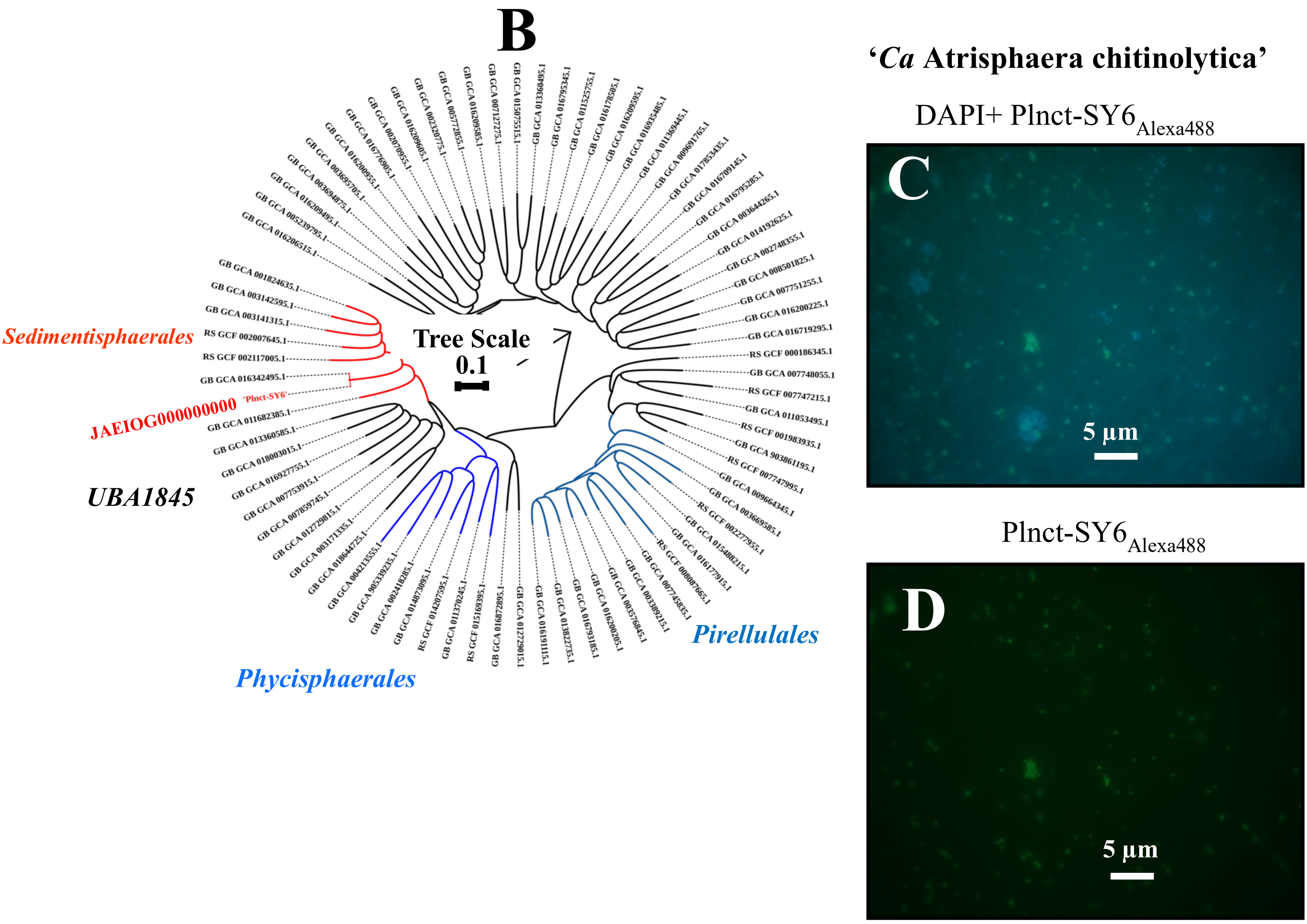


**Fig. S7: (B)** Subtrees of the GTDB-Tk 2.1.1 phylogenomic tree showing the affiliation of *'Ca* Atrisphaera sulfidophila' Plnct-SY6 (shown in red color) with other closely related members of the phylum *Planctomycetota*. The length of the bar indicates 10 nucleotide substitutions per 100 nucleotides. Taxa name shown in blue color are also obtained from the Black Sea. CARD-FISH microscopic analysis of '*Ca* Atrimarinobacter sulfidophilus' strain Cloa-SY6 **(C)** DAPI **(panel C)** and CARD-FISH **(panel D)** microscopical analysis of (C and D) the enrichment culture of *'Ca* Atrisphaera sulfidophila' Plnct-SY6 using 0.02% chitin medium. All microbial cells were made visible by staining with DAPI, while cells of *'Ca* Atrisphaera sulfidophila' Plnct-SY6 were revealed by using the specifically designed fluorescent probe $PB1_{Alexa488}$ (see M&M for details).

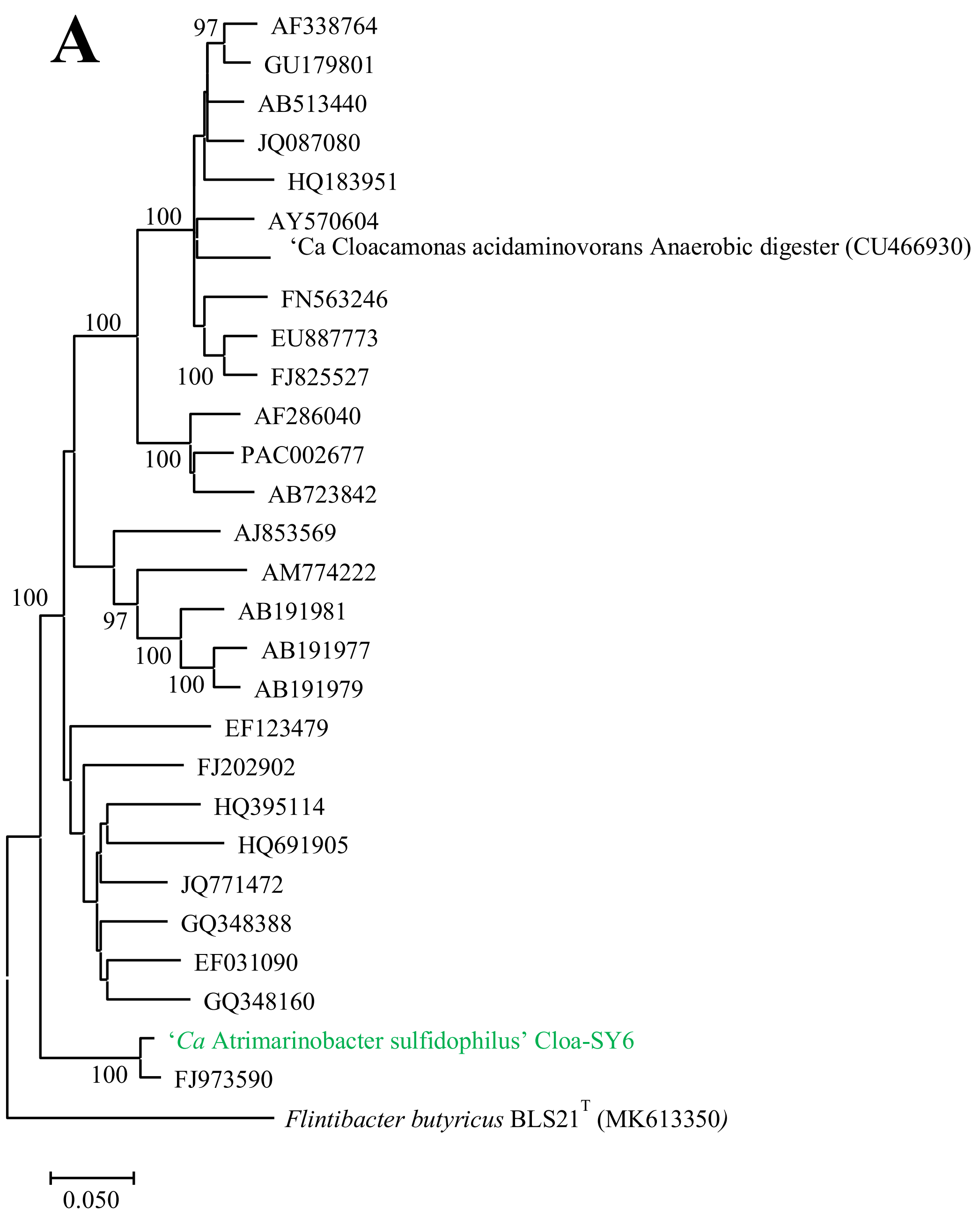


**Fig. S8: (A)** Phylogenetic tree based on 16S rRNA gene sequences showing the relationship of *'Ca* Atrimarinobacter sulfidophilus' Cloa-SY6 and other closely related members of the phylum *Cloacimonadota*. The tree was reconstructed by the maximum-likelihood method using MEGA X software and was rooted by using the 16S rRNA gene sequence of *Flintibacter butyricus* BLS21$^T$ (MK613350) as the outgroup. Numbers at nodes represent bootstrap value (percentages, based on 1000 resamplings). GenBank accession numbers for 16S rRNA gene sequences are shown in parentheses. Bar, 5 nucleotide substitutions per 100 nucleotides.

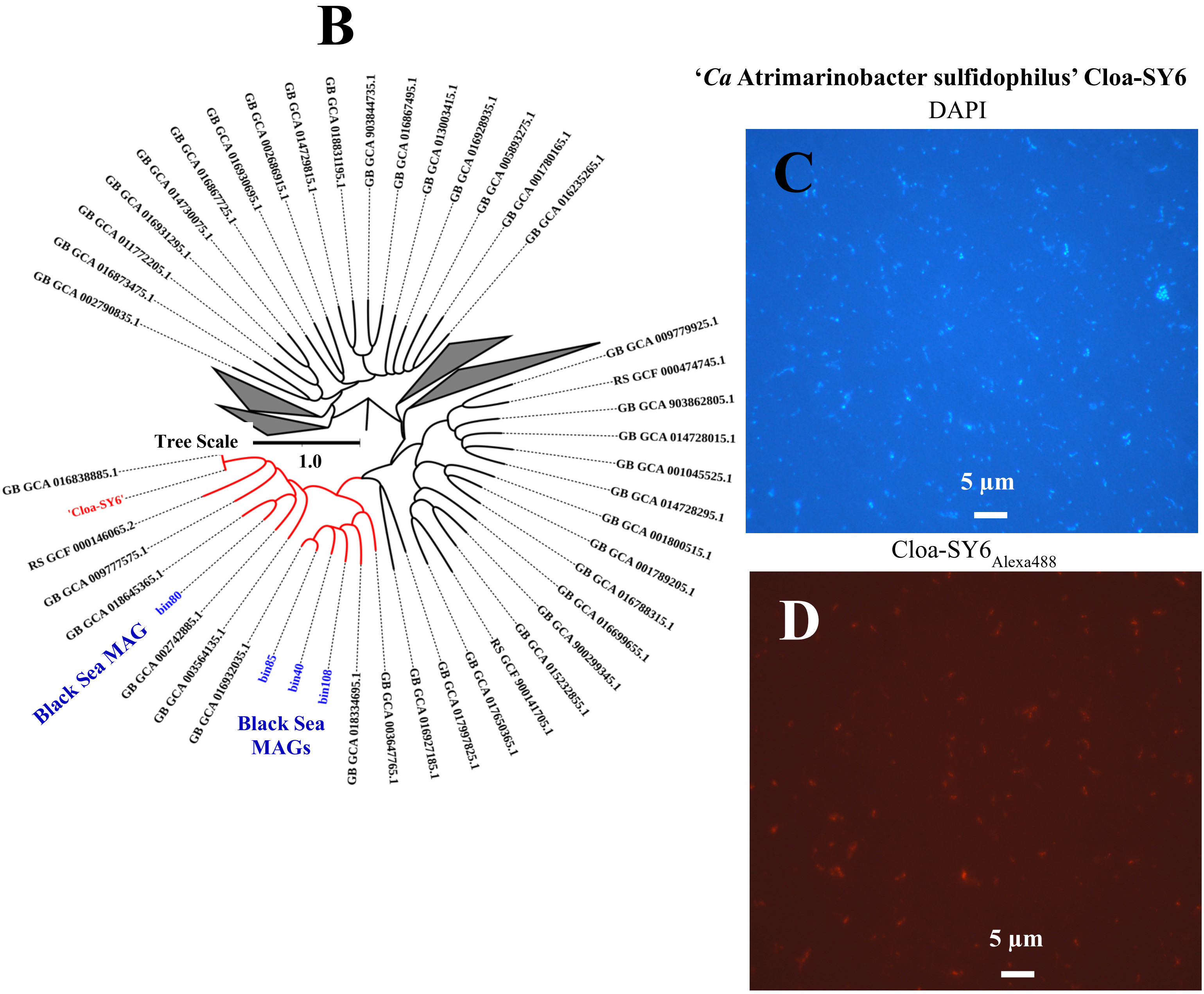


**Fig. S8: (B)** Subtrees of the GTDB-Tk 2.1.1 phylogenomic tree showing the affiliation of *'Ca* Atrimarinobacter sulfidophilus' Cloa-SY6 (shown in red color) within the closely related members of the phylum *Cloacimonadota*. MAGs shown in blue color obtained from the Black Sea waters. The length of the bar indicates 100 nucleotide substitutions per 100 nucleotides. CARD-FISH microscopic analysis of *'Ca* Atrimarinobacter sulfidophilus' Cloa-SY6 **(B)** DAPI **(panel B)** and CARD-FISH **(panel C)** microscopical analysis of (B and C) the enrichment culture of *'Ca* Atrimarinobacter sulfidophilus' Cloa-SY6 using 0.02% propionate medium. All microbial cells were made visible by staining with DAPI, while cells of *'Ca* Atrimarinobacter sulfidophilus' Cloa-SY6 were revealed by using the specifically designed fluorescent probe $PB1_{Alexa488}$ (see M&M for details).

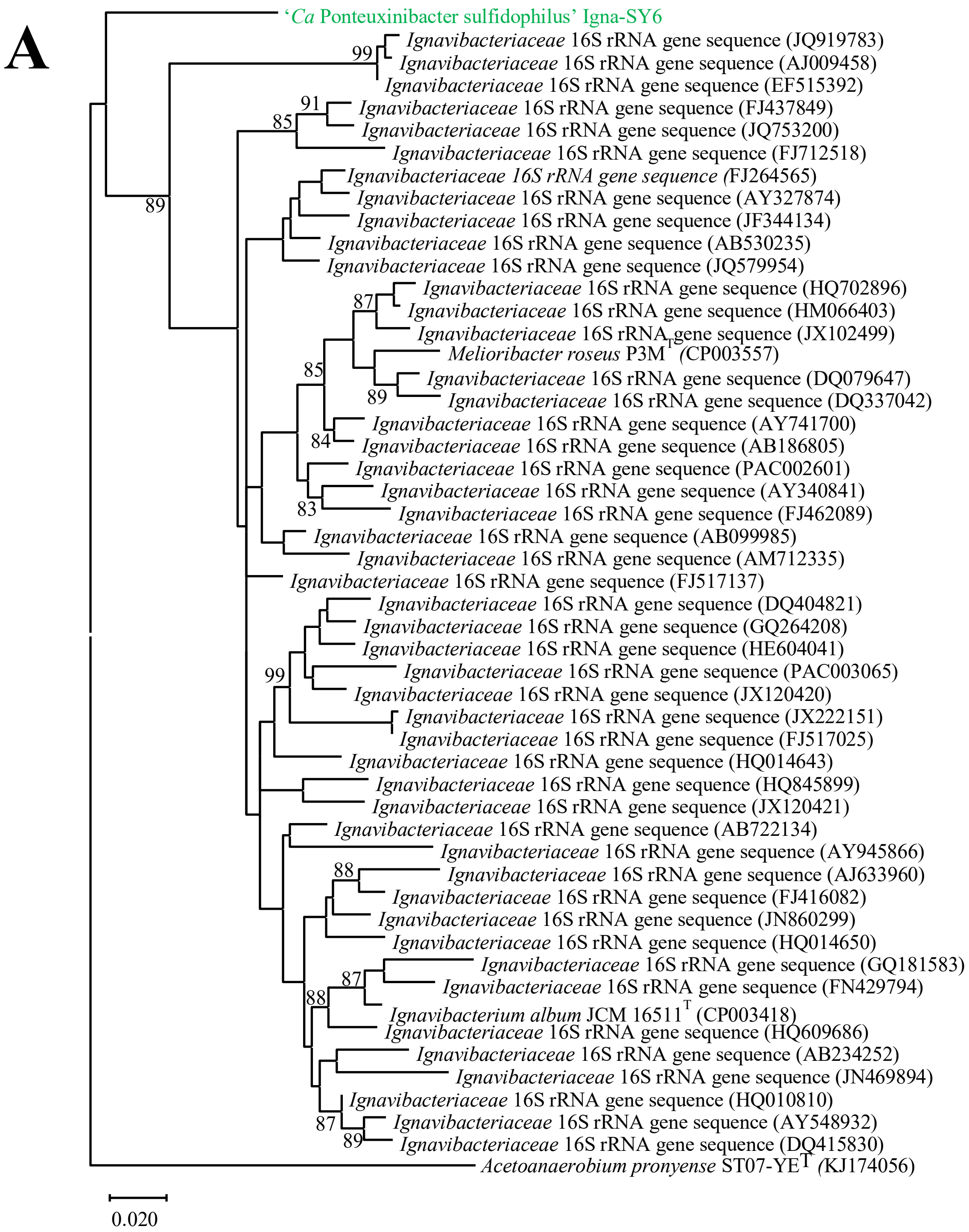


**Fig. S9: (A)** Phylogenetic tree based on 16S rRNA gene sequences showing the relationship of *'Ca* Ponteuxinibacter sulfidophilus' Igna-SY6 and other closely related members of the phylum *Ignavibacteriota*. The tree was reconstructed by the maximum-likelihood method using MEGA X software and was rooted by using *Phycisphaerae* 16S rRNA gene sequence (JN538045) as the outgroup. Numbers at nodes represent bootstrap value (percentages, based on 1000 resamplings). GenBank accession numbers for 16S rRNA gene sequences are shown in parentheses. Bar, 2 nucleotide substitutions per 100 nucleotides.

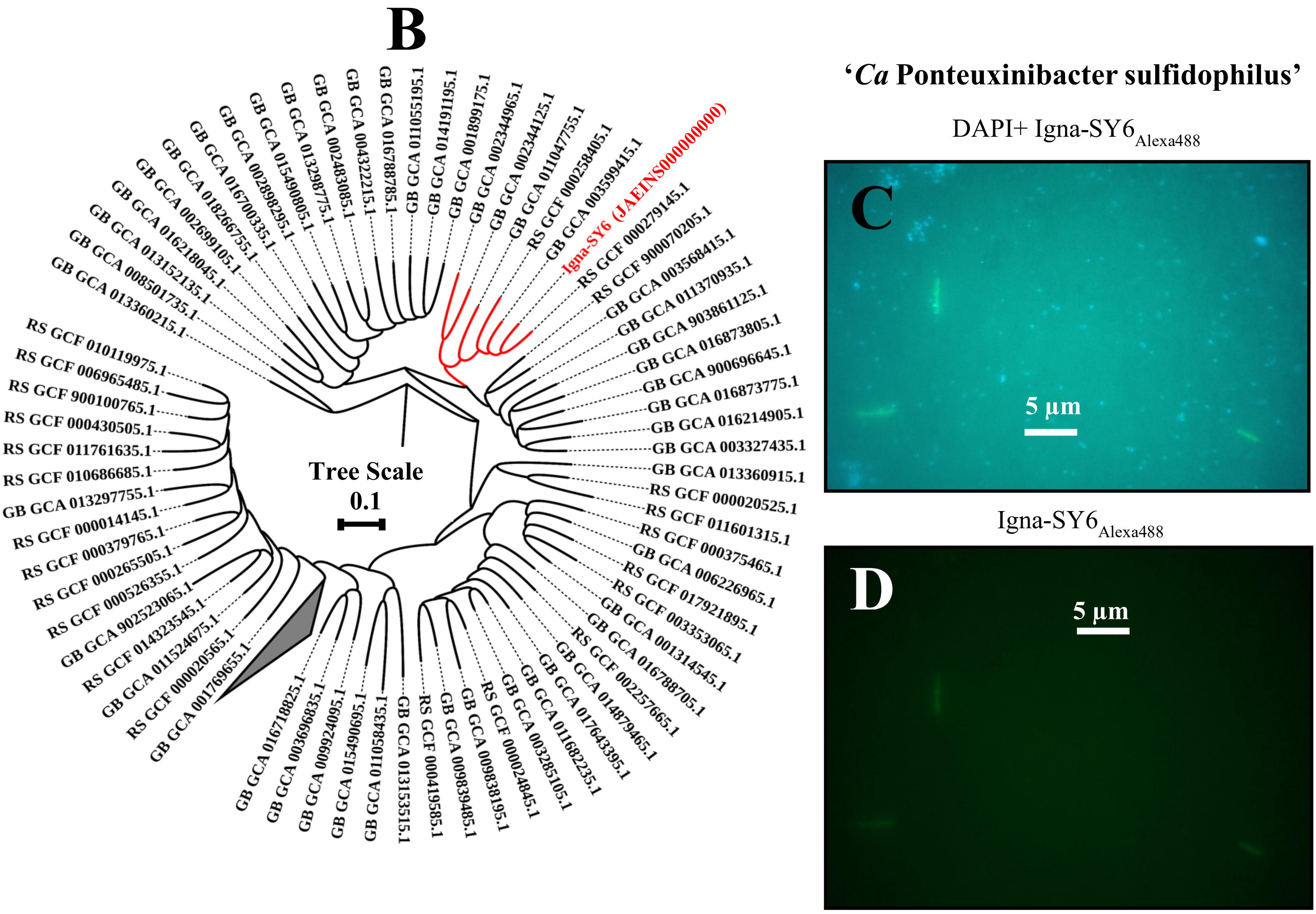


**Fig. S9: (B)** Subtrees of the GTDB-Tk 2.1.1 phylogenomic tree showing the affiliation of '*Ca* Ponteuxinibacter sulfidophilus' Igna-SY6 (shown in red color) within the closely related members of the phylum *Ignavibacteriota*. The length of the bar indicates ten nucleotide substitutions per 100 nucleotides. CARD-FISH microscopic analysis of '*Ca* Ponteuxinibacter sulfidophilus' strain Igna-SY6 **(B)** DAPI **(panel B)** and CARD-FISH **(panel C)** microscopical analysis of (B and C) the enrichment culture of '*Ca* Ponteuxinibacter sulfidophilus' strain Igna-SY6 using 0.02% cellulose medium. All microbial cells were made visible by staining with DAPI, while cells of '*Ca* Ponteuxinibacter sulfidophilus' strain Igna-SY6 were revealed by using the specifically designed fluorescent probe PB1$_{Alexa488}$ (see M&M for details).

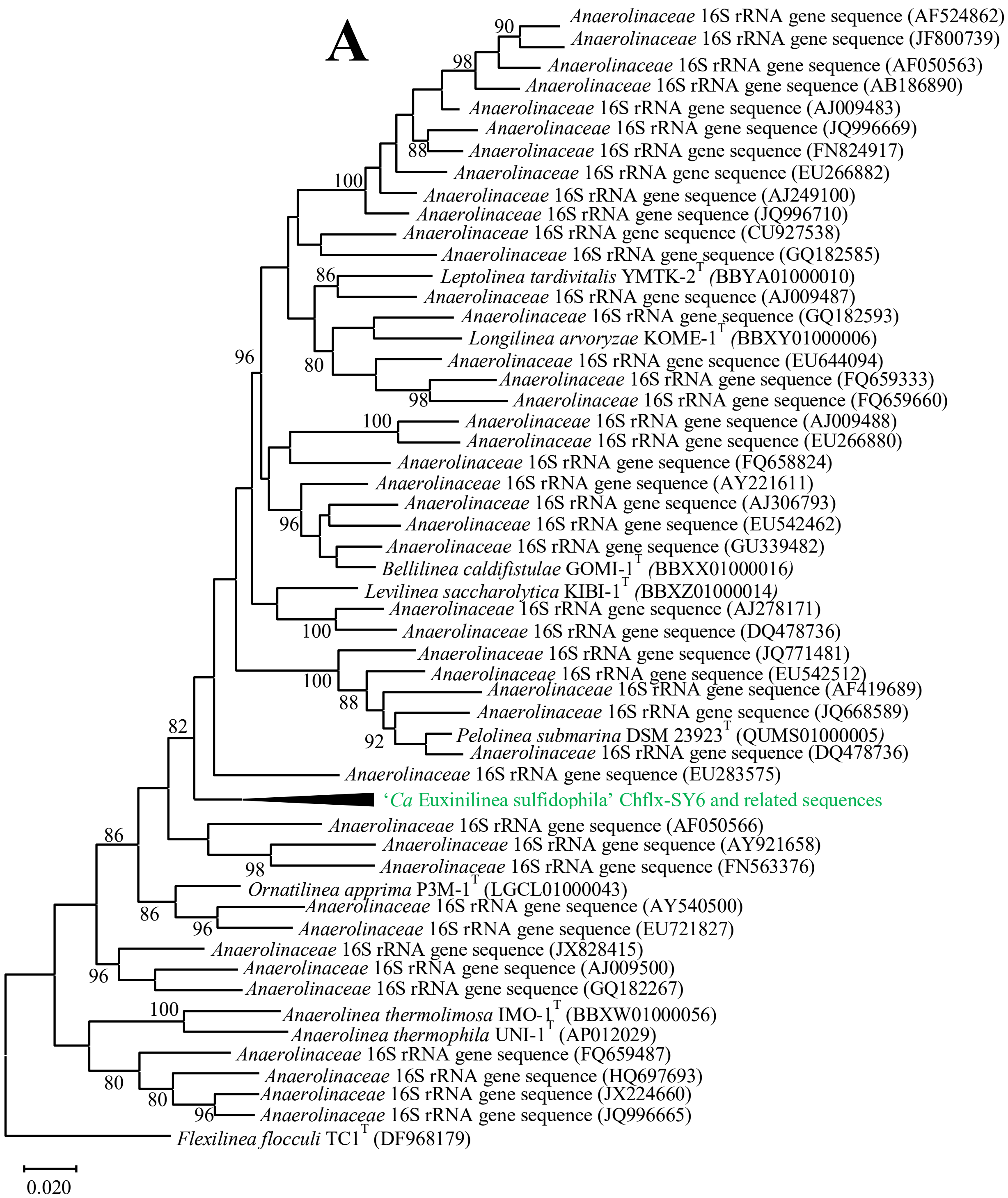


**Fig. S10: (A)** Phylogenetic tree based on 16S rRNA gene sequences showing the relationship of *'Ca* Euxinilinea sulfidophila' strain Chflx-SY6 and other closely related members of the phylum *Chloroflexota*. The tree was reconstructed by the maximum-likelihood method using MEGA X software and was rooted by using the 16S rRNA gene sequence of *Flexilinea flocculi* TC1$^T$ (DF968179) as the outgroup. Numbers at nodes represent bootstrap value (percentages, based on 1000 resamplings). GenBank accession numbers for 16S rRNA gene sequences are shown in parentheses. Bar, 2 nucleotide substitutions per 100 nucleotides.

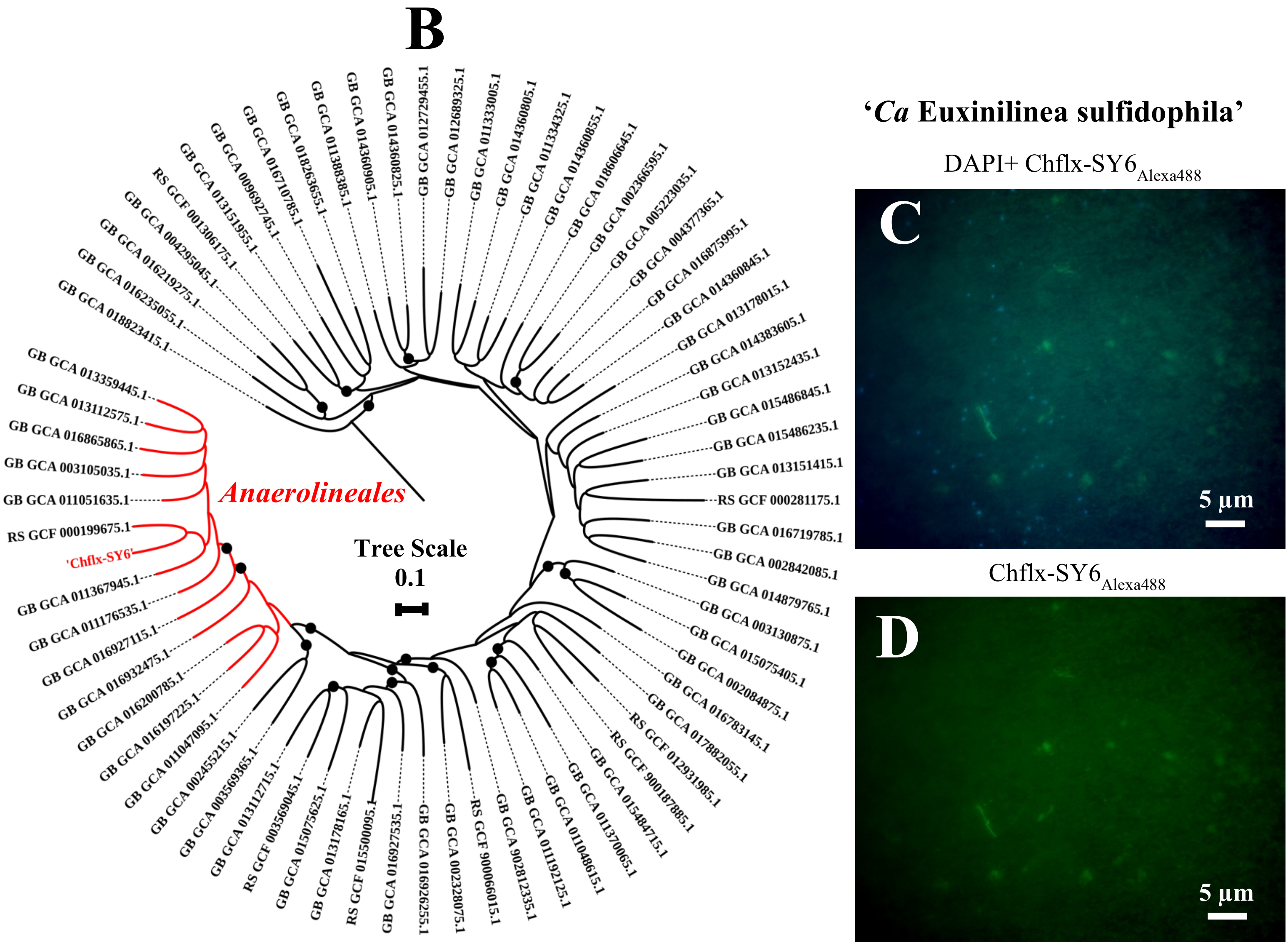


**Fig. S10: (B)** Subtrees of the GTDB-Tk 2.1.1 phylogenomic tree showing the affiliation of '*Ca* Euxinilinea sulfidophila' Chflx-SY6 (labeled in red color) within the closely related members of the phylum *Chloroflexota*. Black circle at the nodes represents the bootstrap value (100). The length of the bar indicates 10 nucleotide substitutions per 100 nucleotides. CARD-FISH microscopic analysis of '*Ca* Euxinilinea sulfidophila' strain Chflx-SY6 **(B)** DAPI **(panel B)** and CARD-FISH **(panel C)** microscopical analysis of (B and C) the enrichment culture of '*Ca* Euxinilinea sulfidophila' strain Chflx-SY6 using 0.02% cellulose medium. All microbial cells were made visible by staining with DAPI, while cells of '*Ca* Euxinilinea sulfidophila' strain Chflx-SY6 were revealed by using the specifically designed fluorescent probe PB1$_{Alexa488}$ (see M&M for details).

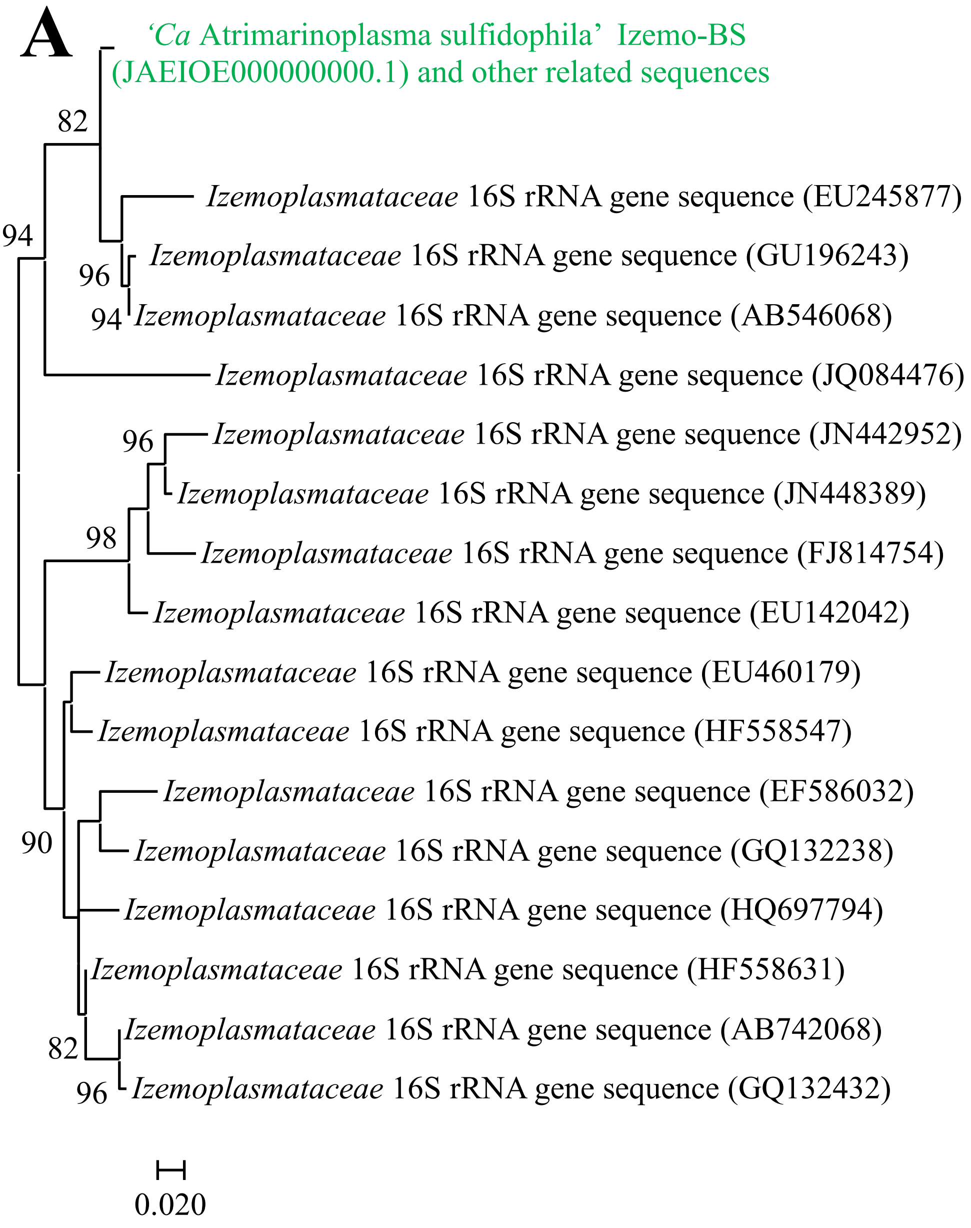


**Fig. S11: (A)** Phylogenetic tree based on 16S rRNA gene sequences showing the relationship of *'Ca* Atrimarinoplasma sulfidophila*'* Izemo-BS and other closely related members of the phylum *Mycoplasmatota*. The tree was reconstructed by the maximum-likelihood method using MEGA X software and was rooted by using *Izemoplasmataceae* 16S rRNA gene sequence (GQ132432) as the outgroup. Numbers at nodes represent bootstrap value (percentages, based on 1000 resamplings). GenBank accession numbers for 16S rRNA gene sequences are shown in parentheses. Bar, 2 nucleotide substitutions per 100 nucleotides.

B

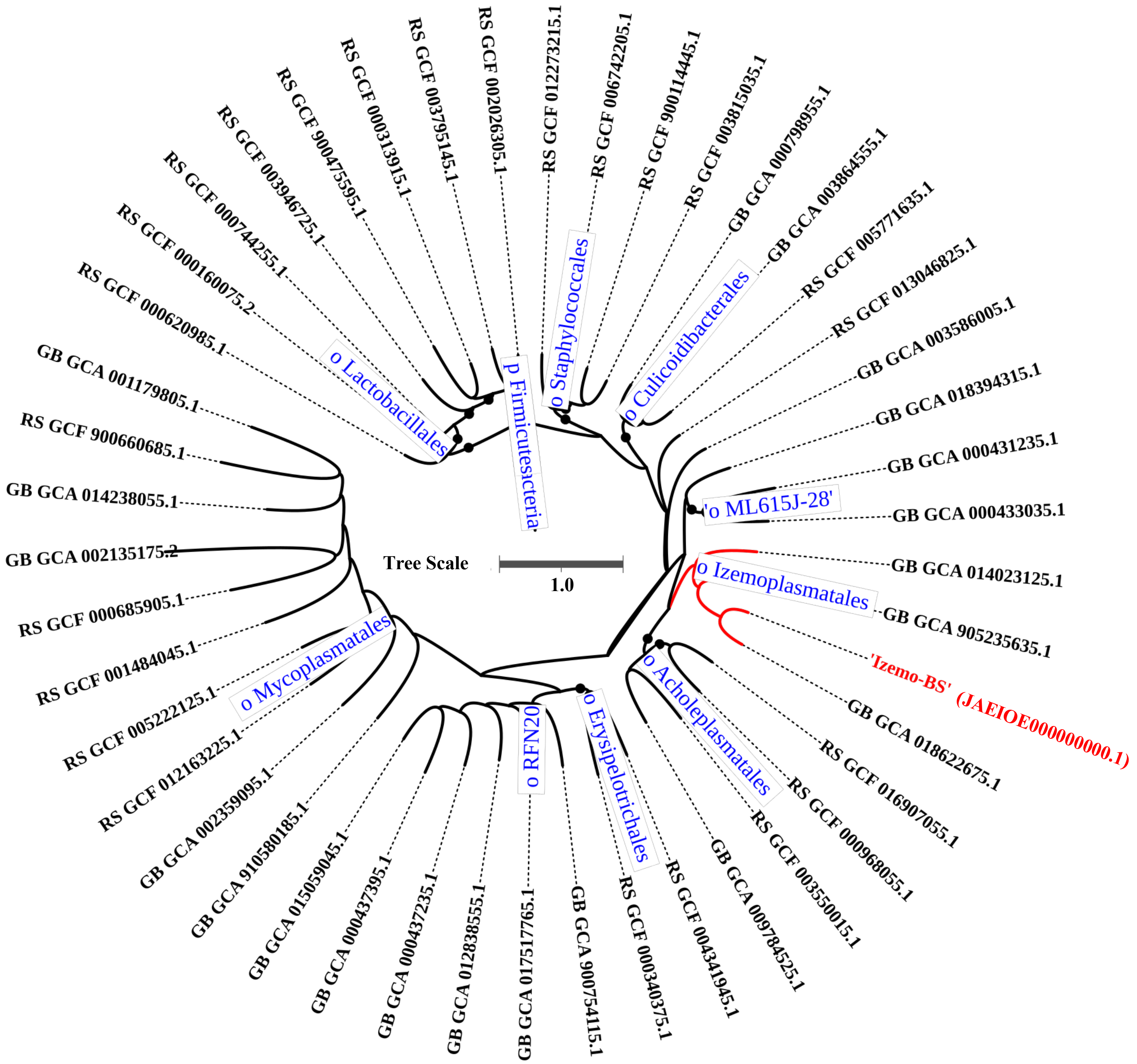


**Fig. S11: (B)** Subtrees of the GTDB-Tk 2.1.1 phylogenomic tree showing the affiliation of '*Ca* Atrimarinoplasma cellobiosiphila' Izemo-BS (shown in red color) with other closely related members of the phylum *Mycoplasmatota*. Black circle at the nodes represents the bootstrap value (100). The length of the bar indicates 100 nucleotide substitutions per 100 nucleotides.

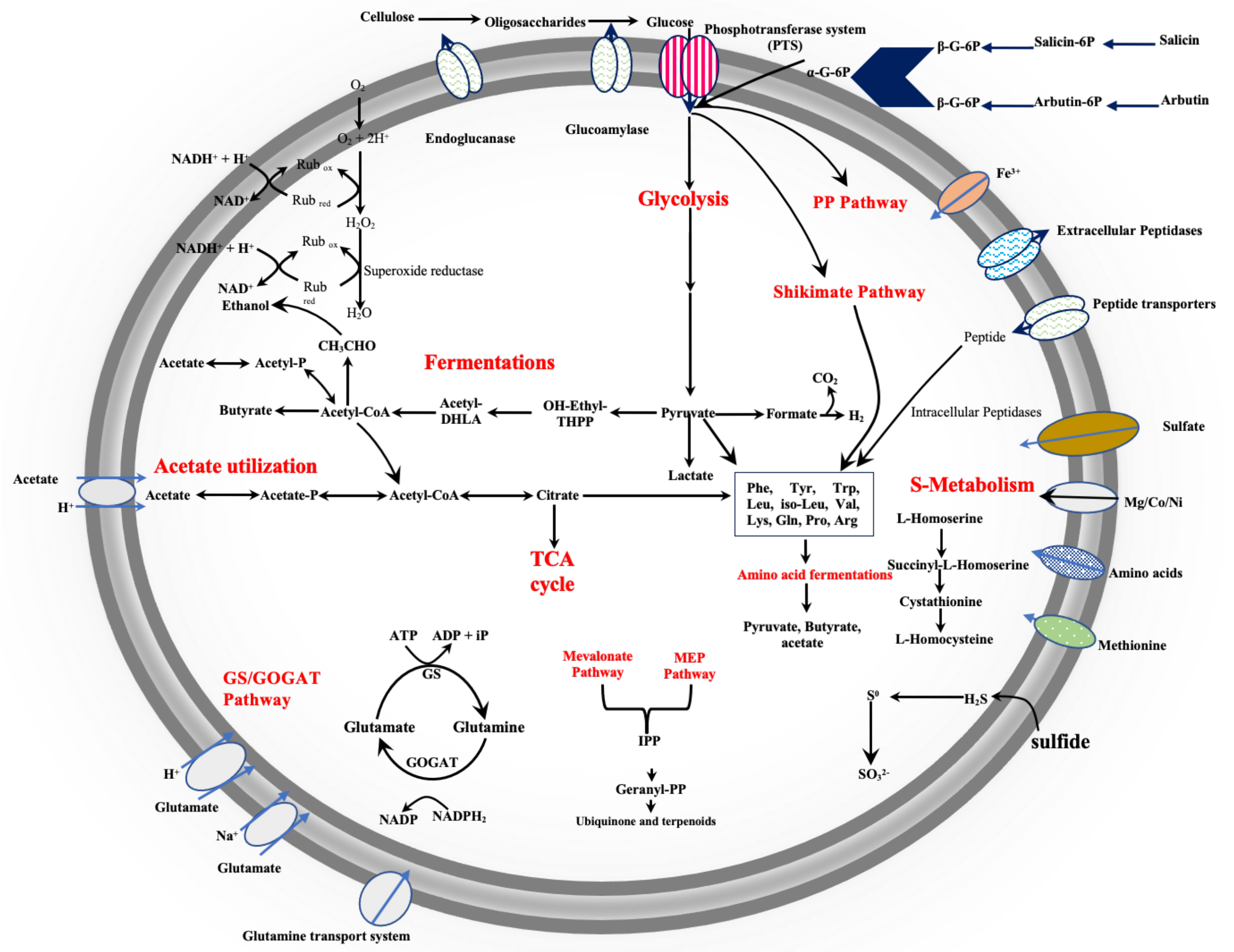


**Fig. S12**: Reconstruction of the central metabolic pathway of *Psychrilyobacter piezotolerans* strain S5 based on different physiological analyses and by the presence of various genes identified in the genome sequence. IPP, isopentenyl pyrophosphate; PP Pathway, pentose phosphate pathway; geranyl-PP, geranyl pyrophosphate; GS, glutamine synthetase; glutamate synthase (GOGAT).

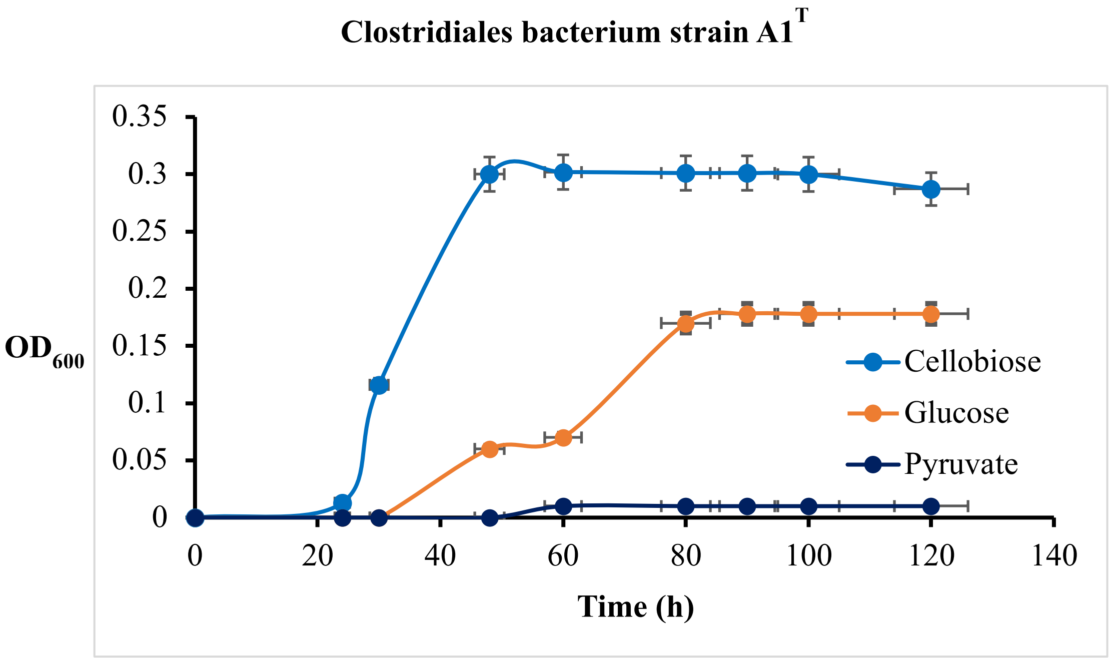


**Fig. S13:** Growth of *Clostridiales* bacterium strain A1$^{T}$ at various carbon substrate at 20 °C under strict anaerobic condition.

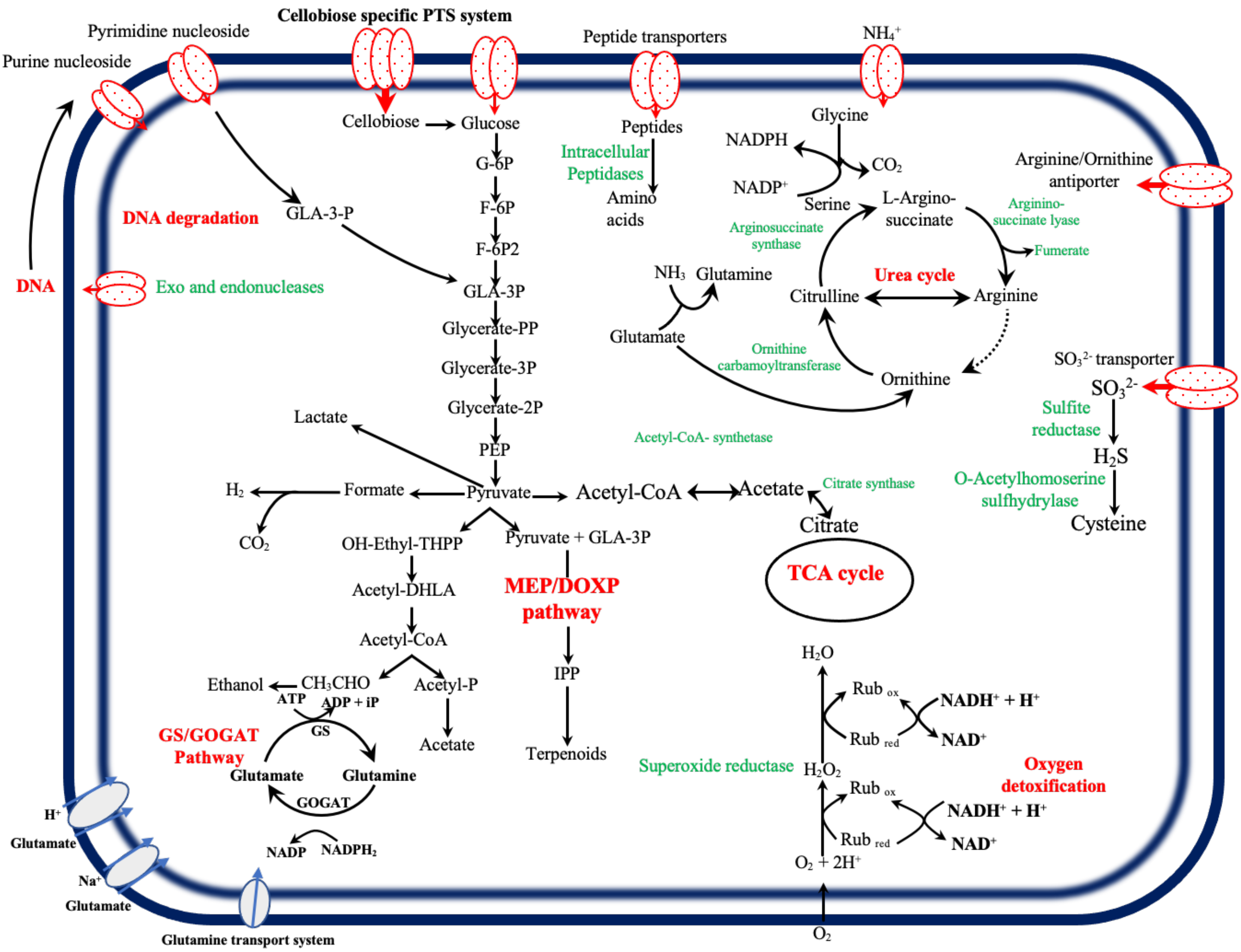


**Fig. S14**: Reconstruction of the central metabolic pathway of *Clostridiales* bacteria strains A1$^T$ and A2 based on different physiological analyses and by the presence of various genes identified in the genome sequence. IPP, isopentenyl pyrophosphate; G-6P, glucose-6-phosphate; F-6P, fructose-6-phosphate; GLA-3P, glyceraldehyde-3-phosphate; PEP, phosphoenol-pyruvate; GS, glutamine synthetase; glutamate synthase (GOGAT).

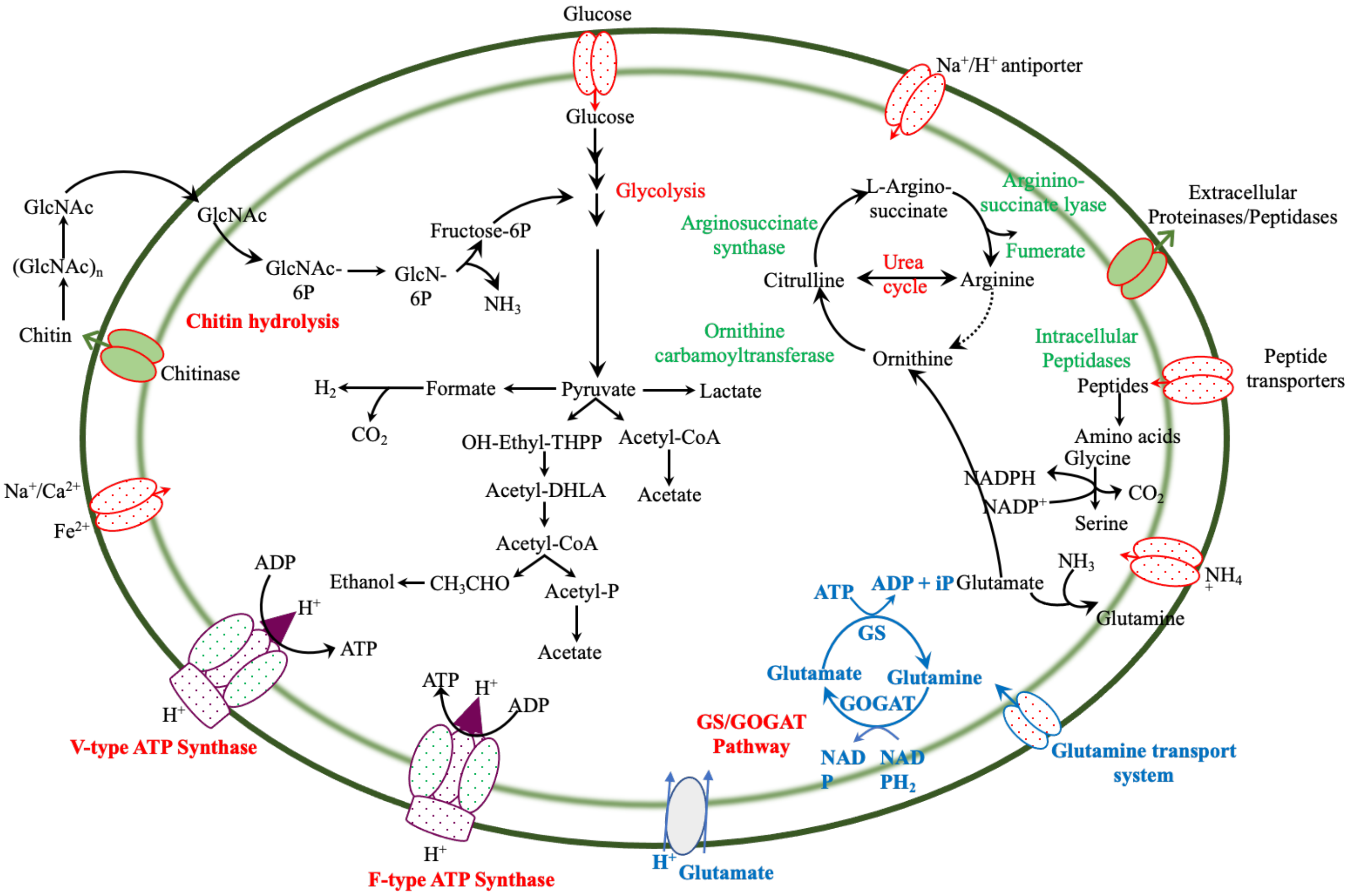


**Fig. S15**: Reconstruction of the central metabolic pathway of '*Ca* Atrisphaera chitinolytica' strain Plnct-SY6 based on different physiological analyses and by the presence of various genes identified in the MAG. GlcNAc, *N*-acetyl-D-glucosamine; GlcNAc-6P, *N*-acetyl-D-glucosamine-6-phosphate; GS, glutamine synthetase; glutamate synthase (GOGAT).

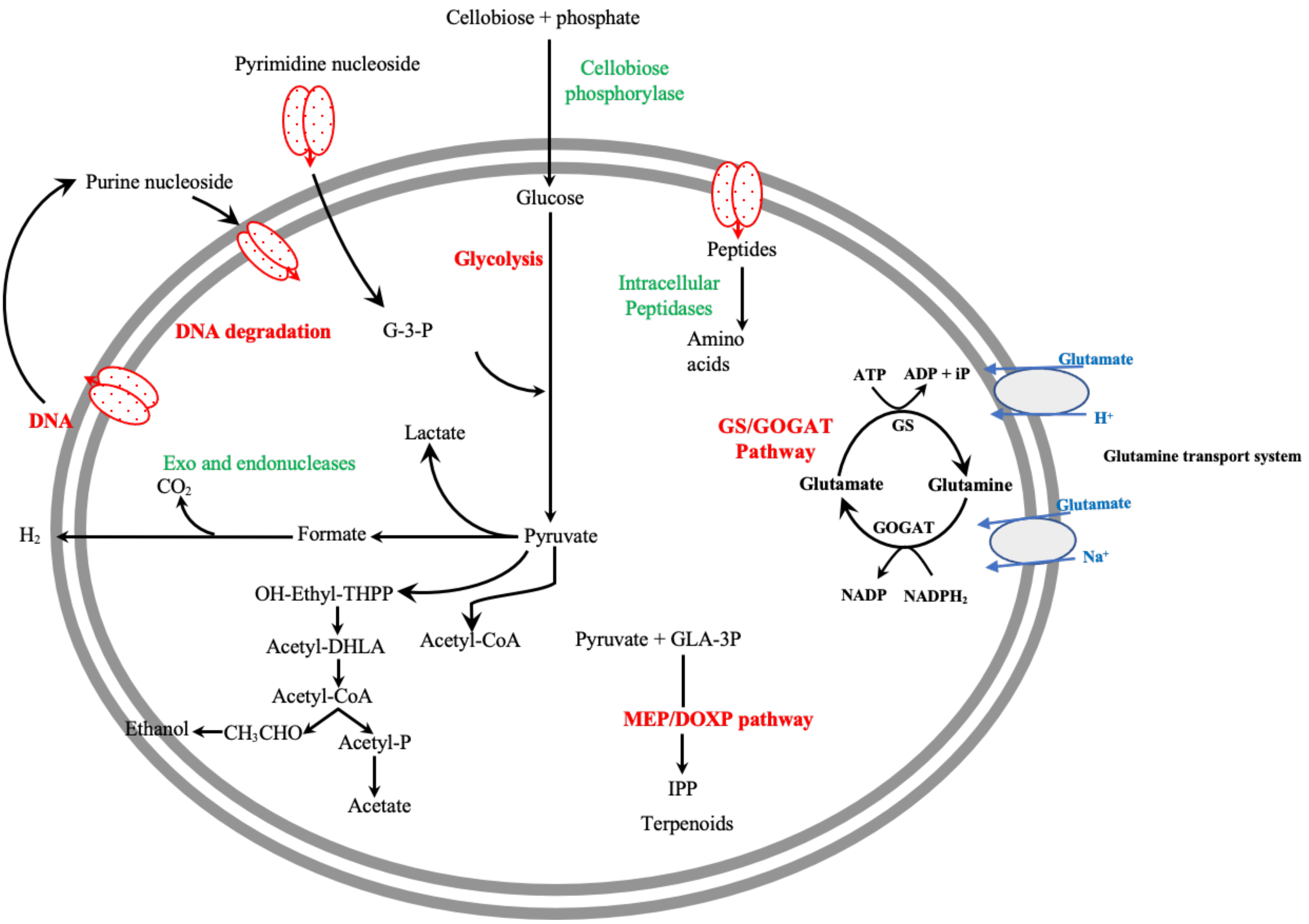


**Fig. S16**: Reconstruction of the central metabolic pathway of '*Ca* Atrimarinoplasma sulfidophila' Izemo-BS based on different physiological analyses and by the presence of various genes identified in the genome sequence. GS, glutamine synthetase; glutamate synthase (GOGAT).

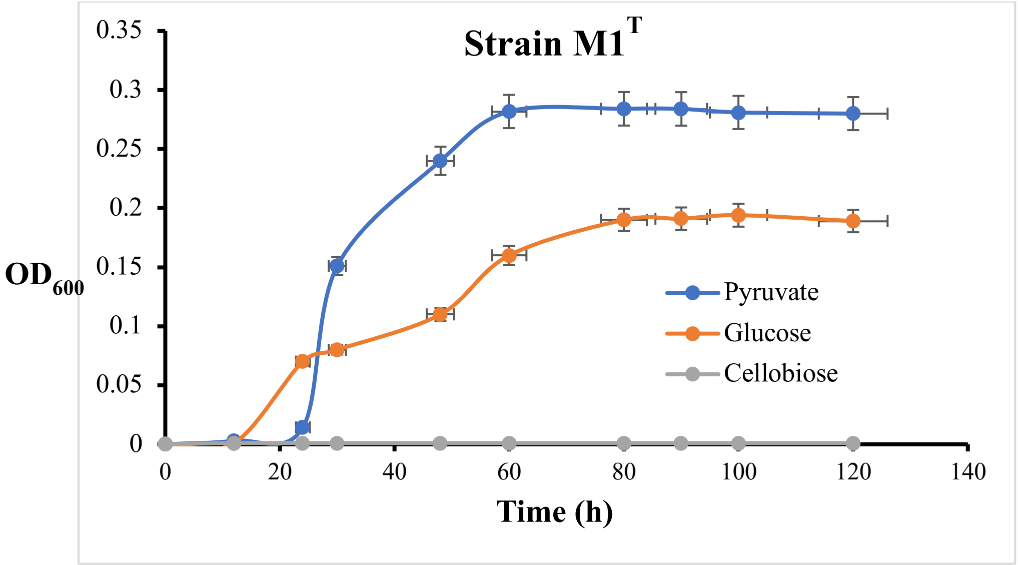


**Fig. S17**: Growth of *Oceanispirochaeta piezotolerans* strain M1$^T$ at various carbon substrate at 20 °C under strict anaerobic condition.

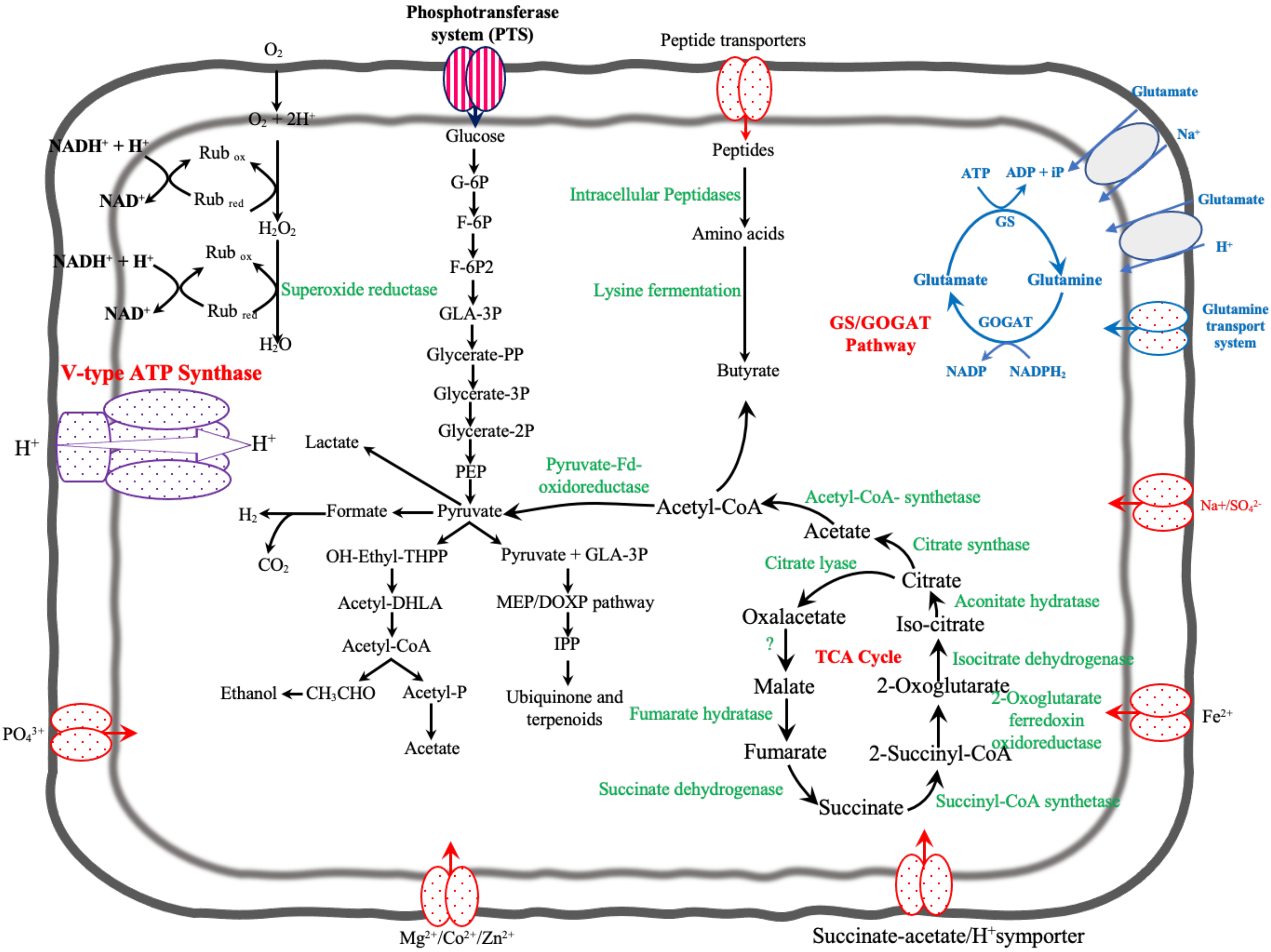


**Fig. S18**: Reconstruction of the central metabolic pathway of *Oceanispirochaeta piezotolerans* strain M1$^T$ and M2 based on different physiological analyses and by the presence of various genes identified in the genome sequence. G-6P, glucose-6-phosphate; F-6P, fructose-6-phosphate; GLA-3P, glyceraldehyde-3-phosphate; PEP, phosphoenol-pyruvate; GS, glutamine synthetase; glutamate synthase (GOGAT).

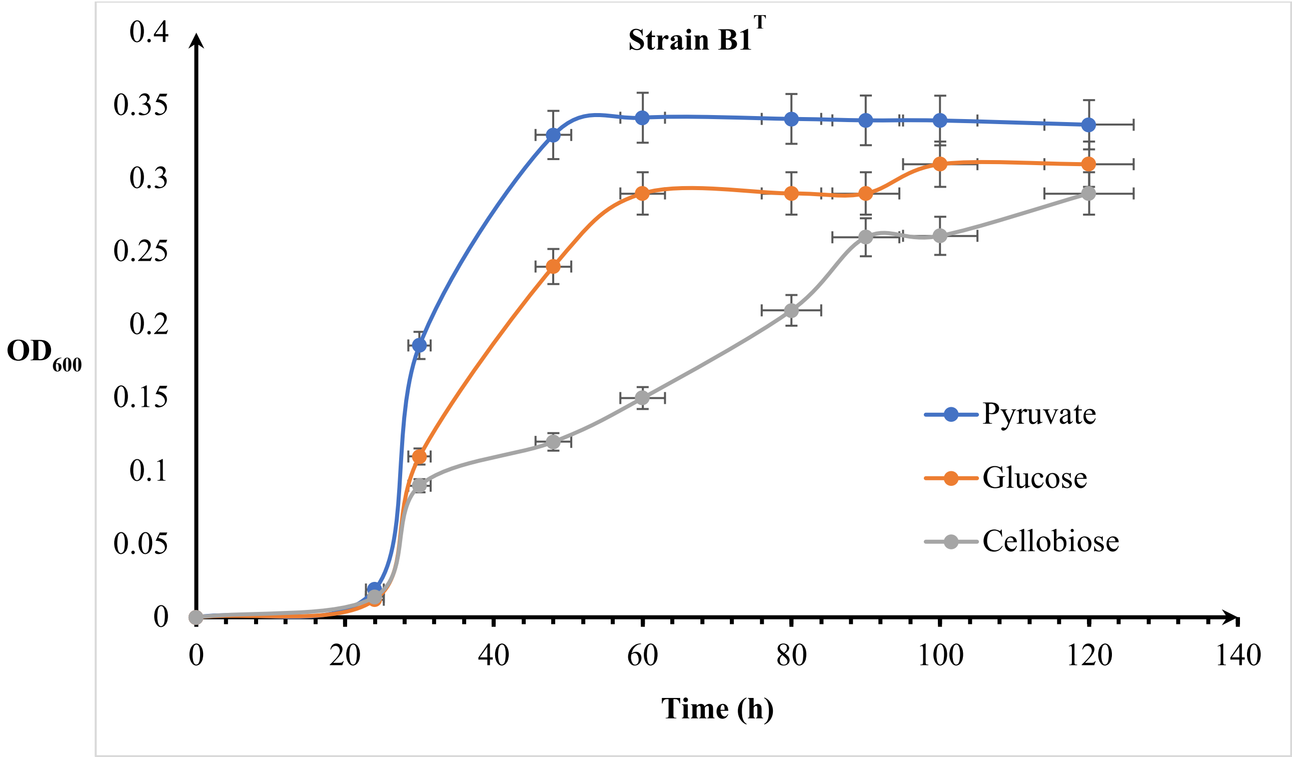


**Fig. S19**: Growth of *Lutibacter atrimaris* strain B1$^T$ at various carbon substrates at 20 °C under microaerophilic condition.

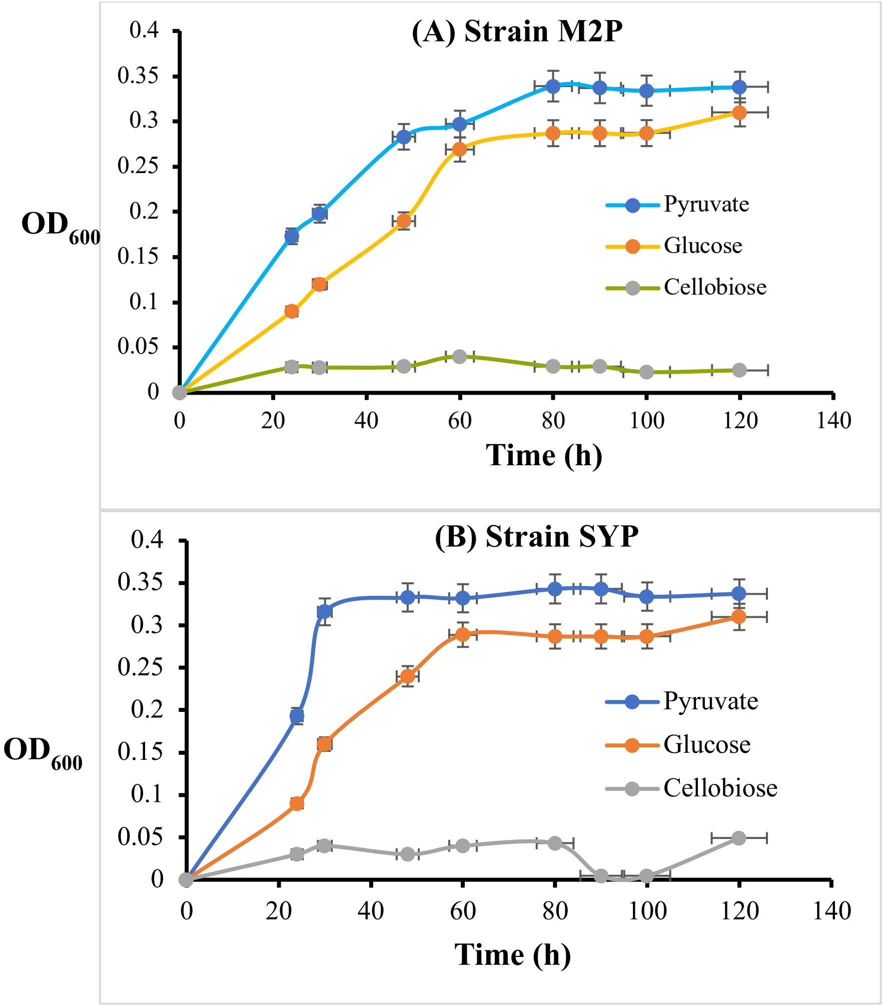


**Fig. S20**: **(A)** Growth of *Ancylomarina euxinus* strain M2P and **(B)** *Labilibaculum euxinus* strain SYP on various carbon substrates at 20 °C under strict anaerobic condition.

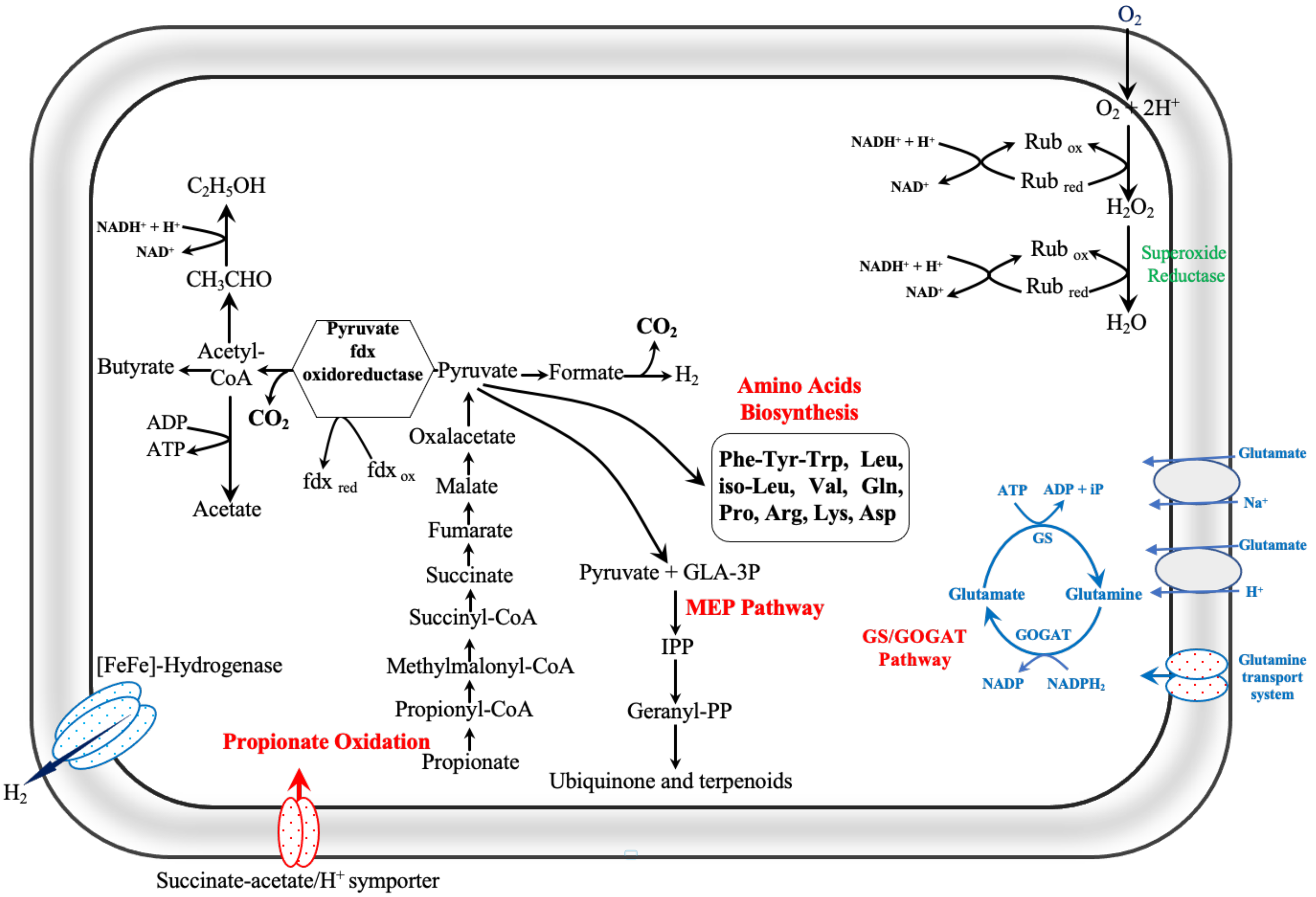


**Fig. S21**: Reconstruction of the central metabolic pathway of '*Ca* Atrimarinobacter sulfidophilus' Cloa-SY6 based on different physiological analyses and by the presence of various genes identified in the MAG. GS, glutamine synthetase; glutamate synthase (GOGAT).

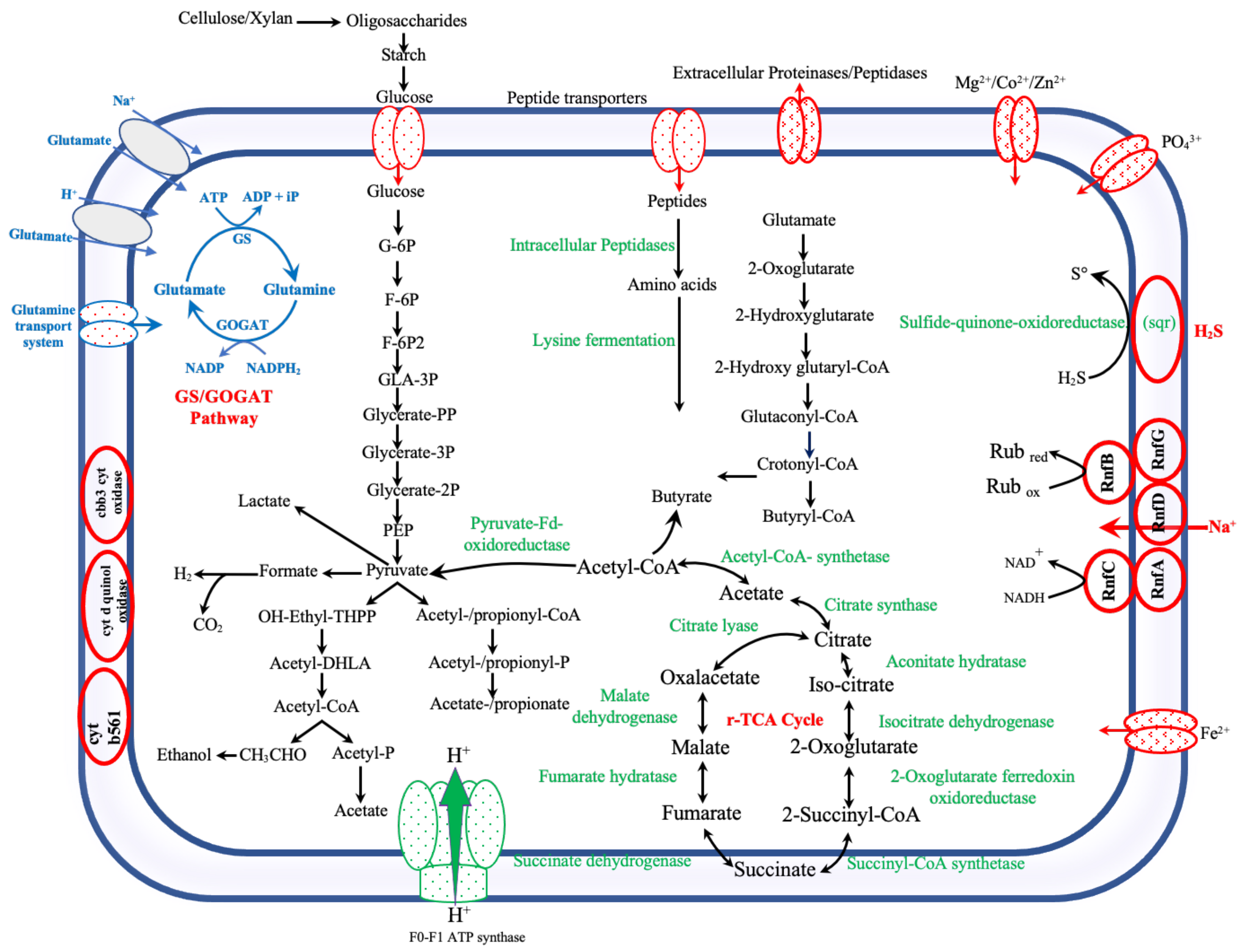


**Fig. S22**: Reconstruction of the central metabolic pathway of '*Ca* Ponteuxinibacter sulfidophilus' strain Igna-SY6 based on different physiological analyses and by the presence of various genes identified in the MAG. G-6P, glucose-6-phosphate; F-6P, fructose-6-phosphate; GLA-3P, glyceraldehyde-3-phosphate; PEP, phosphoenol-pyruvate; GS, glutamine synthetase; glutamate synthase (GOGAT).

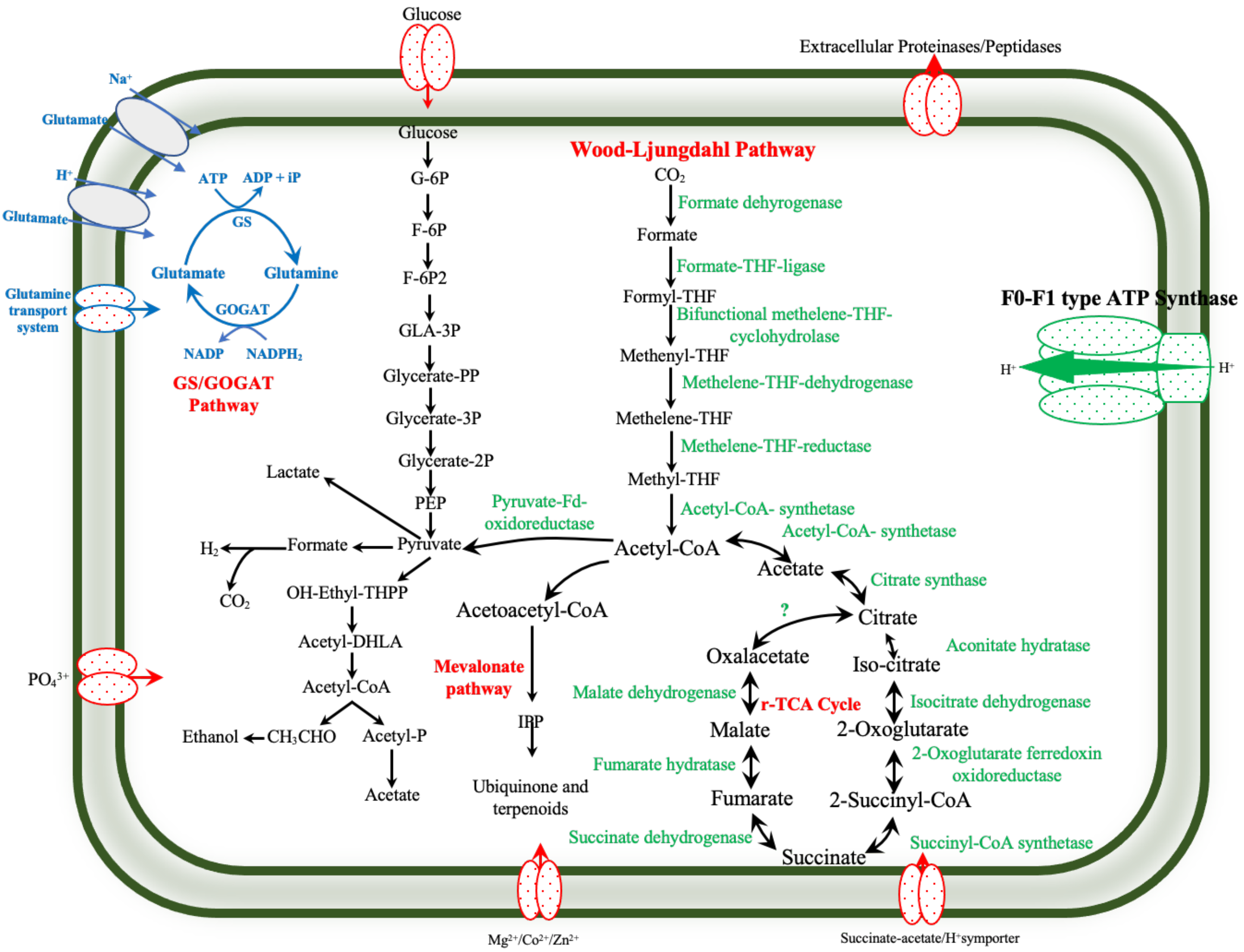


**Fig. S23**: Reconstruction of the central metabolic pathway of '*Ca* Euxinilinea sulfidophila' Chflx-SY6 based on different physiological analyses and by the presence of various genes identified in the MAG. G-6P, glucose-6-phosphate; F-6P, fructose-6-phosphate; GLA-3P, glyceraldehyde-3-phosphate; PEP, phosphoenol-pyruvate; GS, glutamine synthetase; glutamate synthase (GOGAT).

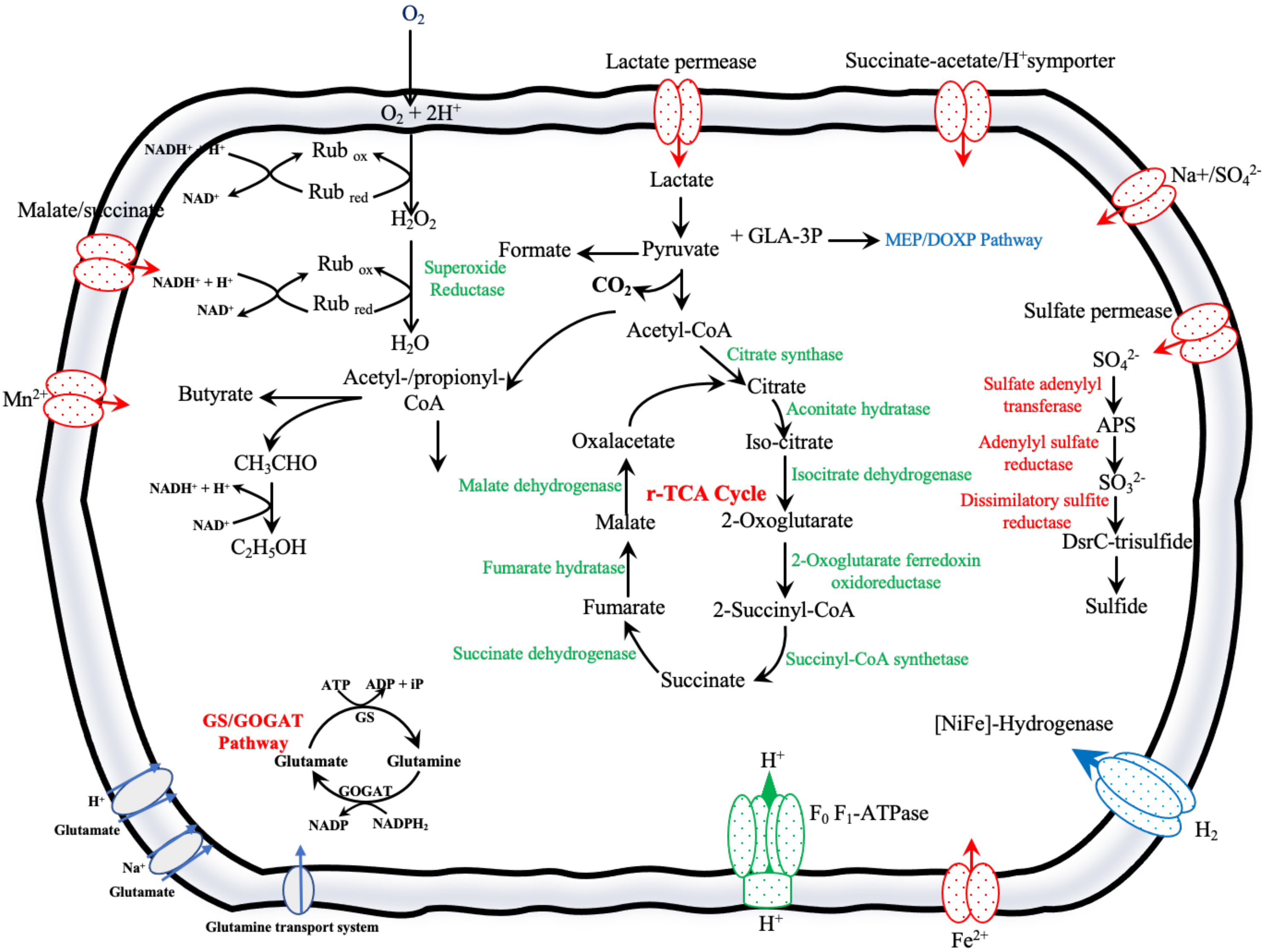


**Fig. S24**: Reconstruction of the central metabolic pathway of '*Pseudodesulfovibrio turroides* strain $S3^T$ and S3-i based on different physiological analyses and by the presence of various genes identified in the MAG. GLA-3P, glyceraldehyde-3-phosphate; GS, glutamine synthetase; glutamate synthase (GOGAT).

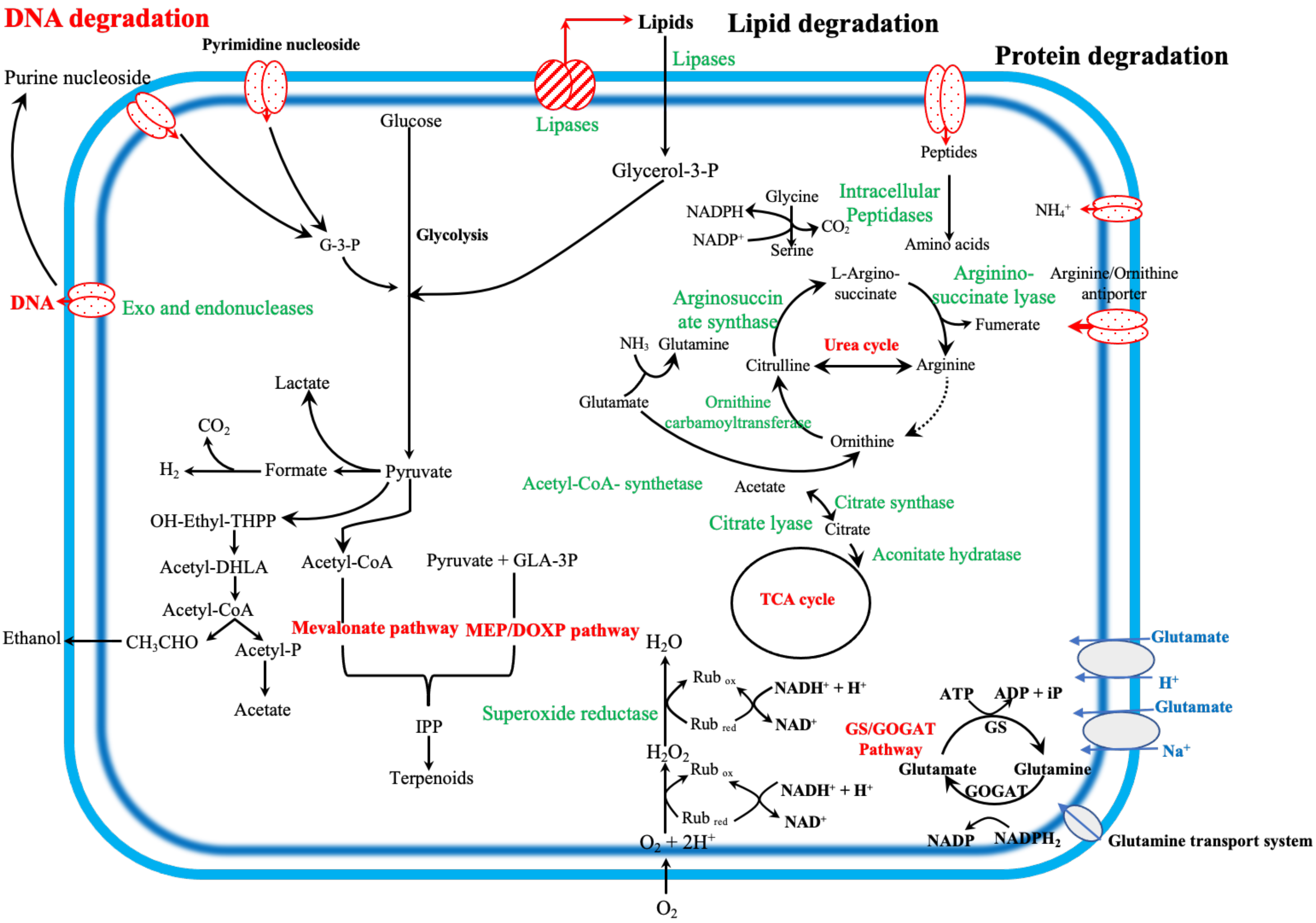


**Fig. S25**: Reconstruction of the central metabolic pathway of members of the phylum *Bacteroidota* (*Lutibacter atrimaris* strain B1$^T$, B2, '*Ca* Bradibacterium flavus strains S6 and L6; *Ancylomarina euxinus* strain M2P, M1P$^T$ and M3P; *Labilibaculum euxinus* strains SYP, A4$^T$ and 44) based on different physiological analyses and by the presence of various genes identified in the MAG. G-3-P, glycerol-3-phosphate; GLA-3P, glyceraldehyde-3-phosphate; GS, glutamine synthetase; glutamate synthase (GOGAT).
